# A platform-agnostic evaluation of non-formalin fixed single cell RNA technologies

**DOI:** 10.64898/2026.01.27.702057

**Authors:** Ellora Haukenfrers, Vaibhav Jain, Stephanie Arvai, Khooshbu Patel, Simon G. Gregory, Karen Abramson, Devi Swain-Lenz

## Abstract

The rapidly advancing field of single cell RNA sequencing (scRNAseq) offers numerous options for transcriptome profiling. However, questions remain as to which chemistry is appropriate for individual experimental goals. Preceding single cell benchmarking studies included previously available methods and involved a mixture of fresh and fixed samples or probe- and non-probe-based capture methods. However, the inherent differences in sample types and methods limited the conclusions to be drawn between analogous technologies. Here, we present a novel, systematic comparison of four widely used non-probe-based, non-formalin *fixed* scRNAseq assays. We build upon past comparisons that used varied computational pipelines by applying both platform-specific and agnostic cell calling algorithms for an unbiased comparison of biological and technical replicates from healthy human PBMCs. Our approach evaluates 10x Genomics, Parse Biosciences (QIAGEN), Scale Biosciences (10x Genomics), and Illumina scRNAseq assays to examine data based on accuracy, sensitivity, precision, power, and efficiency using agnostic and platform-specific cell calling. While metrics vary between assays, there are clear advantages and limitations to each technology, including experimental time and financial costs. In summary, our study highlights the need for carefully considered project design of non-formalin fixed scRNAseq assays, which is determined by many factors and dependent on an investigator’s specific research aims and available resources.

**Graphical Abstract:** 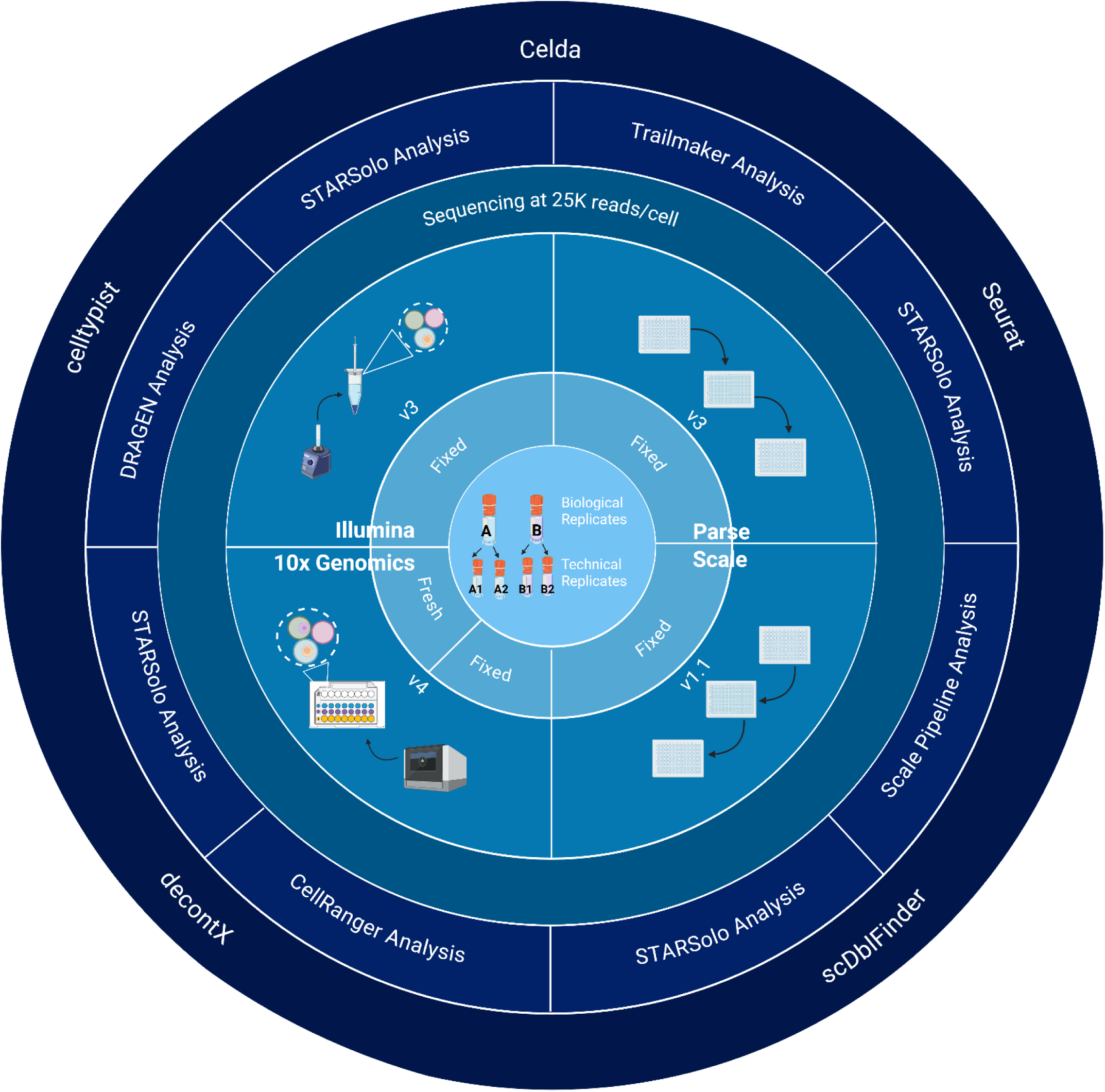

## BACKGROUND

Single cell transcriptome profiling methods are fundamental approaches in biological and medical research. Single cell RNA-sequencing (scRNAseq) measures RNA transcription at single cell resolution, enabling the identification of rare cell subtypes and examination of homeostatic developmental pathways, cell trajectories, gene regulatory networks, and signaling networks in heterogenous samples.^1,2^ As a result, scRNAseq has contributed to advances in oncology,^3–6^ immunology,^4–9^ and developmental biology,^4–6,8^ among other fields, by providing insight into disease mechanisms for improved prognosis and clinical intervention.^4,6,7,10–13^ Single cell technologies have evolved to the point that many commercial products are available at any given time, but each presents with their own strengths and weaknesses.^4,14,15^ In this study, we adopt the scRNAseq benchmark framework proposed by Ziegenhain et al., in which we evaluate platforms based on accuracy, sensitivity, precision, power, and efficiency (Table 1).^16^ For the sake of cost and practicality, we limit our end-to-end platform-specific and agnostic evaluations of the four most prevalent scRNAseq technologies by characterizing biological and technical replicates of human peripheral blood mononuclear cells (PBMCs). Finally, to control for variation between sample preparation and technologies, we limit our evaluation to the most recent non-probe-based, non-formalin fixed scRNAseq assays.

**Table 1.** Metrics Definitions Definitions of assessment metrics and method of how each metric is evaluated in this paper.

| ASSESSMENT METRIC DEFINITIONS |  |  |
| --- | --- | --- |
| Metric | Definition | What We Will Assess |
| Accuracy | The agreement between the expected proportion of mRNAs and the readouts for absolute gene expression in cells | Compare observed cell type proportions with expected proportions in healthy PBMCs, reproducibility analysis |
| Sensitivity | The likelihood of representing a particular mRNA transcript as a cDNA molecule in the final library | Number of detected genes per cell, total number of genes detected across cells, even coverage along mRNA transcripts, proportion of low abundance cell types |
| Precision | Consistency of gene expression measurements | With biological replicates, subsample to analyze dropout probability and amplification noise (coefficient of variation), examining technical variation |
| Power | Impact of sensitivity, precision, and number of cells on ability to detect differential expression levels | True positive rate (TPR) and false discovery rate (FDR), to demonstrate variation between technical replicates due to batch effects and/or sequencing depth |
| Efficiency | Costs required to reach a desired power | Throughput, RNA capture efficiency, cDNA amplification efficiency, efficiency of sequencing depth, ease-of-use, multiplet rate |

### Chemistries of Evaluated scRNAseq Platforms

The most mature technology we evaluate is the droplet-based poly-A-capture approach using 10x Genomics microfluidic ‘Chromium’ instrument, released in 2016. The patented droplet methodology captures single cells in barcoded Gel-bead-in-emulsions (GEMs), utilizing 3’-capture-based whole transcriptome profiling, with a purported cell capture efficiency up to 80% in their new GEM-X (v4) chemistry.^17^ The platform’s use is widespread because of its time in the market, high throughput and quality, versatility, and user-friendly protocol.^2^ However, the platform requires an upfront equipment purchase and ongoing maintenance agreements, limiting access for some labs.

Recognizing the need for lower cost options with increased scalability, three companies released instrument-free methods in recent years. Rosenberg and Roco developed the plate-based Split Pool Ligation-based Transcriptome sequencing (SPLiT-seq) approach for fixed cells or nuclei, commercialized as the Parse Biosciences ‘Evercode Whole Transcriptome kit’ in 2021.^8,18^ Parse (acquired by QIAGEN in 2025) subsequently introduced random hexamers to the v3 Evercode assay in addition to the oligo(T) primed 3’ capture that enables elevated representation of the gene body and a cell capture efficiency around 28%.^14,19,20^ Similar to Parse, with an expected cell capture rate around 28%, Scale Biosciences (acquired by 10x Genomics in 2025) released a plate-based combinatorial indexing ‘RNA Single Cell Sequencing Kit’ with 3’ poly-A-capture on fixed cells or nuclei in 2022.^21^ Also in 2022, Fluent Biosciences (acquired by Illumina in 2024) released the Protein Interaction Profile Sequencing (PIP-Seq) method, with a purported 80% cell capture using an instrument-free approach for 3’ poly-A-capture within templated emulsions.^22^ Illumina and 10x uniquely allow for hundreds of cells or nuclei as input (fresh or fixed) while Parse, and Scale require thousands.

Here, we discuss important scRNAseq considerations prior to data generation and provide a head-to-head comparison of the aforementioned platforms. We assess the impact of oligo (dT) 3’ priming and hexamer priming strategies on gene body coverage and metrics such as accuracy, sensitivity, precision, power, and efficiency to enable researchers to make informed decisions regarding study design, including cost implications.

### Common Challenges in scRNAseq

Regardless of the platform used, scRNAseq methods share fundamental challenges that add non-biological variation to experiments.^2,23–25^ The largest source of variation is the preparation of single cell suspensions (cryopreservation, thawing, fixation, etc.), which can disrupt cellular activity, viability and mRNA levels.^4,8,13^ To mitigate some of these technical issues, we control for variation by using consistent cell thawing methods in this study. Other technical factors, such as the efficiency of mRNA capture, reverse transcription (RT), cDNA amplification, and sequencing depth can contribute to the missing gene expression data (i.e. dropout rate), which we examine in this study.^26^ Additionally, mitochondrial RNA (mtRNA) content, though a biologically relevant marker of cell stress or activity, can impact the efficiency of mRNA conversion to readable cDNA.^15^mtRNA is present in higher proportions in fresh compared to fixed samples due to disproportionate loss of cytosolic transcripts and decreased cell stress from fixation.^14,27,28^ The loss of cytosolic transcripts during fixation is thought to be caused by the enrichment of nuclear pre-mRNA, leading to an increased proportion of intronic reads in fixed samples.^27^Combinatorial barcoding methods detect decreased mtRNA content because of the many wash steps following sample preparation.^28,29^

Next-generation library preparation introduces technical variation inherent to most sequencing technologies today. The low concentration of RNA in single cells (∼1pg of RNA per PBMC) necessitates many PCR amplification cycles to reach minimum sequencing thresholds, which introduces amplification bias at the 3’ end and downstream amplification noise.^2,4,27^ Additionally, the molecular sampling of scRNAs is stochastic, which leads to overdispersion of the data, where the variance increases with the mean. To model expression variance, Poisson noise represents expected biological variation while extra-Poisson variability represents technical imprecision.^30^ In this study, we examine the extra-Poisson variability to understand the impact of 3’ amplification bias and downstream amplification on gene expression measurements.

Multiplets create another common challenge in scRNA-seq. Multiplet refers to the phenomenon when two or more cells are mistakenly assigned the same barcode and assessed as a single sample.^8^ A positive correlation has previously been observed between increasing multiplet rates and cell loading into droplet-based instruments.^24,28,31^ In combinatorial barcoding methods, a high doublet rate indicates cell aggregation: a known risk with fixed samples, which are often sticky.^28,32^ Here, we assess the variability due to multiplet rate using a consensus-based approach between two doublet detection tools.

Transcriptome data is typically aligned to a reference genome and quality checked (QC’ed) using company-specific proprietary pipelines. However, unknown differences in each pipeline impede accurate comparisons between technologies.^13^ In terms of basic analysis, De Simone et al. reported varied cell recovery in Illumina’s data processing pipelines, highlighting the need for agnostic cell calling in true unbiased comparison of both assay and analysis tool performance.^27^ To address this issue, we apply an agnostic approach to cell calling, contrasting results with platform-specific pipelines for alignment, QC, and count analysis (basic analysis).

### 10x Genomics Flex vs. Fixed 3’ GEM-X in Benchmarking Studies

Previous scRNAseq benchmarking studies have assessed the performance of 10x Genomics Flex probe-based assays of human and mouse protein-coding genes in fixed cells,^27–29,33^ while others have compared 10x Genomics *fresh* 3’ or 5’ NextGEM or GEM-X assays to those from *fixed* samples in Parse and Scale.^34^ However, this study represents an evaluation of broadly applicable, species-agnostic 3’ oligo(dT) primed transcriptome profiling using 10x Genomics 3’ GEM-X, Illumina, Parse, and Scale assays of poly-adenylated RNAs (Illumina, Scale) and non-coding RNA (Parse WT v3, 10x 3’ GEM-X).^14,17,20^ ^28^ Our study is novel for its use of platform-specific and agnostic analyses of gene body coverage and assay performance. It also represents the first evaluation of 10x Genomics new ‘Cell Fixation Protocol for GEM-X Single Cell 3’ & 5’ Assays’ with 3’ GEM-X (non-probe-based) chemistry and other non-formalin fixed methods including Illumina, Parse, and Scale.

## RESULTS

### Library Construction and Sequencing

To compare different single cell transcriptome kits, we constructed libraries from each platform across biological and technical replicates (Suppl Fig 1, Suppl Table 1). Briefly, commercially available vials of cryopreserved PBMCs (STEMCELL Technologies) from 2 donors (‘A’ and ‘B’) were run in duplicate across each fixed workflow (10x Genomics Chromium GEM-X Single Cell 3’ Reagent Kit v4, Illumina PIPseq V T10 3ʹ Single Cell RNA Kit, Parse Biosciences Evercode WT v3, and Scale Biosciences Single Cell RNA Sequencing Kit v1.1), providing technical replicates (A1/A2 and B1/B2). We designated the different donor’s cells as biological replicates since identical processing was performed for each assay (‘A’ vs. ‘B’).^5,35^ The 10x Genomics ‘GEM-X Universal 3’ Gene Expression’ kit was run on fresh cells (‘10x fresh’), alongside DSP-methanol fixed cells (‘10x fixed’), providing a reference of pre-fixation gene expression levels. All libraries were sequenced at the minimum depth recommended by vendors (20,000 reads per cell) on an Illumina NovaSeq X Plus instrument using vendor read length recommendations. Due to uneven sampling in the final library, Scale libraries were re-sequenced at the same depth with additional input to reach the targeted 10,000 cells per sample. In addition to platform-specific analysis pipelines, we applied a novel agnostic analysis approach to untrimmed data for unbiased assessment of platform performance. To do this, all technologies were down sampled to 223,082,108 reads (lowest reads, in Scale A1/A2) in both platform-specific and agnostic analyses to provide a common starting point.

### Accuracy

We assessed accuracy by comparing the expected cell type proportions in healthy PBMCs to outputs from platform-specific and agnostic cell calling algorithms using Celltypist Immune models.^36^ We then verified the robustness of cell-type assignments by reducing the number of cells sampled from each sample within a sequencing platform, termed reproducibility analysis. Per STEMCELL Technologies’ documentation, healthy PBMCs contain approximately 28.2-65.6% T-cells, 5-12.6% NK cells, 4.8-13.2% B-cells, and 5.9-18.1% monocytes (Table 2).^37^ Sample-specific results (average of biological and technical replicates) across the four assays displayed an overall high agreement between platform-specific and agnostic analysis methods (Fig 1). All values for A1/A2 were within the expected range provided by STEMCell (Suppl Table 2). Across all platforms, A1/A2 contained a median of 64.6% T-cells in platform-specific and 59.5% in agnostic analysis, demonstrating slight variation between cell calling of biological replicates across different analyses. However, a high concordance is seen in the remaining major cell populations. In both platform-specific and agnostic analyses for B1/B2, we saw a high proportion of B-cells and a low proportion of NK cells, with high concordance between biological replicates.

**Figure 1.**
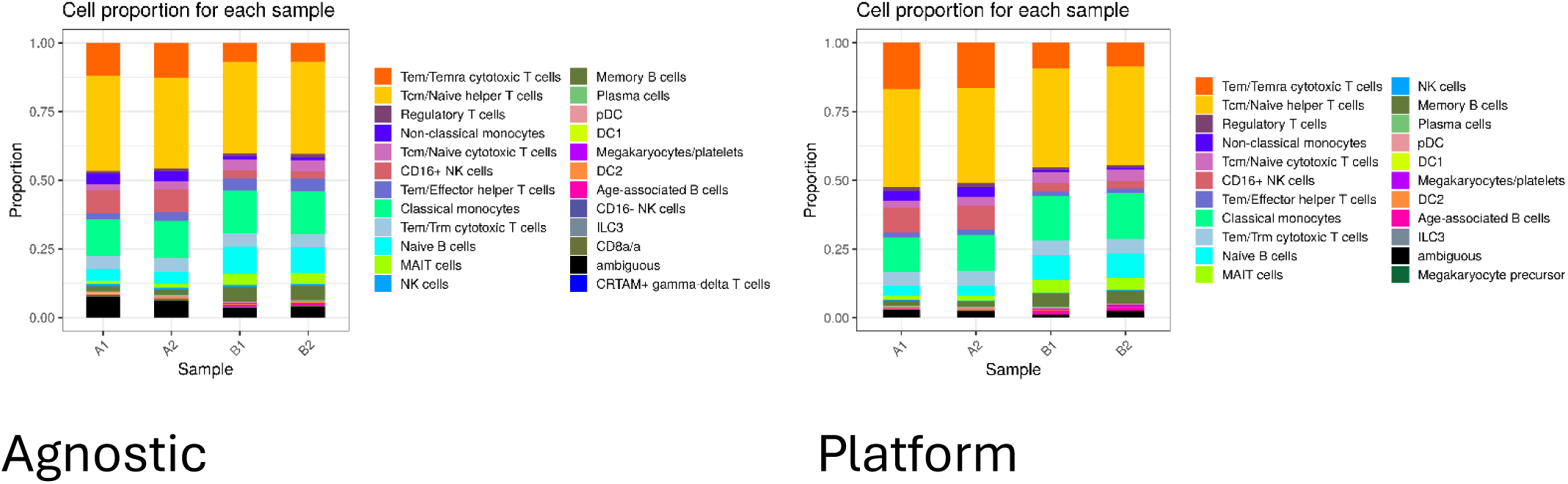
Cell type proportions per-sample, across all technologies. Plotted as proportion of cell type by sample. Color-coded by cell type (see legend). Left panel shows data from agnostic, and right panel shows data obtained from platform specific.

**Table 2.**
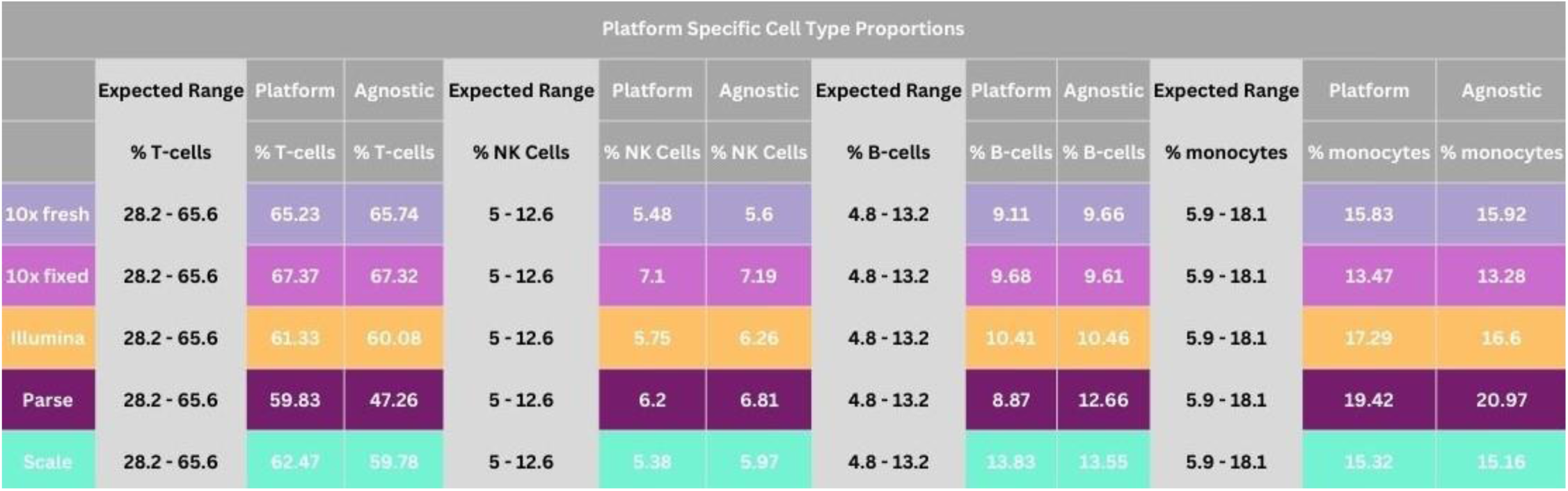
Cell Type Proportion Comparison with STEMCell data Contains broad level cell type proportions seen in reference STEMCell data, platform analysis, and agnostic analysis. Includes T-cells, NK cells, B-cells, and monocytes. Values are averaged across all samples and reported on a per-platform basis. Color-coded by platform.

While concordance was seen between cell type proportions and replicates, some variation was identified in platform-specific analyses (Table 2). For 10x fresh and 10x fixed, T-cells in platform-specific and agnostic analysis were approximately in the expected range. In Parse platform-specific and agnostic results, we reported monocytes slightly outside the expected range. Scale’s B-cell proportions were also marginally outside the recommended range in both platform-specific and agnostic results. Across both analyses, all values fell within the expected cell type proportions for Illumina. Overall, the agnostic and platform-specific analyses calculated the expected (or near the expected) proportions of cell types, indicating that each platform provides relatively accurate resolution of cell types, with the highest-level accuracy in Illumina, 10x fresh, and 10x fixed.

To determine the effect of the amount of cell input on stability of cell type identification, we performed reproducibility analysis.^38^ First, we considered the pairwise Jaccard matrix to examine the overlap between sets of sampled barcodes across iterations of cells for each analysis. Results from the Jaccard matrix across platform-specific and agnostic analyses showed that iterations worked appropriately (Suppl Fig 2). We next calculated the adjusted rand index (ARI) and percent agreement between each platform and reference labels. Platform-specific ARIs are between 0.795 – 0.815 demonstrating overall reproducibility, revealing global label similarity between full and down sampled iterations. In platform-specific results, reproducibility was high, with a slightly decreased ARI score in Scale (Fig 2A). Reproducibility was moderate in agnostic results with ARIs between 0.54-0.55, and slightly lower but still moderate reproducibility in Scale (Fig 2B). Proportion correlation verified that cell type abundances were preserved in the random subset of cells. We observed very high correlation between reference and predicted cell types across both analyses (Suppl Fig 3). To identify any unstable cell types due to false positive and negative cell type calls, we measured F1 scores of per-cell-type performance across iterations. F1 scores represent the harmonic mean of precision and recall analyses to indicate the quality of an applied model.^39^ Our platform-specific and agnostic F1 results indicated that the same cell types were observed across platforms, meaning cell types were stable (Fig 3). However, F1 analysis between platform-specific and agnostic results differed, i.e. the same cell types were conserved between technologies but not across different analytical approaches. Overall, results indicated a high degree of accuracy in all assays, with Scale’s reproducibility marginally weaker than the other technologies.

**Figure 2.**
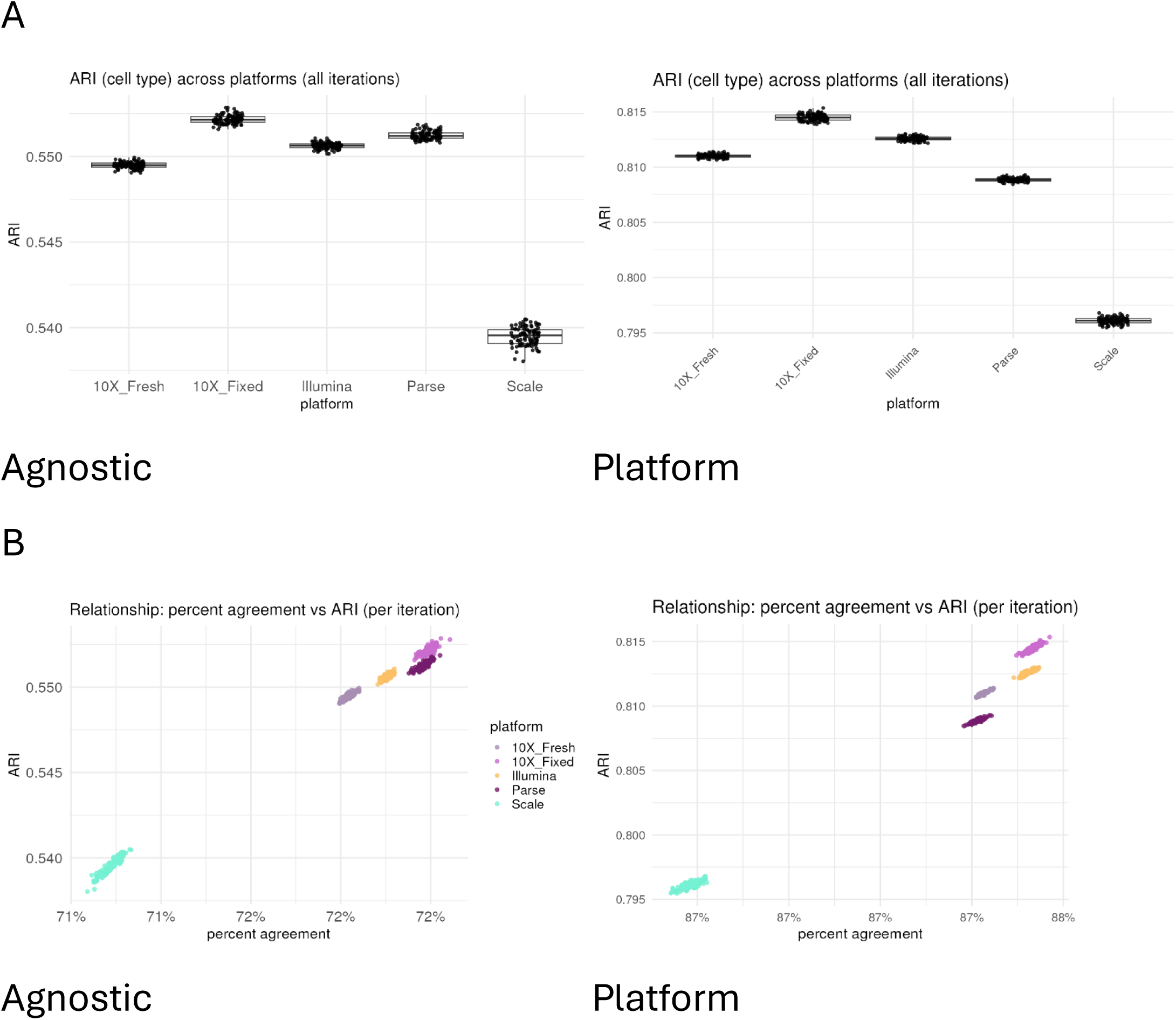
Reproducibility of single-cell platforms. (A) Adjusted Rand Index (ARI) plots, demonstrating ARI for each evaluated platform. Left panel shows data from agnostic, and right panel shows data obtained from platform specific. (B) Relationship between percent agreement and ARI, for each iteration, color-coded by platform. Left panel shows data from agnostic, and right panel shows data obtained from platform specific.

**Figure 3.**
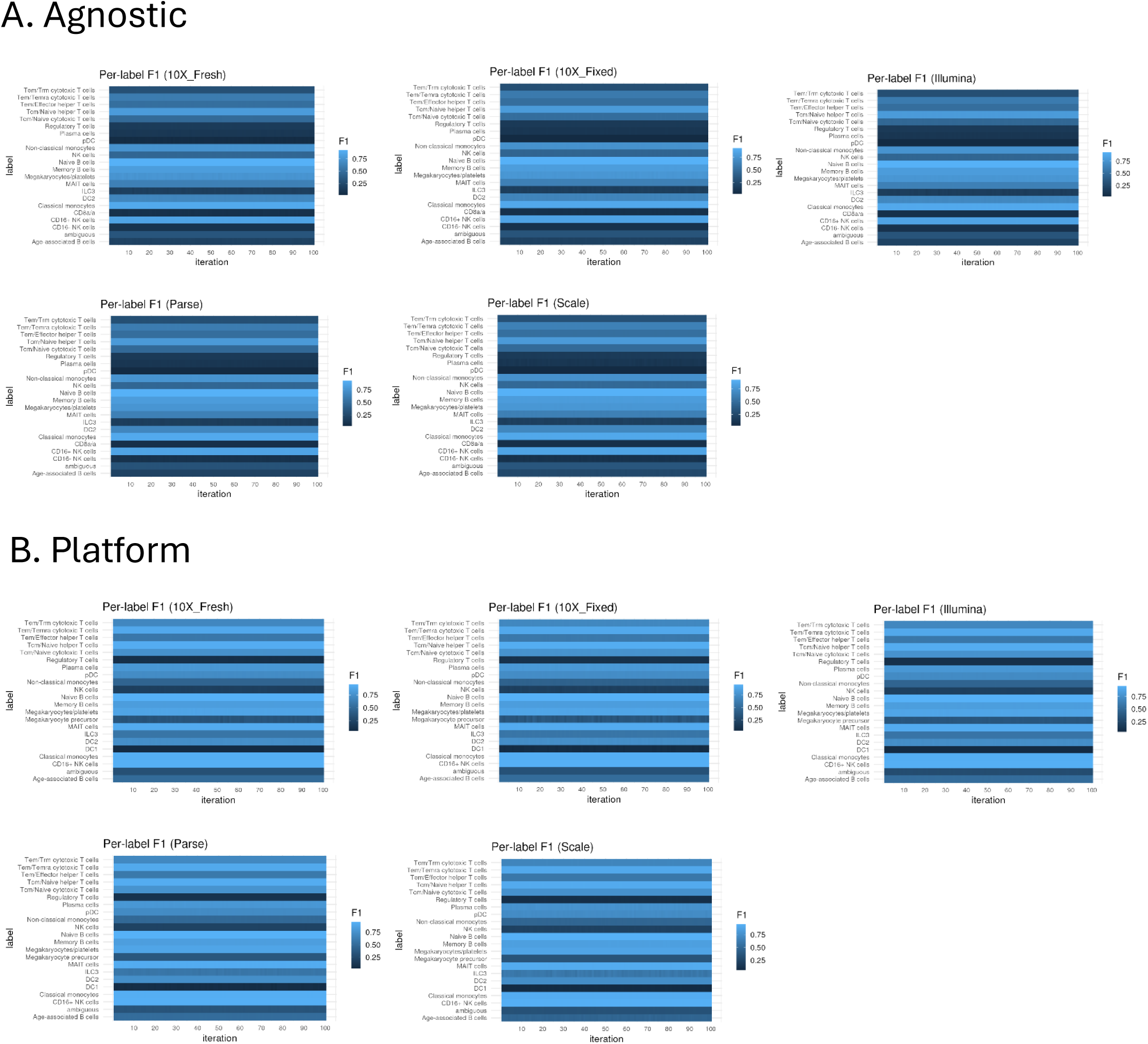
Cell-type annotation robustness under platform-specific down sampling quantified by per-class F1 scores. For the indicated platform, cells were repeatedly down sampled to a fixed number per biological sample (100 iterations; x-axis corresponds to iteration index). Per-cell type performance was summarized as the F1 score computed from the confusion matrix between reference labels and transferred labels. Heatmap rows correspond to cell types; color intensity indicates F1 score, with darker/lighter colors representing lower/higher concordance, respectively. (A) shows F1 score heatmap per cell type for samples processed by agnostic pipeline. (B) shows F1 score heatmap per cell type for samples processed by platform specific pipeline.

### Sensitivity

We assessed sensitivity as the likelihood of representing a single cell’s mRNA transcript in the final scRNAseq library.^16^ Specifically, we examined total number of detected genes, median genes per cell, the detection of low abundance (<1%) cell types on assay sensitivity, and the effect of 3’ capture.

We first counted the total number of genes detected by each platform. In the platform-specific analysis, the most genes across technical and biological replicates were detected in Scale and then Parse (Fig 4A). In the agnostic analysis, we similarly observed Scale’s detection of the most total genes, and then 10x fresh. With the agnostic approach, we observed similar median values of total genes in Parse, while significant discrepancies were seen in 10x fresh, 10x fixed, Illumina and Scale (Suppl Table 3). This effect was particularly pronounced in Scale, detecting 11,796 fewer genes in agnostic results compared to the platform-specific analysis pipeline. In terms of total genes, platform-specific and agnostic results report Scale as detecting the most genes in 10,000 cells.

**Figure 4.**
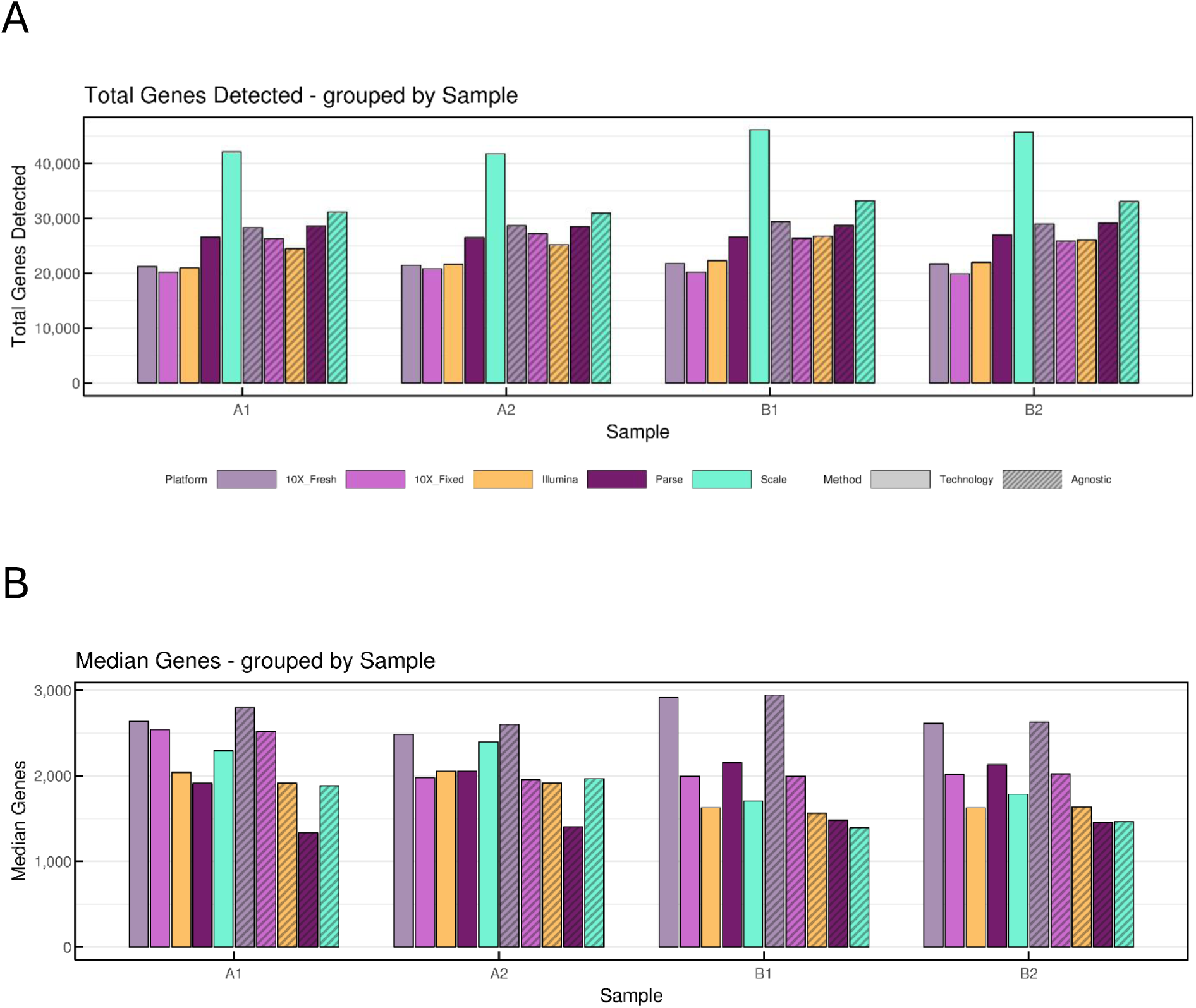
Genes detected in single-cell platforms. Solid bars represent platform analysis, and hashed bars represent agnostic analysis. (A) Total genes detected per sample, color-coded by platform. (B) Median genes detected per cell for each sample, color-coded by platform.

Next, we calculated the median genes detected per cell (Fig 4B). The median genes detected per cell also varied between platform-specific analysis and their agnostic counterpart, with minimal differences seen in 10x fresh, 10x fixed, and Illumina. In Parse and Scale, >2,000 median genes per cell were detected in platform-specific pipelines while <1,600 appeared in agnostic results (Suppl Table 3). In platform-specific analyses, the most median genes per cell were recorded in 10x fresh. In agnostic analyses, the highest median genes per cell were also in 10x fresh, followed by 10x fixed. Based on median genes per cell, 10x fresh has the highest sensitivity. For fixed technologies, platform-specific analysis results indicated the highest sensitivity in Parse and Scale while agnostic results point to 10x fixed as the most sensitive assay, indicating some differences based on the applied cell calling algorithm.

We then assessed each platform’s ability to detect rare cell types. Dendritic cells (DC), plasmablasts, hematopoietic stem cells (HSC), and progenitor cells exist at low abundances (<1%) in healthy PBMCs.^7,19^ In our study, only DCs and HSCs were detected (Fig 5, Suppl Table 2). Platform-specific analyses identified <1% DCs in 10x fresh, Illumina, and Scale, with 1% in 10x fixed and 1.1% in Parse. Across all assays in agnostic results, we identified <1% DCs. These results demonstrate that all four single cell platforms can resolve some low frequency cell types at 20,000 reads per cell, but not all. In Parse platform-specific read-outs, >1% DC indicates some variability in call calling, impacting sensitivity.

**Figure 5.**
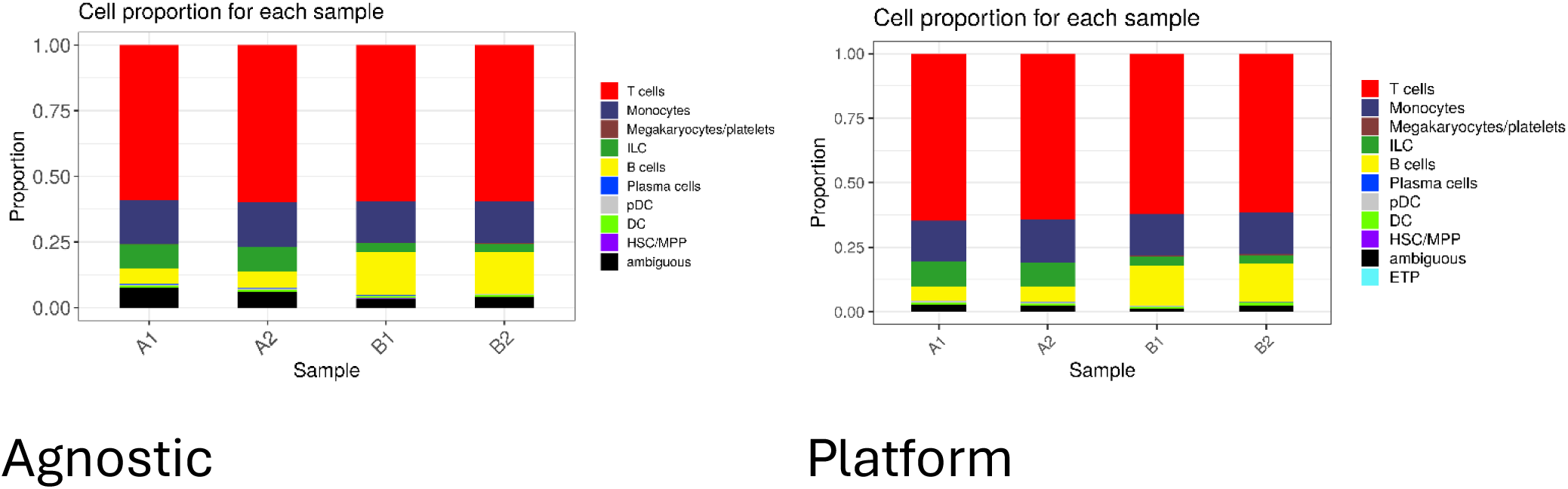
Broad-level Cell Type Proportions. Cell type proportions per sample, across all technologies. Plotted as broad cell type proportions per sample. Color-coded by cell type (see legend). Left panel shows data from agnostic analysis, and right panel shows data obtained from platform specific.

To examine the effect of 3’ capture on gene body coverage, we performed metagene analysis (Fig 6).^27^ All methods employ a 3’ capture, with additional random hexamers in Parse to facilitate whole transcriptome coverage. 10x also advertises whole transcriptome coverage through increased sensitivity in their most recent version (v4, GEM-X). Platform-specific analyses showed the expected 3’ bias in all assays, with the weakest bias in Parse’s assay. However, overall gene body coverage in the Illumina and 10x fixed assays was higher than the average normalized coverage of Parse’s assay, especially at the 5’ transcription start site (TSS). The agnostic analyses also demonstrated the expected capture-based methods’ bias at the 3’ transcription end site (TES) with the least bias in Parse (individual sample metagene coverage plots - Suppl Fig 4). The agnostic approach established that the most uniform gene body coverage, including at the TSS, is derived from the Parse platform, most likely because of the inclusion of random hexamers in the RNA amplification.

**Figure 6.**
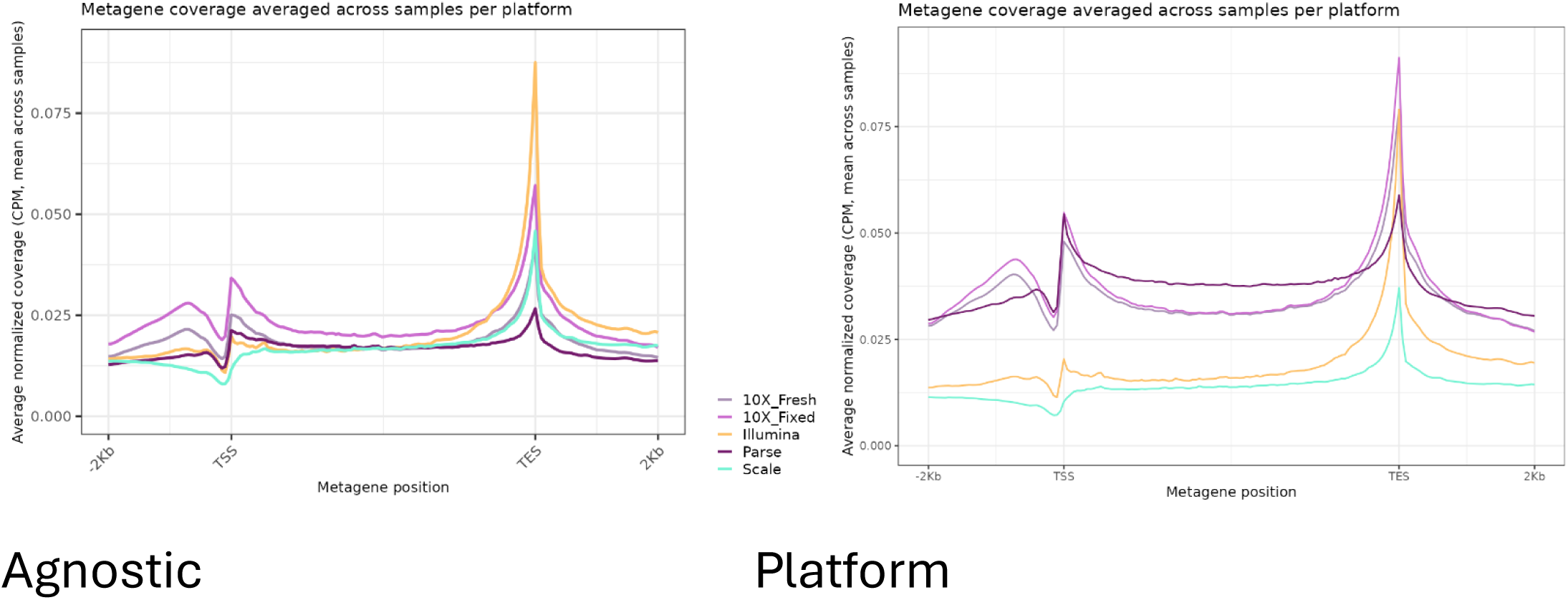
Metagene analyses. Metagene analysis/gene body coverage results, mapping the average normalized coverage (in mean Counts per Million/CPM across all samples) across metagene position. Transcription end site (TES) represents the 3’ and transcription start site (TSS) represents the 5’ end of the metagene. Color-coded by platform. Left panel shows data from agnostic analyses, and right panel shows data obtained from platform specific analyses.

Overall, when assessing sensitivity, each platform offers best performance for different metrics. Scale is the most sensitive when only considering total detected genes and results from company-provided pipelines. However, significant variation was seen between outputs from Scale’s pipeline and agnostic results, indicating substantial variability in cell calling algorithms that impact sensitivity. Scale also had the lowest counts per million (CPM) across gene bodies in platform-specific metagene results which could be due to the increased detection of non-mRNA transcripts (Fig 6). 10x and Illumina had the highest CPM when considering agnostic gene body coverage at the 3’ end, while 10x had the highest agnostic median genes per cell. Gene body coverage is skewed to the 3’ end of transcripts with Parse having the least 3’ bias and most uniform coverage of the gene, especially when considering non-mRNA transcripts.

### Precision

We assessed each platform’s precision by measuring the technical variation in each biological replicate. Since samples A and B are biological replicates, we can attribute any intra-sample variability to technical non-biological factors that contribute to zero counts, or dropout, for genes that are expressed. The dropout probability of a gene measures an assay’s precision.^13,26^ Extra-Poisson variability allows us to examine the impacts of 3’ amplification bias by quantifying amplification noise. Here, we performed precision analysis by examining the overall dropout rate across each platform and the extra-Poisson amplification noise, as demonstrated by Ziegenhain et al.^16^

Dropout rate was determined by selecting a common set of genes, analyzing these genes in a set subsample of cells, then estimating the number of cells with zero counts (Fig 7A). Across all analyses and platforms, results indicated high assay precision, with some minor variations. Using platform-specific analysis pipelines, the mean dropout rate was virtually identical between platforms, while the agnostic analysis shows 10x fresh with the lowest mean dropout rate (0.91), followed by Illumina and 10x fixed (0.93). The highest dropout rate in agnostic results was seen in Parse and Scale (0.94), suggesting the greatest technical variation and least sensitivity for the given gene set. The dropout rate between biological replicates in platform-specific and agnostic analyses demonstrated similar results, with the lowest in 10x fixed, then 10x fresh (Fig 7B). The highest per-sample dropout rate in agnostic results was observed in Parse and Scale, further indicating the most technical variation within biological replicates.

**Figure 7.**
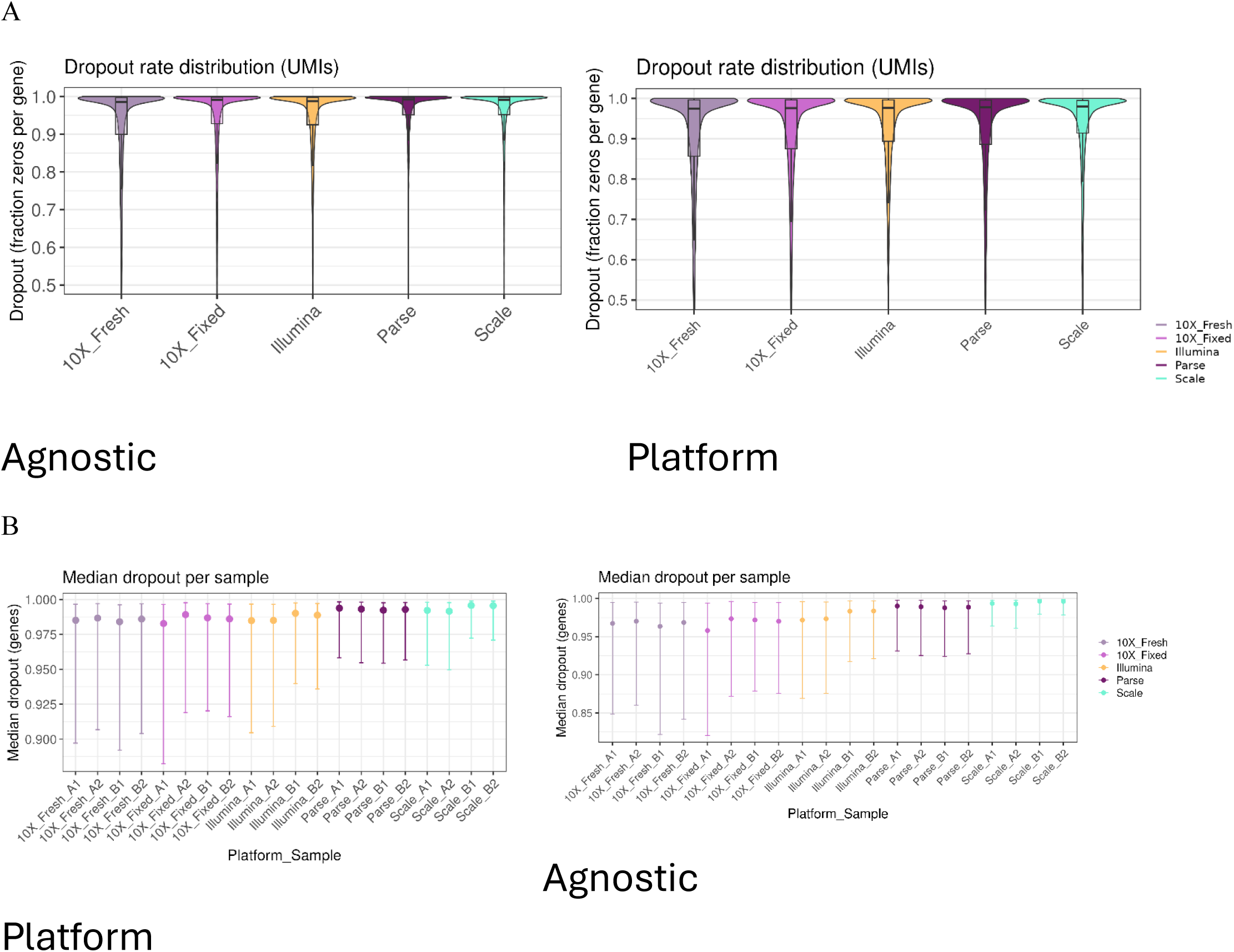
Dropout rate for each platform. Left panel shows data from agnostic analyses, and right panel shows data obtained from platform specific analyses. Color coded by platform. (A) Dropout rate for each platform, with dropout rate (fraction of zeros per gene) plotted for each platform as a violin plot. (B) Median dropout of genes per sample, modelled for each sample across platforms.

To assess amplification noise and determine the impact of technical variation, we subtracted the amount of noise expected due to Poisson sampling from the coefficient of variation (CV). All platforms demonstrated low amounts of extra-Poisson variability, with some minor differences between technologies. Platform-specific analyses showed Scale to be near the Poisson limit, demonstrating low amplification noise and high reproducibility (Fig 8, Suppl Table 4). The remaining platforms all indicated slightly higher amplification noise and thus increased per-cell library depth heterogeneity. In agnostic analysis, results varied. Illumina was near the Poisson limit, indicating the lowest amplification noise and high reproducibility. Next, Parse and Scale were over-dispersed, demonstrating minor and increased technical noise. Comparatively, 10x fixed had higher amplification noise, possibly due to RNA degradation from the fixation. 10x fresh showed the highest extra-Poisson variability, suggesting greater amplification noise and per-cell library depth heterogeneity due to increased 3’ amplification bias.

**Figure 8.**
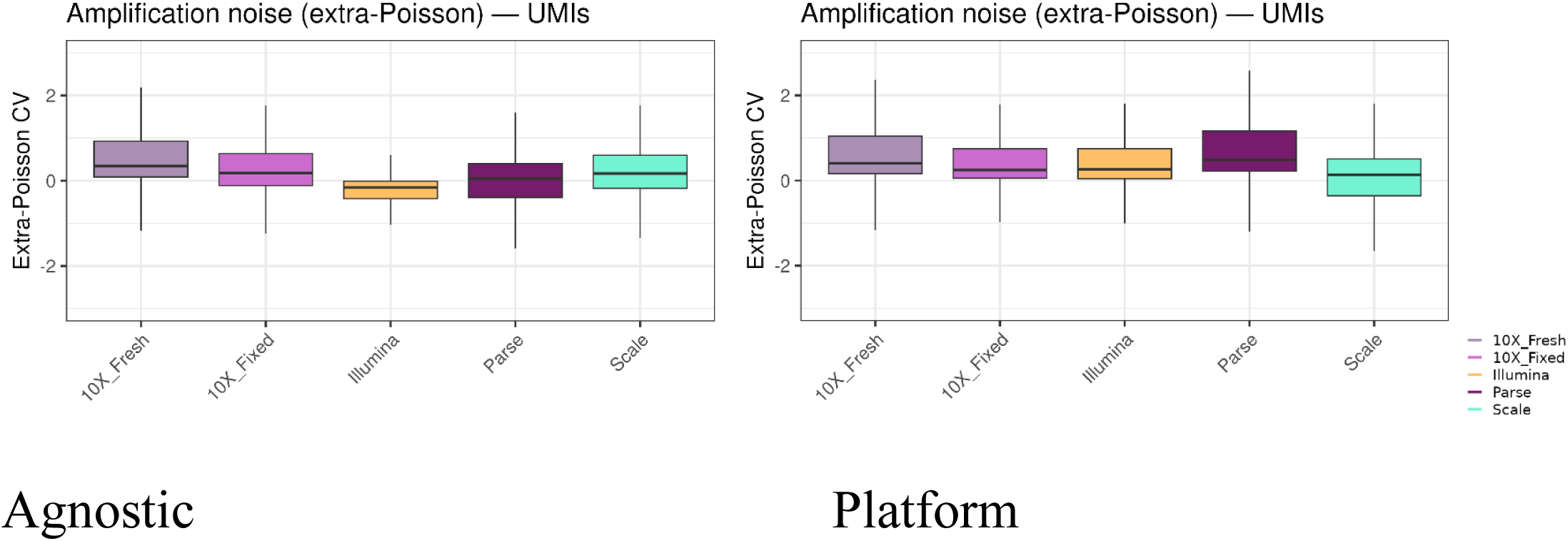
Amplification noise analyses. Amplification noise is measured as ‘extra-Poisson coefficient of variation (CV)’ for each platform, averaged across all samples, color-coded by platform. Left panel shows data from agnostic analyses, and right panel shows data obtained from platform specific analyses.

With respect to precision, 10x fresh and fixed assays demonstrated a low dropout rate and high gene detection sensitivity compared to other methods, while Illumina had decreased technical variation. Thus, in different ways, 10x and Illumina are the most precise assays.

### Power

To analyze assay power, we estimated the mean, variance, and dropout rate for each gene and method, which exposed technical variation due to batch effects and/or sequencing depth. For the purposes of these analyses, we followed the framework presented by Ziegenhain et al. by first fitting empirical noise models to capture each platform’s characteristic noise and sparsity, then simulating count data and testing differential expression (DE) between two simulations. We defined DE as genes with a Benjamini-Hochberg adjusted p-value <0.05 for a two-group t-test.

We measured performance by running 100 simulations on each gene to compute the true positive rate (TPR), representing statistical power, and the false discovery rate (FDR), representing error control. We averaged rates across replicates and summarized as power curves.^16^ The agnostic analyses for TPR showed all platforms behaving in a similar manner apart from the Illumina assay, which demonstrated substantially lower power than the other methods (Fig 9A). However, Illumina’s platform-specific results displayed similar power to other methods, indicating variability in power analysis due to cell calling algorithms. Platform-specific analyses showed broadly similar power of detection between technologies which plateaus >500 cells per group. Next, we examined FDR. Across all platforms, minimal genes were falsely discovered with increasing cell numbers in both platform-specific and agnostic analyses (Fig 9B) and within the standard deviations of each platform.

**Figure 9.**
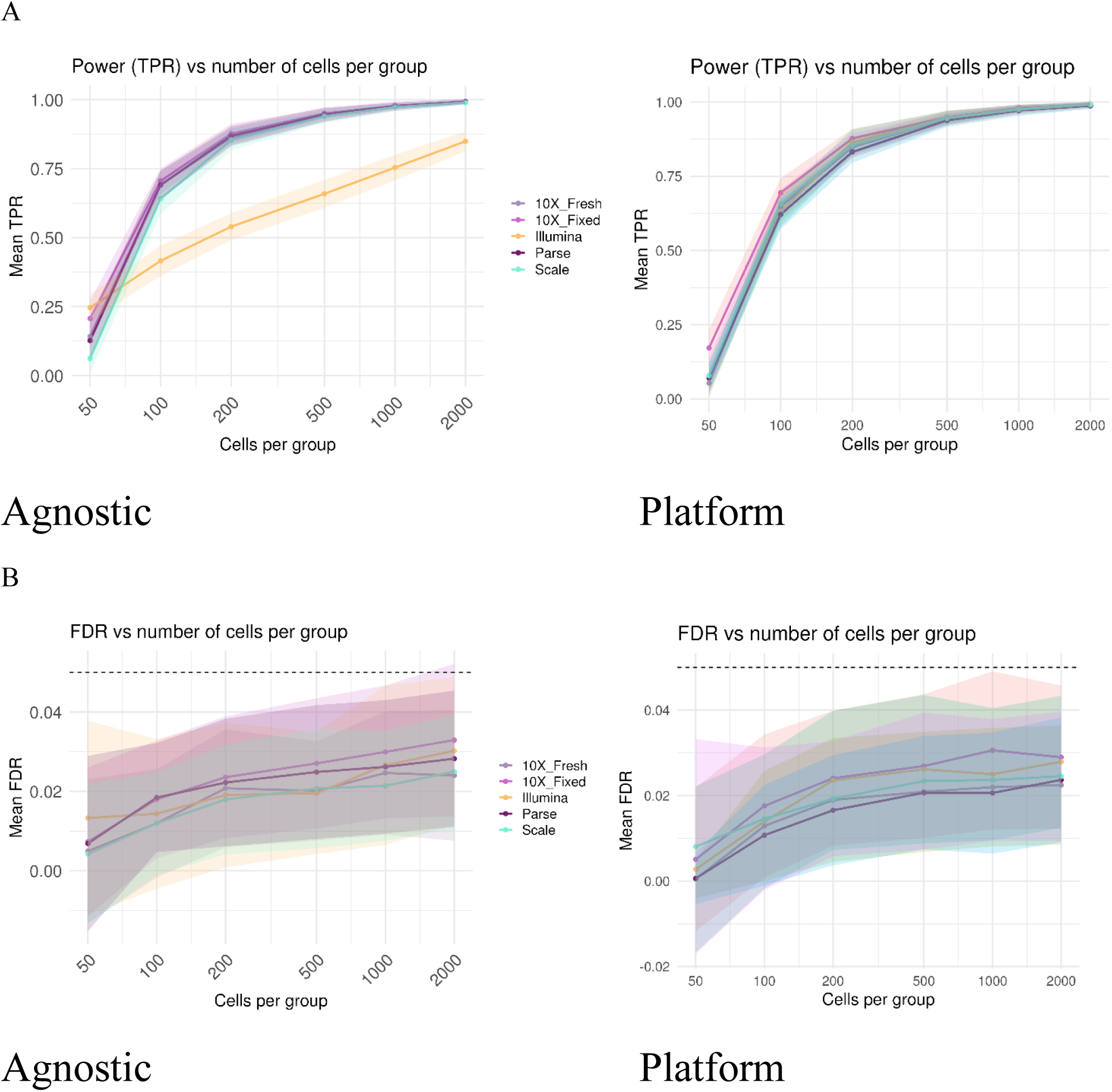
True Positive Rate (TPR) and False Discovery Rate (FDR). Left panels show data from agnostic analyses, and right panels show data obtained from platform specific analyses. (A) TPR vs Number of Cells per Group (Power). Line plot showing the mean TPR across simulation replicates (*n_reps* = 100), and shaded ribbons indicate ±1 SD across cell per platform from 50-2000 per group. (B) FDR vs Number of Cells per Group lines show the mean FDR across simulation replicates and shaded ribbons indicate ±1 SD. The dashed horizontal line marks the nominal FDR threshold of 0.05 used for significance calling.

Overall, each of the platforms had broadly similar power to detect gene expression above assay noise, with 10x fresh marginally better than 10x fixed and other assays. However, the Illumina platform had much less power than other assays in detecting differentially expressed genes (DEGs) using the agnostic analytical approach.

### Efficiency

Cost is a significant consideration when considering scRNAseq platforms. With this in mind, we calculated the overall cost of generating and sequencing single cell libraries, termed ‘efficiency’.^16^ Here, we considered the impact of user experience on efficiency in Supplemental Table 5 using metrics defined by Elz et al. as the ‘ease-of-use’ of each platform, including sample flexibility (fixation processes, storage time, input requirements), amount of pipetting, throughput potential, experimental risk of losing sample, and ease of initial data processing.^28^ These ‘efficiency’ metrics are important as they impact labor, experimental design, and overall cost. Additionally, we considered protocol clarity and difficulty, cell capture rate, sequencing saturation, multiplet rate, and the number of safe stopping points in assessing efficiency of each scRNAseq assay.

### Cost Analysis

Here, we performed an in-depth cost analysis (including sequencing and labor) to evaluate the efficiency and accessibility of each technology (Table 3). To move beyond the oft-quoted cost per cell provided by platform manufacturers, we included the costs of reagents and consumables as well as start-up costs for Illumina- and 10x Genomics-specific equipment and the service contract costs for equipment maintenance. Importantly, we also included the rate of labor, an often-overlooked cost, which is calculated based on the Association of Biomolecular Resource Facilities (ABRF) average technician’s salary, at a rate of $25/hour^40^ from protocol run times (Table 4, Suppl Table 6, and Suppl Table 8). For sequencing, prices were based on the NovaSeq X Plus 10B flow cell. A 100-cycle kit was used for calculating all sequencing costs, except in Parse, which required a 200-cycle kit. Overall, Illumina had the lowest cost per cell and is thus the most ‘efficient’ in terms of pricing.

**Table 3.** Cost Analysis Summary Cost analysis per platform, including equipment costs, equipment service agreement costs, costs of fixation (where applicable), cost of running 40K cells, and cost of running the maximum throughput for each kit. These values include labor costs. Color-coded by platform.

| scRNAseq Cost Analysis |  |  |  |  |  |
| --- | --- | --- | --- | --- | --- |
| Company | Equipment purchase (one time) | Equipment service agreement (per year) | Fixation (per rxn) | scRNAseq (per cell) when running 40k cells | scRNAseq (per cell) when maximizing the kit |
| 10x Genomics (fresh) | \$25,000 - \$65,000 | \$5,000 - \$12,000 | n/a | \$0.20 | \$0.11 |
| 10x Genomics (fixed) | \$25,000 - \$65,000 | \$5,000 - \$12,000 | \$10.09 | \$0.20 | \$0.11 |
| Illumina | \$3,500.00 | n/a | \$23.16 | \$0.16 | \$0.09 |
| Parse | \$300.00 | n/a | \$57.34 | \$0.32 | \$0.15 |
| Scale | n/a | n/a | \$30.21 | \$0.25 | \$0.12 |

**Table 4.** Protocol Timing Summary Protocol timing summary table, including the amount of total time required for fixation and each day of the scRNAseq workflow. Color-coded by platform.

| scRNAseq Protocol Timing |  |  |  |  |  |  |  |
| --- | --- | --- | --- | --- | --- | --- | --- |
| Company | Chemistry Version | # Samples | Fixation (hrs) | Day 1 (hrs) | Day 2 (hrs) | Day 3 (hrs) | Total (hrs) |
| 10x Fresh | 3' GEM-X | 4 | N/A | 2.33 | 2.75 | 3.9 | 9 |
| 10x Fixed | 3' GEM-X | 4 | 1.5 | 2.33 | 2.75 | 3.9 | 10.5 |
| Illumina | V T10 | 4 | 1 | 6.8 | 3.9 | 2.5 | 14.2 |
| Parse | v3 | 4 | 1 | 5.8 | 6 | 5.5 | 18.3 |
| Scale | v1.1 | 4 | 0.75 | 1.9 | 3.33 | 3.2 | 9.2 |

### Flexibility

Many researchers require flexible assays due to a variety of factors, including large sample sizes and long sample collection periods. Flexibility is determined by several factors, the importance of which varies between investigators and projects. Cell fixation increases flexibility by allowing longer storage times (except with Illumina) and enabling protocol selection based on equipment access, input specifications, and kit/reagent costs instead of sample availability (Suppl Table 5).^28^ To demonstrate the impact of fixation on RNA quality, we assessed RNA intactness post-fixation and found it to be sufficient for single cell profiling across all assays (Suppl Table 7). In terms of sample compatibility, Parse and Scale can only be run on fixed cells or nuclei, while 10x and Illumina can profile fresh or fixed cells or nuclei, though Illumina highly advises running fresh cells. To increase flexibility in experimental design, 10x, Parse, and Scale allow fixation of 100K-1M cells per reaction, while Illumina recommended an input of up to 1M cells. Overall, 10x is the most efficient when considering sample compatibility, allowing for fresh or fixed cells, as well as a designated lower input cell number.

The consideration of safe stopping points is also important in evaluating protocol flexibility. In this comparison, stopping points were kept as consistent as possible across assays to ensure comparable results, although samples were processed on different days (Suppl Fig 1). While these stopping points were common between all four platforms in this experiment, there are more possible stopping points in all protocols. The Parse protocol includes the most stopping points, which allows flexibility to split the assay across multiple days. However, longer stop times are enabled with 10x 3’, providing more flexibility in experimental scheduling. Assay run times were also considered, as shown in Table 4. In brief, 10x fresh and Scale had the shortest protocols, followed by 10x fixed. The longest protocol was Parse. For more detailed information on workflow timing, including sequencing and analysis, see Supplemental Table 8. Overall, in terms of protocol timing, the ideal chemistry is dependent on a project’s experimental design, samples, and available resources. However, 10x provides one of the fastest protocols with extended stopping points to increase efficiency.

### Protocol Clarity and Efficiency

Here, we assessed the impact of protocol clarity and user-friendliness on experimental efficiency. Overall, we found all protocols to be clear with varying levels of practical difficulty. While each protocol contained sufficient detail for complicated steps, they each posed specific obstacles.

The new DSP-methanol fixation from 10x Genomics presented some challenging techniques which risk sample loss. This includes a step with dropwise addition of a reagent while vortexing an open tube. Additionally, we experienced repeated clogs across three attempts of 10x fixed GEM generation and were unable to fully achieve a successful, non-clogged chip with all four fixed samples. We selected samples across multiple GEM generations to avoid analysis of clogged data. In day one of Illumina, we found the process of PIP creation to be lengthy (Suppl Table 8). Additionally, all cDNA in Illumina’s protocol must be used for library generation and PIPs can only be stored for up to one week, beyond which a full rerun of the experiment is required to prepare or re-amplify libraries. In comparison, 10x only uses a quarter of the generated cDNA for library preparation and cDNA can be stored for up to 4 weeks, leaving flexibility for re-amplification or preparations in the case of failed library construction. Similarly in Parse, cDNA is stored and barcoded samples can be kept for up to 6 months. In Scale, the final distribution plate is kept, and fixed samples can be stored for up to 1 year (Suppl Table 5).

With Parse and Scale protocols, there are many time-consuming, hands-on pipetting steps, increasing the likelihood of user error. Although the Scale protocol was straightforward in the lab, subsequent data analysis indicated uneven sample distribution with disproportionate representation of samples B1 and B2 (∼28,000 cells) compared to A1 and A2 (∼13,000 cells). To ensure the uneven distribution occurred independently of sequencing, Scale libraries were re-sequenced with double the recommended input for ∼10,000 cells/sample, allowing us to include all samples in downstream analysis. Despite re-sequencing, unequal sampling was still observed between A1/A2 and B1/B2 (Suppl Table 3). In considering protocol clarity and efficiency, the 10x fresh protocol is the simplest and most straightforward, closely followed by Illumina. However, storage times are superior in Scale, where samples can be kept for up to 1 year.

### Cell Stress

When considering cell stress, we observed similar trends to past studies,^27–29^ with some interesting discrepancies between platform-specific and agnostic pipeline results (Fig 10). In platform-specific results, we found mtRNA content to be highest in Illumina, then 10x fresh, with the lowest in Parse and Scale. In our agnostic analyses, the highest mtRNA was in the fresh 10x sample, followed by 10x fixed, with the lowest in Illumina, and Scale (Suppl Table 3). For per-sample mtRNA plots pre- and post-QC from platform-specific results, see Supplemental Figure 5. As expected, the fresh sample contained the highest mtRNA content in agnostic analyses. However, interestingly, platform-specific read-outs indicated a higher percentage in Illumina. This demonstrated some variability between Illumina’s DRAGEN pipeline and the agnostic cell calling algorithm. Across both platform-specific and agnostic results, Scale contained the lowest mtRNA content, and thus experienced the least cell stress, indicating its’ efficiency in converting mRNA to readable cDNA.

**Figure 10.**
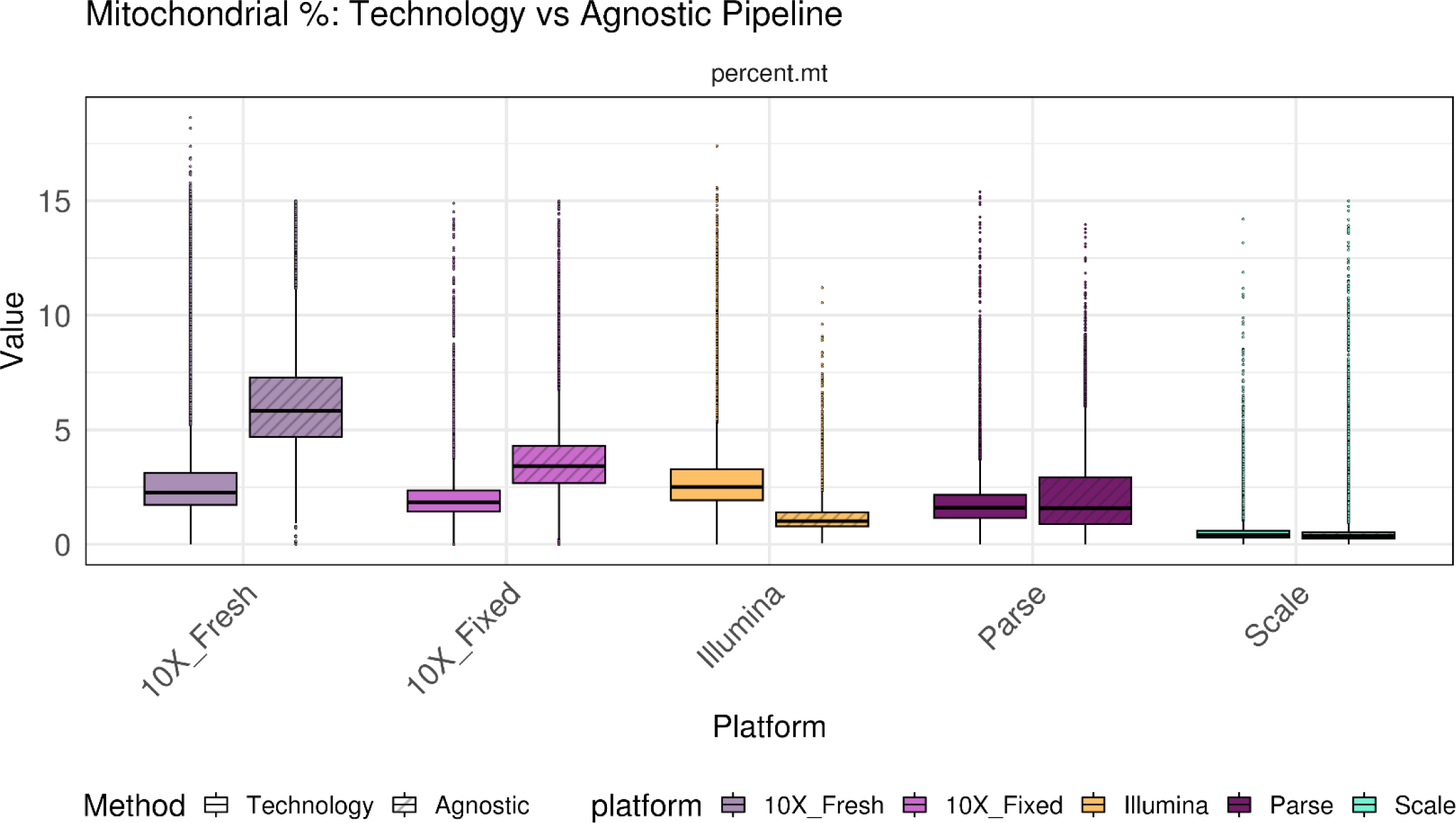
Percentage of Mitochondrial RNA Content. Percentage of mitochondrial RNA content per platform, modelled for both platform specific (solid) and agnostic (hashed) analyses. Color-coded by platform.

### Cell Capture Rate

Another factor affecting the cost of an experiment is the cell capture efficiency, which is the number of cells recovered as compared to the cellular input.^19^ Using platform-specific pipelines, we observed the highest pre-QC median cell capture rates in 10x assays. The lowest was in Parse. In agnostic data, we reported the same results (Suppl Table 3). Across both analyses, the highest cell capture rate efficiency was detected in 10x fresh, followed by 10x fixed, indicating higher efficiency in these assays.

### Multiplet Rate

Multiplets represent when multiple cells are mistakenly identified as a single cell, impacting gene expression in downstream analysis. Parse does not provide a tool for empirical doublet detection, so we employed simulation-based estimates using the consensus between SCDS and scDbtFinder.^27,34^ In platform-specific analyses, the highest median doublets were recorded in Illumina, then 10x fixed, with the lowest in 10x fresh (Suppl Table 3). For the agnostic analysis, we reported the highest in Parse, then 10x fixed, with the lowest in Illumina. Significantly more doublets were detected in the agnostic analysis, indicating substantial variation from company-provided cell calling algorithms. The agnostic analysis showed the lowest doublet rate in Illumina, while platform-specific results indicated that this assay contained the highest proportion of multiplets, demonstrating meaningful differences between platform-specific and agnostic analysis pipelines.

### Sequencing Saturation and Efficiency

The metric of sequencing saturation determines the read number at which no additional novel genes will be identified. All libraries were sequenced to the recommended minimum depth of 20,000 reads per cell. Here, we compared sequencing saturation using pre-filtration data on a common number of reads across samples (Fig 11A). For platform-specific readouts, Illumina had the highest median sequencing saturation, followed by 10x fixed (Suppl Table 2). The lowest median sequencing saturation was seen in Parse using their platform-specific analysis pipeline. The agnostic analysis showed the highest median sequencing saturation in Illumina, then 10x fixed, with the lowest in Parse.

**Figure 11.**
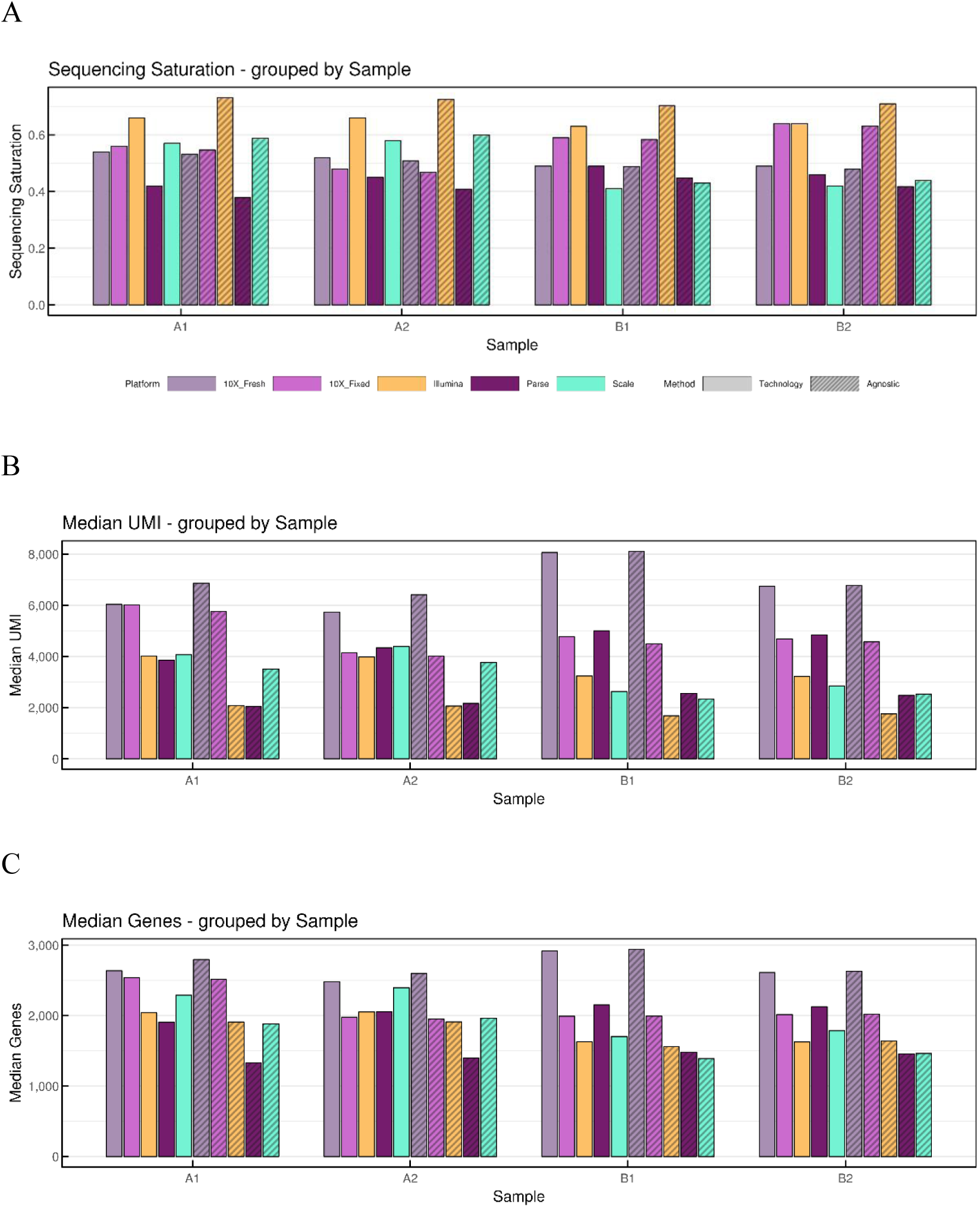
A. Sequencing statistics of single-cell platforms. Solid bars represent platform-specific measurements, and hashed bars represent agnostic measurements. Color-coded by legend. (A) Sequencing saturation for each sample. (B) Median UMI counts for each sample. (C) Median gene counts for each sample.

To evaluate sequencing efficiency, we considered read utilization by examining the fraction of reads in cells, median UMI count, and median gene count.^28^ The metric of ‘median fraction of reads in cells’ was not available from platform-specific analysis pipelines. In agnostic results, median fraction of reads in cells was highest in Scale, then 10x fresh (Suppl Table 3). Median UMI counts, across all samples and in both analyses, were highest in 10x assays (Figure 11B), while median gene counts in platform-specific results were highest in 10x fresh, then Parse (Figure 11C) and lowest in Illumina. In the agnostic analysis, median gene counts were highest in the 10x assays, with the lowest in Parse.

Overall, in the agnostic assessment of sequencing efficiency based on read utilization, the highest median fraction reads per cell were in Scale followed by 10x fresh, while the highest median UMI and gene counts were in 10x assays. Interestingly, while Parse’s platform-specific analysis shows high median gene counts, these were the lowest values in agnostic read-outs. In analyzing sequencing saturation and efficiency, while Illumina reached sequencing saturation the quickest and Scale had the highest median fraction reads per cell, the highest overall efficiency was seen in 10x assays.

### Ease of initial data processing

Another important variable influencing total assay cost is the time taken for initial data processing. Here, the ease of raw data processing was assessed based on each company’s available support documentation, ease of algorithm usage, required troubleshooting, and analysis run time for 4 samples (Suppl Table 5). 10x Genomics count pipeline is well-documented and offers a clear workflow for simple processing of initial data. Count outputs were obtained by running the pipeline on the 10x Genomics cloud. Run times can be seen in Supplemental Table 8. Illumina’s DRAGEN Single-cell RNA pipeline is also straightforward, offering clear step-by-step instructions, and a short run time. In contrast, Parse requires customers to create an account to access support documentation, we found their support site difficult to navigate, and run time was substantially longer than other assays. Parse also uses the split-pipe platform, which works on pooled samples, making it problematic to obtain sample-specific FASTQ files. We faced a similar issue in Scale’s pipeline, Scale RNA, as the latest software version does not generate FASTQ files for individual samples. Additionally, the creation of an environment for High Performance Computing (HPC) in Scale RNA was difficult to set up. Scale RNA offers two methods: a Conda environment that failed to run on HPC and Singularity, which required many adjusted NextFlow config parameters to be successful. For more detailed information on the analysis, see Methods. In summary, 10x and Illumina pipelines were the simplest and quickest to run, making them the most efficient.

### Efficiency Takeaways

Amalgamating all this information, Illumina was the most efficient assay in terms of cost, protocol clarity and efficiency (of the fixed assays), doublet rate, sequencing saturation, and ease of initial data processing. 10x was the most efficient assay in terms of flexibility, protocol timing, cell capture efficiency, read utilization and ease of initial data processing. 10x fresh was the most efficient overall in terms of protocol clarity and efficiency. Scale was the most efficient in terms of storage times and cell stress, represented as mtRNA content.

## DISCUSSION

### Accuracy

Multiple methods such as spike-in controls and known cell proportions have been used to measure accuracy.^16^ We compared the expected proportion of major cell types in healthy STEMCell PBMCs to the resolved proportion of cell types across platform-specific and agnostic analyses. Different cell prediction tools may give different proportions of cell types based on a tool’s annotation method. However, our results corresponded with previously published cell type proportions for healthy PBMCs. All platforms reproducibly resolved the same broad level of cell types at different proportions, as was observed in past studies.^15,24,28^ Differences in F1 results are attributed to variation in cell calling algorithms between platform-specific and agnostic analyses, causing different calling of cell types across algorithms. However, between platforms, cell types are conserved to indicate good analytical reproducibility. While we determined the highest degree of broad-level accuracy to be in Illumina, De Simone et al., observed that Illumina’s cell subtype proportions varied significantly from flow cytometry data, specifically in CD14+ monocytes and CD4+ (helper) T-cells. This variation was hypothesized to be due to differences in sample preparation from other workflows.^27^ In this study, CD14+ monocyte proportions were slightly outside expected values for Illumina in only platform-specific analysis, while agnostic results show CD14+ monocytes were within expected proportions. Parse, however, demonstrated values outside expected proportions in both analyses for CD14+ monocytes. Helper T-cells were identified within expected proportions in both analyses across all technologies; however, effector helper T-cells were not resolved in Parse and Scale (Suppl Table 3). This may be caused by the difficulty in resolving transcriptionally related cells (CD4+ T-cells and CD8+ T-cells)^24^, similarly observed by Elz et al.^28^ We note that CD8+ (cytotoxic) T-cells were also detected at low proportions in Parse and Scale, indicating a decreased ability in resolving certain niche cell subsets, particularly in Scale, as documented by De Simone et. al.^27^ 10x assays did not demonstrate this phenomenon, indicating successful resolution of transcriptionally related cells and niche cell subsets. We conclude that Illumina is of highest accuracy for the broad-level cell type classifications, while 10x is superior when considering cell subtypes.

### Sensitivity

We observed an increase in Scale’s total detected genes compared to other technologies. However, we also noted an uneven distribution of sequencing reads between biological replicates in Scale. After extensive troubleshooting with Scale support, we attribute this unequal sampling to Step 1.3 of the scRNAseq protocol, where counted cells are diluted to a set concentration of 2000 cells/uL, with no recommended counting step following the dilution. We recommend incorporating this additional counting step to avoid uneven sampling across the initial plate. Sampling does not explain the discrepancies between platform-specific and agnostic read-outs, however, which demonstrated substantially more genes in platform-specific results. This discrepancy indicates variability in sensitivity depending on the applied cell calling algorithm, despite both methods’ foundation in STARSolo. This discrepancy between algorithms may account for the high gene counts observed by this study and Elz et al. in Scale. Most of these genes in Elz et al. were only present in trace amounts, made up of unannotated genomic regions or having unknown biological functions.^28^

Platform-specific results indicated the highest median genes per cell in 10x fresh, (similarly to ^27,28^) followed by Parse then Scale. Agnostic results detected more total genes in 10x fresh, then Parse, with the highest median genes per cell in 10x fixed. The use of different cell calling algorithms revealed the discrepancies in sensitivity between platform-specific and agnostic analysis pipelines, indicating that other metrics must be considered in evaluating sensitivity. We considered gene body coverage and abundance of low frequency cell types, which again demonstrated substantial variability based on the applied analysis tool. Slight discrepancies with agnostic results indicated a level of pipeline-biased notation of low abundance cell types, however more sequencing depth is required to accurately resolve lowly expressed transcripts.^41^ Large differences were seen in gene body coverage results, where platform-specific outputs indicated Parse having the most uniform coverage while agnostic results pointed towards 10x fixed. Interestingly, the 3’ bias was more pronounced in platform-specific results across assays, demonstrating increased transcript abundance at the TES. However, while 3’ bias was higher in platform-specific analyses, so was the overall gene body coverage for 10x samples. We hypothesize that the variation between platform-specific and agnostic sensitivity analyses is due to the differing number of reads in cells between analysis methods.^42^ While platform-specific pipelines performed specific trimming and filtering during alignment, agnostic analysis was performed with a common alignment code for all platforms. This led to significantly differing values for reads in cells, affecting sensitivity metrics. There was a particularly noticeable decrease in Illumina and Parse’s reads per cell in agnostic results compared to platform-specific outputs, leading to decreased sensitivity in agnostic cell calling.

Compiling all this information, 10x fixed appeared to have the highest sensitivity based on agnostic results, with the highest median genes per cell and averaged normalized coverage of the gene. While Parse seemed to be of second highest sensitivity, limited conclusions can be drawn as decreased reads in cells in agnostic analyses indicate lower sensitivity. Additional sequencing would also help increase 10x fixed and Parse’s sensitivity, since, at 20,000 reads/cell, sequencing saturation was only 41.3% and 56.3% respectively in agnostic results. A similar trend in sequencing was observed by Elz et al.^28^ Overall, unless investigators are seeking information on non-protein coding or long non-coding RNAs (lncRNAs), 10x assays are recommended as the most sensitive. If you require whole gene body coverage, we recommend Parse.

### Precision

Gene detection is inversely proportional to dropout rate. Thus, it was expected for 10x fresh to have the lowest dropout as it detected the most genes per cell. The slight increase in dropout rate between biological replicates for 10x fresh and 10x fixed is likely indicative of RNA loss during the 10x fixation protocol. This was also indicated by the high extra-Poisson variability seen in the 10x fresh sample, likely reflecting technical variation due to PCR efficiency and genuine biological RNA-content differences found in fresh tissue, collectively increasing variance in UMI counts across cells. Despite the presence of degraded RNA, the 10x fixed sample demonstrated less amplification noise and per-cell library depth heterogeneity when compared to the 10x fresh sample. In platform-specific results, Scale was near the Poisson limit, representing decreased amplification noise and increased reproducibility while implying a limited 3’ amplification bias. However, across both sets of analyses, the per-sample dropout rate was high in Parse and Scale, indicating lower read distribution and less efficient gene detection in plate-based methods.^9^ In our agnostic analysis, Illumina was near the Poisson limit, indicating a decreased 3’ amplification bias due to technical noise than other methods, despite the 3’ bias seen in gene body coverage results. This indicates that the 3’ bias seen in Illumina is due to transcriptomic coverage, not amplification noise. Moderate dropout is observed in Illumina. The variation between platform-specific and agnostic results in considering assay precision again reveals the impact of different platform-specific cell calling algorithms, with agnostic results indicating Illumina had the least 3’ amplification bias and a moderate dropout rate, while the lowest overall agnostic dropout rate was seen in 10x fresh, then 10x fixed. Curiously, significantly more amplification noise was registered in platform-specific results as opposed to agnostic results, potentially due to differences in filtering between pipelines. Collectively, the low dropout rate and high gene detection sensitivity in 10x assays indicated higher transcript capture from cells as compared to other methods, but Illumina had higher precision due to decreased technical variation.^9^ If researchers are seeking to detect lowly expressed transcripts at 20,000 reads per cell, 10x is of highest precision, as previously reported.^28^ Based on our analyses, if researchers are seeking high intra-sample replicability with limited technical variation, Illumina is the most precise.

### Power

The presence of high amplification noise inflates variance across replicates, leading to increased TPRs. Illumina’s agnostic mean extra-Poisson CV value was negative, indicating that only noise due to Poisson sampling was observed in these samples. This confers a lower ability to detect differences in gene expression and thus a lower TPR, as observed in agnostic results. All other platforms produced positive mean extra-Poisson CV values and thus had amplification noise, increasing the TPR in agnostic results. Despite low TPR, Illumina demonstrated a similar FDR to other methods, indicating lower power. The negative mean extra-Poisson CV value was not observed in platform-specific results for Illumina, thus a higher TPR is detected. This indicated that the low TPR in agnostic analyses was due to differences in cell calling algorithms. Across both analyses, 10x fixed had the highest TPR, indicating that many true DEGs can be detected. However, 10x fixed also had the highest FDR, indicating stronger filtering or validation is required to assess false discoveries. Parse and Scale both had relatively stable power that misses biological effects, with low to median TPR and FDR values. 10x fixed was of highest power in detecting levels of differential expression. These results echo those of previous studies.^27,28^

### Efficiency

In our platform comparison, we restricted analysis to the often used 10,000 cells per sample and sequencing at 20,000 reads per cell. Based on this comparison, we established the Illumina assay to have the lowest cost per cell, the simplest fixed protocol and easiest data processing, akin to Elz et al.^28^ In terms of flexibility, sample compatibility, possible stopping points in the protocol, cell capture efficiency, and ease of initial data processing we found 10x to be the most efficient. We also found 10x fresh to be the most efficient protocol overall. However, we do not think the 10x DSP-methanol fixation is very user-friendly and believe it requires simplification in the applied techniques. We hope that with their acquisition of Scale, 10x will adopt methodologies from Scale’s simpler fixation protocol. Also, while Parse has the most stopping points, 10x stopping points are longer, and Scale provides the longest storage time for fixed samples; each kit provides different flexibility in designing experiments. While we determined Illumina to reach sequencing saturation the quickest at 20,000 reads per cell, agnostically 10x has the most efficient read utilization (similarly to Elz et. al), followed by Scale.^28^

In scRNAseq analyses, the presence of mtRNA transcripts may reflect metabolic or functional activity, but for the purposes of this study we used it as a metric for cell stress and low-quality cells that affect data quality and must be filtered out.^13,15,43^ Osorio et al. found that an increase in mtRNA levels lends to a decreased number of genes with detectable expression due to cell breakout events.^43^ In this study, a similar trend was observed as previous studies,^27–29^ with the highest proportion of mtRNA in agnostic analyses of the 10x fresh sample. Similarly to Elz et al., we reported a high mtRNA content in platform-specific analysis results for Illumina, potentially due to incomplete fixation of cells,^28^ although this was not observed during cell counting. We saw the lowest mtRNA content in Parse and Scale, potentially due to the many wash steps that may remove ambient RNA.^29^ The highest degree of intronic reads was also seen in Parse and Scale agnostic results, suggesting the loss of cytosolic transcripts during fixation (Fig 12). Interestingly, there were more intronic reads in 10x fresh than 10x fixed and Illumina in platform-specific analyses, implying that 10x and Illumina’s fixation protocols do not lose as many cytosolic transcripts, which has implications for RNA trajectory analyses.^44^

**Figure 12.**
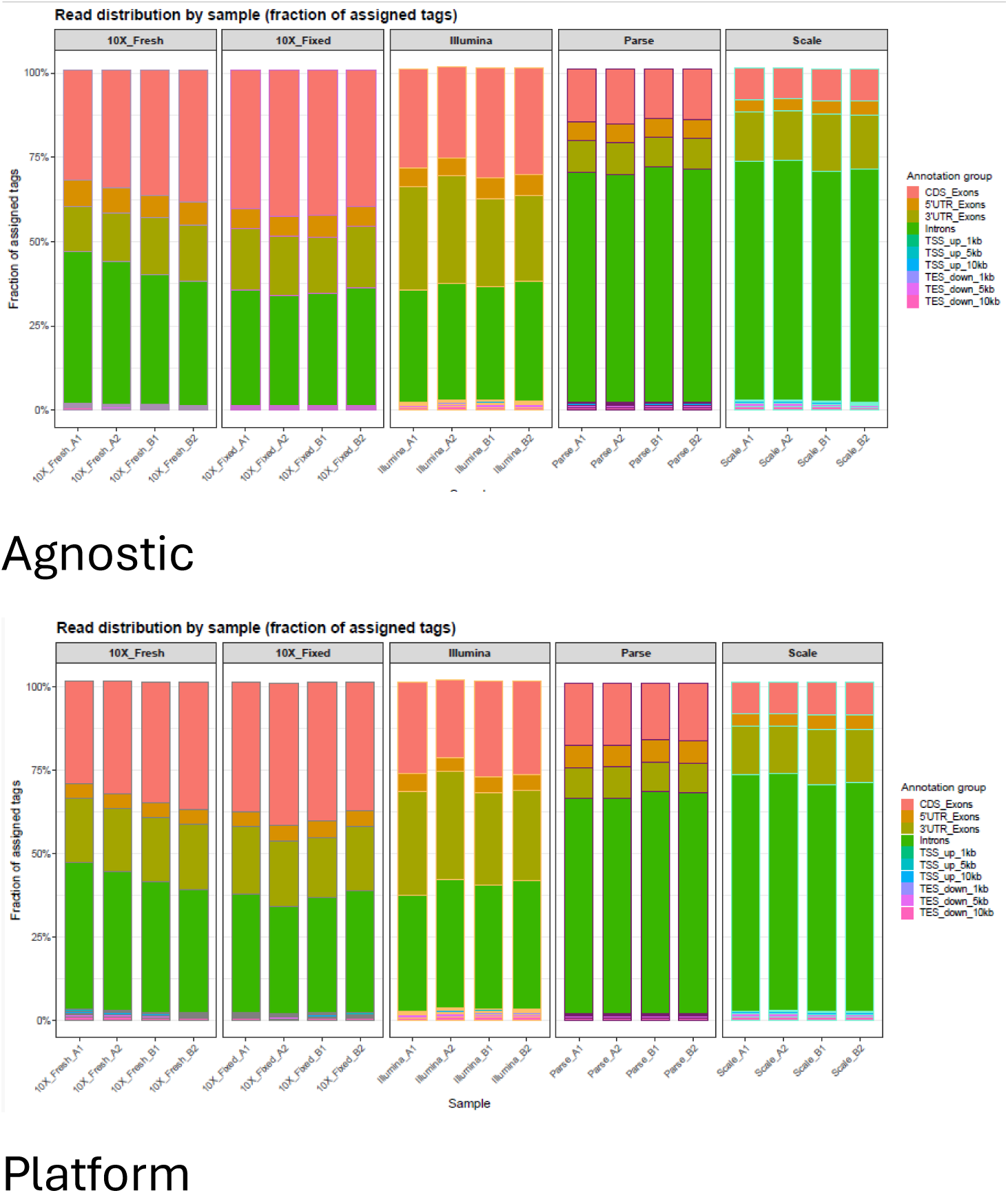
Read distribution. Read distribution by sample, representing the number of reads that are tagged as intronic, exonic, from the transcription start site (TSS), or from the transcription end site (TES). Plotted as fraction of assigned tags by sample. Color-coded by tag. Top panel shows data from agnostic analyses, and bottom panel shows data from platform-specific analyses.

Increased multiplets were detected in the agnostic analysis versus the platform-specific analyses, with the highest in Parse (8.5%). This is consistent with previous studies that identified a high multiplet rate in Parse due to uneven RNA content in samples or cell clumping.^15,28,34^ In platform-specific results, Parse’s multiplet rate was much lower (0.96%), indicating variation due to the cell calling algorithm. We hypothesize that this is due to differences in trimming and filtering that occur during alignment in company platforms.

Similarly to Elz et al., we found Illumina and 10x to have the simplest data processing using company-provided pipelines. While running Parse and Scale pipelines, we obtained pooled FASTQ files. However, for subsequent quality control steps, per-sample FASTQ files were required. Parse support page has its own codes which we were able to use to get per-sample FASTQ files. Alternatively, Scale’s latest version v2.0.1 does not have defined method to obtain FASTQ file for each individual sample. The older version v1.6.3 had a method to generate FASTQ files for each sample which was also confirmed by Scale Biosciences’ support team, so we used this version for our analysis. This was an arduous process to follow with eventual desired files.

### Study Limitations

Similarly to De Simone et al. and Elz et al., we were limited to healthy PBMCs, necessitating further studies across a diverse set of sample and cell types and disease states. We were also unable to obtain flow cytometry data for our specific PBMC lot numbers, limiting the precision of conclusions drawn from our ‘accuracy’ metric. Due to budget constraints, sample size and targeted number of cells were limited. For a benchmark of kits with maximum throughput, see Elz et al.^28^ Assays were run on different aliquots of matched healthy PBMCs purchased from STEMCELL on different days and batch effects may have been introduced, which were removed in downstream analysis. Additionally, in the period following data generation for this study, newer or different versions of these assays that may maximize throughput through multiplexing have been introduced. For example, a newer ‘Quantum Scale’ kit and new 10x Chromium kit, including multiplexing for smaller cell numbers that can increase scalability and decreased cost per cell. Likewise, improved platform-specific analysis may improve the analysis of data.

## CONCLUSIONS

Our study provides experimental and analytical evaluation of the strengths and weaknesses associated with four commercially available scRNAseq 3’ oligo (dT) assays, including 10x Genomics new 3’ DSP-methanol fixed assay. Overall, we found Illumina to be most accurate at the broad-level cell type classifications, best in precision in terms of sample replicability, and best in ‘efficiency’ for data generation. We determined 10x fixed to be most accurate when identifying cell subtypes, overall sensitivity, having the best precision when considering lowly expressed transcripts at minimum sequencing depth, and the highest power in detecting levels of differential expression. Profiling outside the 3’ UTR or of non-polyadenylated mRNAs may be benefitted by Parse’s broad isoform discovery and examination of alternative splicing events, however some level of this information is also accessible with the 10x GEM-X assay. In summary, our extensive evaluation across multiple metrics provides clarity to researchers to determine which is the best commercially available platform for their scRNAseq needs for diverse biomedical and clinical applications.

## MATERIALS & METHODS

### Cell Thawing & Counting

To maintain consistency across platforms, we thawed 2 vials of healthy PBMCs (STEMCELL Technologies® Human CAT #70025.1, Vancouver, BC, Canada) from liquid nitrogen storage, following 10x Genomics® ‘Cell Thawing Protocols for Single Cell Assays – PBMCs/Cell Lines – Dropwise Media Thawing.’ For each single cell technology, 2 PBMC samples were prepared and sequenced in duplicate. We designated LN 2301805101 as donor ‘A’ and LN 230272202C as donor ‘B’. Post-thaw & platform-specific fixation of both donor’s samples, we split each sample into 2 technical replicates. We measured cell number and viability using a Revvity Cellometer K2 instrument and corresponding consumables: Revvity ViaStain AOPI Staining Solution and Revvity Cellometer SD100 Cell Counting Chambers (PN CMT-K2-MX-150, PN CS2-0106-5ML, PN CHT4-SD100-002, Revvity, Walton, MA). We measured cell counts in duplicate on the K2and averaged for input into fixation and single cell technologies.

### Cell Fixation

We piloted fixation before the actual run. Full results are found in Supplemental Table 1.

### Scale Low Volume Fixation for Single Cell RNA Sequencing

We performed fixation following Scale Biosciences ‘Low Volume Fixation for Single Cell RNA Sequencing Kit’ User Demonstrated Protocol with the ‘ScaleBio Sample Fixation Kit’ (Product Code 2020001, Scale Biosciences, San Diego, CA). When prompted (Cell Preparation step 5, Cell Fixation steps 10 & 13), the tube was gently flicked x10 to resuspend. Based on fixed cell recovery and quality, we found no need for optimization.

### 10x Genomics® Fixation of Cells for GEM-X Single Cell 3’ & 5’ Assays

We performed fixation following ‘10x Genomics’ Cell Fixation Protocol for GEM-X Single Cell 3’ & 5’ Assays.’ Following initial testing, we increased centrifugation speed to 500 x g for 10mins, using a fixed angle centrifuge, to improve cell recovery. See Supplemental Table 1 for more information. After adding Dehydration Buffer at step ‘e’, we pipette mixed the sample 3 times. At step ‘r’, approximately 100-200uL of supernatant was left then we mixed 2-3 times before resuspension.

### Parse Evercode Cell Fixation v3

We fixed cells following Parse Biosciences ‘Evercode Cell Fixation’ protocol, using ‘Evercode Cell Fixation v3 kit’ (PN ECFC3300, Parse Biosciences, Seattle, WA). We started the protocol from step 1.2 with use of Protein Lo-bind Eppendorf tubes (Cat. No. 022431081, Eppendorf, Hamburg, Germany). Per manufacturer recommendations, we evaluated fixed cells with a hemocytometer alongside automated cell counter, using Trypan Blue stain, to measure precise cell concentration. Based on fixed cell recovery and quality, we found no need for optimization.

### Illumina DSP-Methanol Fixation for Cells

We performed fixation using Illumina ‘DSP-Methanol Fixation for Cells protocol’ and ‘Pipseq V T10 3’ Single Cell RNA Kit Bundle’ according to manufacturer recommendations, without adjustments (PN 20135691, Illumina Inc., San Diego, CA). Based on fixed cell recovery and quality, we found no need for optimization.

### RNA Extraction & Quality Check

We extracted RNA from fresh & fixed cells to verify intact RNA post-fixation for each single cell platform. See Supplemental Table 7 for more information.

### Single Cell cDNA library preparation

We performed single cell RNA sequencing according to each manufacturer’s instructions, with the following adjustments. Full results are found in Supplemental Table 1.

#### Scale Single Cell RNA Sequencing Kit, v1.1

Immediately following fixation, we prepared single cell cDNA libraries with the ‘Single Cell RNA Sequencing Kit, v1.1’ (Product Code 950884, Scale Biosciences, San Diego, CA) using a multi-channel pipette. We loaded duplicate aliquots of each sample (A1, A2, B1 & B2) into the RT Barcode plate, as technical replicates See Supplemental Figure 4 for our Scale experimental worksheet, including RT Barcode plate loading. Throughout the protocol, we centrifuged barcoding plates at 1000 x g for 1min in 4°C to collect all the reagent at bottom of wells. In Step 2 of the protocol, column 24 rows B, D, F, H, J, L, N, and P were missed during pipetting of Ligation Master Mix into 384-well Ligation Barcode Plate, leading to decreased final barcode count. For more information, see Results section.

The following day, we thawed samples on ice and continued the protocol from Step 4 through 4.17. Prior to both thermal cycler programs, we added a hold step at the program’s starting temperatures, per FAS recommendation, to allow equilibration of the cycler block.

On the third day, library preparation was performed from Step 5. Following Tagmentation (Step 5.8, Program 6), we added a 22°C 5 min step to the thermal cycler program, then samples were immediately removed to stop the Tagmentation reaction. At the end of Tagmentation Stop program (Step 5.16, Program 7), a 4°C hold step was incorporated to increase workflow flexibility. We then quantified the final pooled library’s concentration with the Invitrogen Qubit dsDNA BR Assay Kit (REF Q32856, Thermo Fisher Scientific, Waltham, MA) and diluted to 1ng/uL for final size quantification with the Agilent TapeStation HS D5000 kit (PN 5067-5592 &PN 5067-5593, Agilent Technologies, Santa Clara, CA).

#### 10x Genomics® Chromium Single Cell Universal 3’ Gene Expression

Immediately following cell counting of fresh or fixed cells, we proceeded with Gel Bead-in-Emulsion (GEM) generation following ‘Chromium GEM-X Single Cell 3’ Reagent Kits v4 User Guide, using the Chromium Single Cell Universal 3’ Gene Expression kits (PN 1000711, 10x Genomics, Pleasanton, CA). We loaded duplicate aliquots of each sample (A1, A2, B1 & B2) into individual wells of the microfluidic chip, as technical replicates.

On day 2, we purified GEMs then amplified and QC’ed cDNA (Step 1.4-2.4). We measured cDNA concentration and size with a 1:5 dilution on Agilent TapeStation system, with HS D5000 consumables.

The following day, we prepared libraries from Step 3.1, based on the TapeStation’s estimated cDNA yield. For fresh samples, we ran the SI Index PCR (Step 3.5), and purified libraries the following morning. We purified fixed libraries immediately after Index PCR. We quantified the concentration and size of libraries (1:10 dilution) on the Agilent TapeStation, using HS D1000 consumables (PN 5067-5584 &PN 5067-5585, Agilent Technologies, Santa Clara, CA).

#### Parse Biosciences Evercode WT v3

We began the Parse Biosciences Evercode WT v3 protocol immediately following fixation, loading duplicate aliquots of each sample (A1, A2, B1 & B2) into Barcoding Round 1 plate as technical replicates. See Supplemental Figure 5 for sample loading table. Per Parse training recommendation, at Step 1.2.9.v., we pooled all wells of the barcoding round 1 plate, 2 at a time, into a 15mL tube, using a multi-channel p200 set to 180uL.

The next day, we processed 4 of the 8 prepared sub-libraries for cDNA amplification (Step 2.1). At step 2.5.2, we diluted cDNA to load ∼3ng/sample onto the Agilent TapeStation system with HS D5000 consumables.

On the following day, we thawed cDNA on ice for library preparation from Step 3.1. Following the Post-Barcoding Round 4 Size Selection (Step 3.6), we quantified the concentration and size of libraries (1:10 dilution) on the Agilent TapeStation, using HS D1000 consumables.

#### Illumina Pipseq V T10 3’ Single Cell RNA Kit

Immediately following fixation, we proceeded to the ‘PIPseq V T10 3’ Single Cell RNA Kit’ user guide, using ‘Pipseq V T10 3’ Single Cell RNA Kit Bundle’ (PN 20135691, Illumina Inc., San Diego, CA). We loaded duplicate aliquots of each sample (A1, A2, B1 & B2) into individual PIP-containing tubes, as technical replicates (Step 4.1.13).

The next day, we isolated and reversed transcribed mRNA into cDNA with whole transcriptome amplification (Step 5.2-5.4). We quantified isolated cDNA concentration using the Qubit HS dsDNA assay kit (Cat. No. Q32851, Thermo Fisher Scientific, Waltham, MA), then diluted to approximately 1-2ng/uL for cDNA size quantification using the TapeStation HS D5000 consumables.

We used the entire volume of cDNA to prepare libraries (Step 5.7). We quantified final library concentration (1:10 dilution) using Qubit HS dsDNA assay kit then we quantified the size of libraries (1:10 dilution) on the Agilent TapeStation, using HS D1000 consumables.

#### Library Quantification, Sequencing & Initial Data Processing

We quantified the concentration of each library using the Roche KAPA Library Quant kit for Illumina sequencing (REF #07960336001, Roche Diagnostics, Indianapolis, IN) and sent samples for sequencing at the Duke Sequencing and Genomic Technologies (SGT) core facility. The SGT measured library concentration using Qubit 1X dsDNA HS Assay Kit (ThermoFisher Cat. No. Q33231). To calculate average library size, the SGT used Fragment Analyzer HS DNA kit (Agilent Part Numer: DNF-474-0500) with a range of 200 – 1000 bp apart from samples prepped with the Scale protocol. Scale Libraries had broader peaks, and to properly calculate library size, a range of 200 – 3000 bp was used (Supplemental Figure 6). To minimize batch effects, all libraries were loaded according to manufacturers’ specifications on the same Illumina NovaSeq X Series 10B flow cell, 200 cycle, Reagent Kit (Cat. # 20085595, Illumina, San Diego, CA). Each library type (I.E. 10x fixed, 10x fresh, Illumina 3’, Scale, and Parse) was sequenced on a different lane of the flow cell at a targeted sequencing depth of 20,000 reads/cell. Additional Scale sub-libraries were rerun at a separate time on a 1.5B flow cell with the same conditions as the first sequencing run.

We converted BCL files to FASTQ files using Illumina’s bcl-convert (Version 4.3.13) conversion software. We performed raw sequencing data quality checks using Babraham Institute’s ‘FastQC’ (Version 0.11.9) and Seqera’s ‘multiQC’ (version 1.27.1) open-source aggregation quality control tools. To run FastQC we used command: fastqc -o /output_dir/ *_R1.fastq.gz *_R2.fastq.gz and to run MultiQC we used command: multiqc /fastqc_dir.

#### Read Down Sampling and Preprocessing

To minimize biases due to differences in sequencing depth, we randomly down sampled raw FASTQ files from each sample to match the sample with the lowest total read count—223,082,108 reads from the Scale A2 sample. We used seqtk v1.2-r94 (https://github.com/lh3/seqtk) with a fixed random seed of 100 to ensure reproducibility across runs. Both Read 1 and Read 2 FASTQ files were down sampled proportionally. We used the resulting down sampled files for all subsequent preprocessing and alignment steps, including barcode and UMI extraction.

To Run Scale and Parse technology count pipelines we required downsampled pooled fastq files. To get the downsampled pooled fastq, reads from original pooled fastq where the sequence identifier matches with the individual downsampled fastq sequence identifier were utilized which was suggested by technology customer support.

#### Cell Calling – Technology Specific Pipeline

We performed cell calling with each samples’ platform-specific pipeline to map, demultiplex, and quantify single cell data. For Parse Biosciences samples, the Split-Pipe pipeline (v1.5.0) with chemistry version 3 was employed, utilizing all sublibraries per sample to generate cell count matrices. For 10X Genomics Fixed and Fresh samples, Fastq files were processed through 10X Genomics CellRanger v9.0.1 - GEM-X 3’ Gene Expression v4 on cloud analysis platform, while samples generated through Illumina’s workflow were analyzed with the DRAGEN Single Cell RNA pipeline (v4.4.2) available on BaseSpace. Scale Biosciences’ samples were analyzed using their tool ScaleRNA (v1.6.3) which incorporates Nextflow workflow with Singularity. Scale pipeline used STARsolo (v2.7.10b) as part of their cell calling pipeline

### Agnostic Pipeline

#### Barcode and UMI Extraction for STARsolo Input

To enable accurate demultiplexing and quantification with STAR (version 2.7.11b), we preprocessed down sampled FASTQ files to extract and restructure cell barcodes and unique molecular identifiers (UMIs) according to the respective library structures. We created new FASTQ files containing cell barcodes and UMIs for each of the single-cell RNA seq platforms detailed bellowed: *10x Genomics*. We extracted the first 16 bases of Read 1 as the cell barcode and the next 12 bases as the UMI. New FASTQ files contained only the sequence and quality strings of these 28 bases. We modified the read header to append the barcode and UMI in the format @readID_barcode-UMI.

#### Illumina

We assembled the cell barcode from four non-contiguous sequence regions of Read 1 (bases 1–8, 12–17, 21–26, and 32–39), while the IMI was extracted from bases 40–42. We concatenated these components to create a synthetic 30-bp read comprising the barcode and IMI, with corresponding quality scores, and a similarly modified read header as in 10x Genomics.

#### Parse Biosciences

We assembled the cell barcode from three non-contiguous sequence regions of Read 2 (bases 10–17, 30–37, and 50–57) while the UMI was extracted from bases 1 - 10. We concatenated these components to create a synthetic 30-bp read comprising the barcode and UMI, with corresponding quality scores, The newly assembled sequence and quality strings were written into a modified FASTQ file, maintaining the original read identifiers.

#### Scale Biosciences

FASTQ files provided by the vendor after fastq conversion already contained pre-extracted barcode and UMI information in the expected format compatible with STARsolo i.e. it had both cell barcode and UMI sequence.

### Read Alignment and Gene Quantification with STARsolo

To map reads to the human genome, we first aligned reads from all sequenced libraries to the GRCh38 human genome reference (Ensembl release 113) in STARsolo mode. We used a consistent set of parameters across all technologies, with adjustments to barcode and UMI positions to reflect platform-specific read structures. Alignment was performed with 8 threads using the --soloType CB_UMI_Simple option, and platform-specific values for --soloCBstart, --soloCBlen, --soloUMIstart, and --soloUMIlen were specified according to the structure of the modified FASTQ files. Cell barcodes were not filtered using predefined whitelists (--soloCBwhitelist None), and EmptyDrops_CR was used as the filtering method to identify cell-containing droplets (--soloCellFilter EmptyDrops_CR). The GeneFull_Ex50pAS option was used under --soloFeatures to count gene expression from exonic and partially spliced reads. Only uniquely mapped reads were retained (--soloMultiMappers Unique), and BAM output was sorted by coordinate (--outSAMtype BAM SortedByCoordinate).

All runs included output of comprehensive alignment tags (NH HI nM AS CR UR CB UB GX GN sS sQ sM) and STARsolo matrix files (features.tsv, barcodes.tsv, matrix.mtx). We used the same genome index and the same GTF annotation file (Homo_sapiens.GRCh38.113.gtf) for all alignments. We used the final outputs for downstream quantification and cell-level analysis.

#### Binning 3bp IMI’s in Illumina Cell Count Output

The read structure for Illumina libraries includes a 3-base molecular “bin” (IMI) that is used to distinguish fragments originating from the same molecule. To account for this we reprocessed the STAR-aligned BAMs with an IMI-aware counting pipeline: we derived an empirical barcode include-list from the data, mapped raw CB/CR tags to that include-list, grouped reads by (corrected CB, gene) and collapsed reads into molecules by counting unique (BI, IMI, alignment-start) combinations and applying an IPM-based correction. The IMI-aware pipeline was used for Fluent/PIPseq to respect the chemistry’s molecular labeling and to avoid misinterpretation of the 3-base bin as a conventional UMI.

#### Down Stream analyses in Seurat

We processed both technology specific and agnostic pipeline cell count data using the following parameters. We processed and analyzed raw single-cell count matrices (matrix.mtx, barcodes.tsv and features.tsv) using R (Seurat v5.2.0) and a small supporting Python script for barcode collapsing as required for Parse. We imported each sample with a robust reader function and created seurat objects for each sample with CreateSeuratObject(counts = mat, project = <sample>, min.cells = 10, min.features = 0) (the min.features = 0 argument allowed creation of raw objects prior to downstream filtering). Each Seurat object received two metadata columns, platform (e.g., 10X_Fresh) and sample (e.g., A1), so all objects were traceable by origin.

#### Quality Control and filtering

We calculated the percent mitochondrial and ribosomal content using Seurat’s function PercentageFeatureSet() using pattern = “MT-” and “^RPS|^RPL”. We calculated QC metrics for each sample (number of cells, median UMIs per cell, median genes per cell, median percent.mt) using Matrix::colSums() and Matrix::colSums(counts > 0). Based on visualization of the distributions of percent mitochondrial reads, we filtered out low-quality cells. We subset each Seurat object to contain 1) between 500 and 20,000 transcripts (nUMI (nCount_RNA)); 2) between 250 and 6,000 genes (nFeature_RNA) and <15% mitochondrial content (percent.mt < 15%).

### Doublet annotation and consensus calling

Doublet detection was performed using two orthogonal approaches and recorded in the Seurat object metadata:

1. *scds*: co-expression (cxds), binary classification (bcds), and the hybrid cxds_bcds_hybrid (with estNdbl = TRUE) were run via the scds package Cells called “TRUE” by the hybrid were annotated as scds_doublet = TRUE.
2. *scDblFinder*: scDblFinder::scDblFinder() was run on each per-sample SingleCellExperiment conversion. Cells with scDblFinder.class != “singlet” were annotated as scDblFinder_doublet = TRUE.

We removed cells TRUE for both approaches, for downstream applications.

### Parse barcode collapsing (random hexamer handling)

For Parse Bioscience (combinatorial indexing) libraries, barcode collision due to random hexamer or indexing differences can lead to redundant barcodes mapping to the same canonical cell identifier. We implemented a barcode collapsing procedure and run after QC and doublet removal. We treated the last 18 bases of each cell barcode as a canonical key (suffix length = 18) and grouped barcodes sharing the same canonical suffix. For each group, the corresponding columns of the counts matrix were added to produce a single “collapsed” cell column (UMIs summed by gene). The collapsed matrix was assembled as a sparse dgCMatrix and replaced the Seurat counts for the Parse sample. We retained metadata columns to match collapsed barcode names. This logic was implemented both in a Python prototype (using scipy sparse matrices) and ported to a memory-efficient R implementation that operates on dgCMatrix.

### Ambient RNA decontamination (DecontX)

We estimated and corrected for ambient RNA per sample using DecontX implemented in the celda package (celda::decontX()), operating on sparse matrices to avoid dense coercion. For each sample, we created a SingleCellExperiment object from sample counts (assays = list(counts = counts_pre)), preserving sparsity (dgCMatrix). We ran celda::decontX() to estimate and correct for ambient RNA and extracted the corrected counts with the subset function assays(sce_decont)[[”decontXcounts”]].

### Cell-cycle scoring

To reduce noise due to cell cycle heterogeneity, we normalized each sample (NormalizeData()) and calculated cell cycle phase scores using the function CellCycleScoring() with the cc.genes.updated.2019 gene lists (s- and g2m-gene sets). We computed S.Score, G2M.Score, Phase, and the difference CC.Difference = S.Score - G2M.Score, all of which were stored in metadata for normalization.

### Per-sample SCTransform

We normalized counts for each sample using Seurat’s SCTransform(). Prior to merging, cells were optionally renamed with a sample-specific prefix to guarantee unique cell names across samples. vars.to.regress could include CC.Difference and perecent.mt.

### Final QC summary metrics

For each sample a combined summary table was produced by merging Seurat-derived matrix metrics with STARsolo *_Summary.csv metrics. Calculations and sources are summarized below:

- **From Seurat matrix (post-processing)**: n_cells (number of columns), median_umi (median per-cell UMIs), median_genes (median genes detected per cell), umis_in_cells (sum of UMIs across cells), total_genes_detected (genes with ≥1 UMI).
- **From summary file (sequencing/mapping)**: total_reads, reads_in_cells (reads assigned to cell barcodes), sequencing_saturation, unique_map.
- **Derived metrics**: frac_in_cells = reads_in_cells / total_reads, umis_per_read = umis_in_cells / total_reads, genes_per_umi = median_genes / median_umi, cells_per_million_reads = n_cells / (total_reads / 1e6), mean_reads_per_cell = total_reads / n_cells.

#### Creating a joined assay (layer-joining) for cross-sample analyses

To support cross-sample usage of the per-sample normalized values, the RNA assay layers were combined via JoinLayers() into a joined assay containing counts layer. The joined assay was set as the active assay for normalization/variable-feature detection and subsequent analyses.

#### Variable feature selection, scaling, and cell-cycle scoring

A new joined assay was normalized (NormalizeData) and highly variable features were identified with FindVariableFeatures(selection.method = “vst”). Cell-cycle scores were computed with Seurat’s CellCycleScoring() function. Prior to PCA, ScaleData() was applied to the joined assay with vars.to.regress = c(”CC.Difference”, “percent.mt”) to regress out cell-cycle and mitochondrial effects.

#### PCA and Harmony integration

To observe global data trends, we performed principal component analysis with RunPCA(…, npcs = 50). To integrate platform effects while retaining biological variability, we ran Harmony (RunHarmony) using group.by.vars = “platform.” We use Harmony-corrected embeddings for downstream neighbor graph construction and visualization.

#### Dimensionality reduction and clustering

UMAP embedding was computed on Harmony-corrected PCs using RunUMAP(reduction = “harmony”, dims = 1:30). We computed nearest neighbors with FindNeighbors(reduction = “harmony”, dims = 1:30) and identified clusters with FindClusters() over a vector of resolutions (c(0.6, 0.8, 1.0, 1.2, 1.4, 1.6, 1.8, 2.0)). We evaluated clustree plots to inspect cluster consistency across resolutions, and selected a final resolution of 1.6

#### Lightweight object and CellTypist annotation

We annotated cell types in CellTypist (Python). We first created a lightweight Seurat object from the full object containing 1) RNA assay data slot set to the joined normalized expression matrix (sparse) and 2) metadata and selected reductions (pca, harmony, and umap) for each cell. This lightweight object was exported to H5Seurat and converted to AnnData/H5AD (SeuratDisk::SaveH5Seurat() + Convert(…, dest = “h5ad”)) and loaded in Python for CellTypist annotation. We ran CellTypist with two models, Immune_All_Low.pkl and Immune_All_High.pkl (models downloaded with celltypist.models.download_models(force_update = TRUE)). We predicted cell types with majority_voting = True and mapped back to cell barcodes. The selected fields (predicted_labels, majority_voting, conf_score, over_clustering) were exported to CSV.

#### Visualization and cluster annotation

CellTypist predictions for each model (low/high) were bound to combined Seurat object metadata. Cells with low confidence scores in both high- and low-models (confidence < 0.5 for both) were labeled as “ambiguous” and the corresponding majority_voting labels replaced with “ambiguous.” We used DimPlot() to create a UMAP of CellTypist-derived labels (high_majority_voting, low_majority_voting) and unsupervised clusters. We generated views split by sample and platform. We fixed factor levels as cell types and assigned custom color palettes to major cell types to ensure consistent plotting across figures. Per-sample and per-platform stacked barplots of CellTypist-derived cell-type proportions were produced using ggplot2 (position = “fill”) and saved as PNGs.

### Pseudobulk sample-level analysis

To compare samples to one another, we performed pseudobulk analyses. We computed pseudobulk counts by summing counts per gene across cells for each platform–sample group (metadata column group_sample = paste(platform, sample)). We converted counts to CPM (counts per million) and retained genes with CPM > 1 in at least two pseudobulk samples. We transformed CPM into log2(CPM + 1). We performed PCA of the top 2000 variable genes in log2CPM per-sample platform. We calculated the Pearson correlations between pseudobulk samples (log2CPM), which we visualized as a heatmap (pheatmap).

#### Dropout summary and per-cell detection

We also computed:

- Overall fraction of zero gene counts per platform (via Matrix::nnzero()), and per-cell detection rates (per-cell fraction of genes detected).
- For each platform we reported (i) mean dropout per gene, (ii) median dropout per gene, (iii) proportion of genes with dropout = 1 (genes never detected), and (iv) the distribution of per-cell detection rates.

#### Reproducibility analysis

We evaluated label robustness to reduced cell yield using platform-wise down sampling on the combined Seurat object (joined assay). We selected 2,000 HVGs and computed a fixed PCA reference embedding (PC1–PC30) from gene-wise z-scored log-normalized expression. For each platform, cells within each biological sample were down sampled without replacement to a fixed target per sample (platform-specific minimum by default) and repeated for 100 iterations using deterministic iteration- and sample-specific seeds. Down sampled cells were projected into the reference PCA using the stored PCA loadings, and labels were predicted by k-nearest neighbor (k=10) majority vote in the reference embedding for cell-type annotations and clustering labels. Agreement between predicted and reference labels was quantified using adjusted Rand index (ARI) and percent agreement; cell-type performance was further summarized using per-label precision, recall, and F1. Sampling diversity across iterations was assessed using Jaccard overlap of sampled barcode sets, including a full pairwise iteration-by-iteration Jaccard matrix.

#### Precision and Power analysis

To compare platforms, we performed platform-level precision and power analyses using Seurat objects (RNA assay). For each platform, raw UMI count matrices from all samples were concatenated after restricting to the intersection of genes shared across platforms. Dropout rate was computed per gene as the fraction of cells with zero counts and summarized across genes per platform. Extra-Poisson variability was computed per gene as extra-CV = (SD/mean) − (1/√mean), where mean and variance were estimated using sparse-matrix operations, and distributions were compared across platforms. To estimate DE power, we fit platform-specific mean–variance–dropout relationships from the aggregated UMI counts: dispersion was derived as (variance − mean)/mean² (truncated at ≥0) and smoothed versus mean using a spline, while dropout probability versus mean was modeled using LOESS. Using these fitted relationships, we simulated two-group count matrices by sampling per-gene counts from a negative binomial distribution and applying dropout by setting a fraction of values to zero. Differential expression was introduced by applying a log-fold change (logfc = 1) to 5% of genes in one group. For each simulated dataset, genes were tested for differential expression with per-gene two-sample tests and Benjamini–Hochberg correction (FDR < 0.05). Power was summarized as TPR (fraction of truly perturbed genes detected) and FDR (fraction of detected genes that were not perturbed). Simulations were repeated for 100 iterations per platform using 2000 genes and a grid of n_cells per group = 50, 100, 200, 500, 1000, 2000.

#### Metagene coverage analysis

We performed a metagene coverage analysis using deepTools (3.5.6). For each sample, reads from the final filtered cell BAM (restricted to barcodes passing the cell-calling/UMI filtering step) were converted to per-base coverage tracks using bamCoverage with CPM normalization, a fixed bin size of 50 bp, and duplicate reads ignored. These normalized BigWig files were then summarized across annotated genes using computeMatrix scale-regions in metagene mode. Gene bodies were scaled to a fixed length of 5,000 bp and flanked by 2,000 bp upstream of the transcription start site (TSS) and 2,000 bp downstream of the transcription end site (TES). Per-sample profiles were generated with plotProfile (mean across genes), and a combined matrix was additionally computed across all samples to facilitate cross-sample visualization. For platform-level summaries, coverage profiles were averaged across samples within each platform at each bin, enabling direct comparison of metagene shape and relative enrichment near the TSS or TES as indicators of 5′ or 3′ bias.

#### Sequencing saturation analysis

Sequencing saturation was assessed from raw UMI count matrices stratified by platform. Genes were restricted to those expressed (UMI > 0) in at least 100 cells across the dataset. For each platform and target depth from 2,000–35,000 reads per cell (2,000-step), only cells with total counts ≥ that depth were retained; depths with <200 eligible cells were discarded and, when >10,000 cells were available, a random subset of 10,000 cells was used. Counts were downsampled to the target depth by binomial thinning (downsampleMatrix, scuttle), and the median number of detected genes per cell was computed. The relationship between median genes and depth was modelled with a Michaelis–Menten function using non-linear least squares, yielding platform-specific saturation curves and half-maximal depth (*K*ₘ).

#### Cumulative gene discovery

To examine how rapidly each platform covers the expressed gene space, we computed cumulative gene discovery curves. For each platform, up to 10,000 cells were randomly selected and counts were restricted to genes expressed in ≥100 cells overall. The selected cells were randomly permuted 100 times; for each permutation and for a series of cell numbers between 1 and the total, we recorded the number of distinct genes with at least one UMI in the union of the first *k* cells. For each platform and *k*, the mean and standard deviation of the cumulative gene count across permutations were reported and visualized as mean curves with ±1 SD ribbons.

#### Read distribution across genomic annotation classes

Aligned, coordinate-sorted BAM files were used to quantify read distribution across genomic features. Transcript annotations corresponding to the reference genome build used for alignment were converted from GTF to BED12 format and validated for BED12 compliance. For each sample, genomic feature utilization was quantified using RSeQC (*read_distribution.py version 5.0.1*) with the BED12 transcript annotation. Per-sample outputs were saved as text reports.

Downstream aggregation and visualization were performed in R. RSeQC reports were parsed into a tidy format, samples were mapped to sequencing platforms based on sample identifiers, and tag counts were normalized by total assigned tags per sample to obtain per-feature fractions. Feature distributions were summarized at the platform level (mean across samples per platform) and visualized using stacked bar plots.

#### Software and versions

Analyses were implemented in R (version 4.4.0; Seurat v5.2.0), with major packages: Seurat, SeuratDisk, Matrix, SingleCellExperiment, celda (DecontX), scDblFinder, scds, harmony, ggplot2, pheatmap, patchwork, dplyr, tidyr, tibble. CellTypist annotation was performed in Python (CellTypist v1.7.1) using Immune_All_Low.pkl and Immune_All_High.pkl models; AnnData/scanpy were used for data I/O (H5AD). Exact package versions and the full code used are provided in the project repository.

## ACKNOWLEDGEMENTS

We would like to acknowledge the use of Future House AI during our literature search and LLM model (ChatGPT) for help with coding. We would also like to acknowledge the use of Biorender in generation of the graphical abstract and Figure 1. Thank you to Wes Austin (previously at Scale Biosciences Inc.) and Bob Wilke (previously at Scale Biosciences Inc.) for their comprehensive support. We would like to thank Steve Rozen for bioinformatic advice. Thank you also to Odmaa Bayaraa, Emily Hocke, Alan Smith, and Julia Smith for assistance in manuscript preparation. This project was funded by the Duke University School of Medicine Core Facility Voucher.

