## Supplemental Figures for "A platform-agnostic evaluation of non-formalin fixed single cell RNA technologies"

### Supplemental Figure 1 - Experimental Design

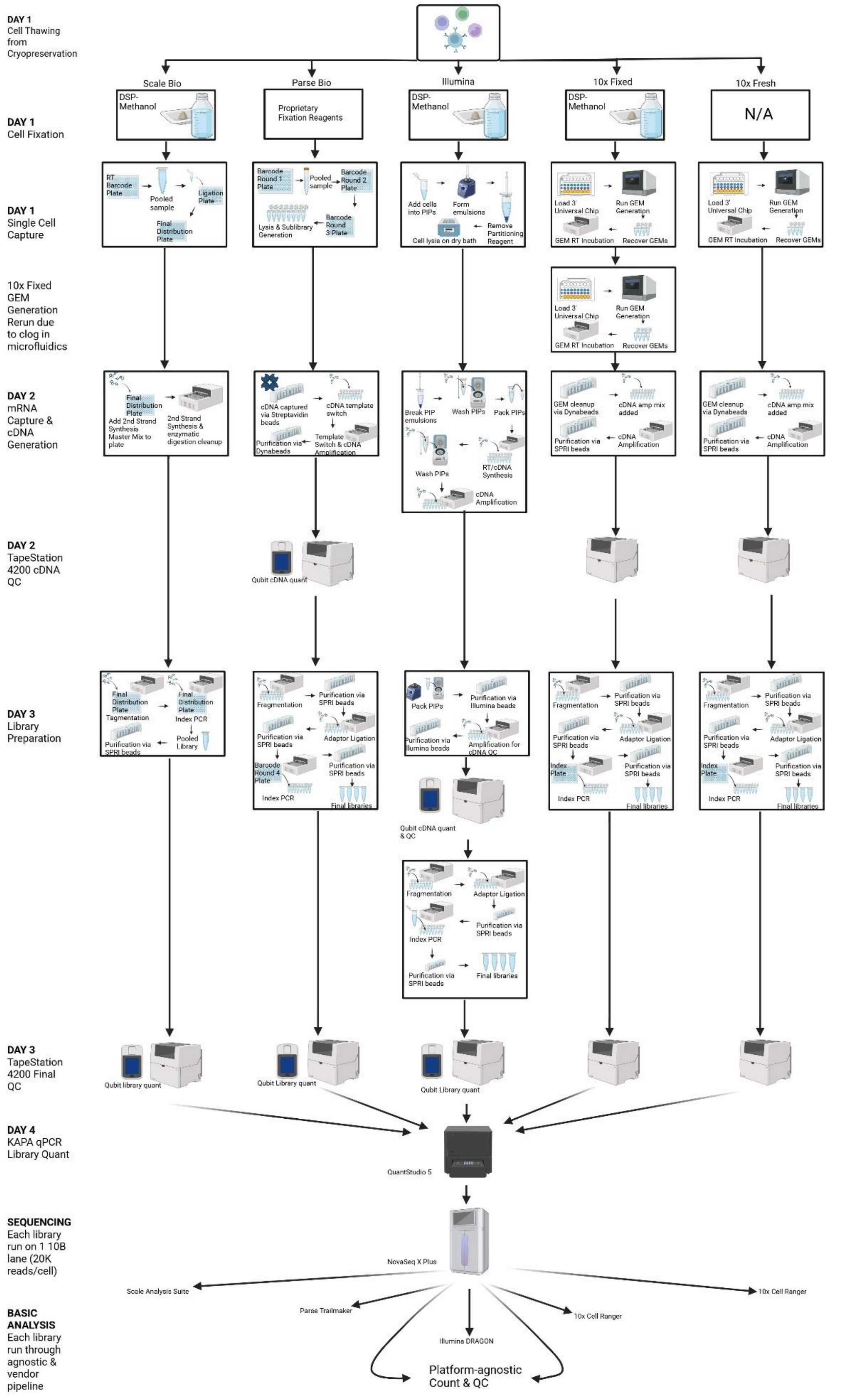


Supplemental Figure 1 – Flow chart of experimental design, broken down by step of scRNAseq experiment

#### Supplemental Table 1 – Fixation & Assay Details

| Fixation & Assay Details | | | | | | | | | | | | | | | |
|---|---|---|---|---|---|---|---|---|---|---|---|---|---|---|---|
| | Thawing Protocol | Fixation Protocol Details | A Fixation Cell Input | A Fixation Cell Input Viability | B Fixation Cell Input | B Fixation Cell Input Viability | Optimized Centrifugation Conditions for Fixation | A Assay Input | A Assay Input Viability | B Assay Input | B Assay Input Viability | Assay Cell Targeting | Assay Protocol Details | # cDNA Cycles | # PCR Cycles |
| 10x Fresh | 10x Genomics CG000447 Rev. A | N/A | N/A | N/A | N/A | N/A | N/A | 14,876 | 95.35% | 14,742 | 89.85% | 10,000 | 10x Genomics, CG000731, Rev B | 11 | 14 |
| 10x Fixed | 10x Genomics CG000447 Rev. A | 10x Genomics, CG000776 , Rev. B | 1,000,000 | 95.10% | 1,000,000 | 84.75% | optimized with 10x R&D: fixed angle benchtop centrifuge 500 x g for 10mins at 4C | 15,246 | N/A | 16,849 | N/A | 10,000 | 10x Genomics, CG000731, Rev B | 11 | 14 |
| Illumina | 10x Genomics CG000447 Rev. A | Illumina, Doc ID: FB0004708, Rev. 1 | 1,000,000 | 95.10% | 1,000,000 | 84.75% | 500rcf for 5min at 4°C | 16,998 | N/A | 17,500 | N/A | 10,000 | Illumina, ID: FB0004762 Revision 1.5 | 12 | 15 |
| Parse | 10x Genomics CG000447 Rev. A | Parse Biosciences, UM0027, v1.1 | 1,000,000 | 98.20% | 1,000,000 | 92.40% | 400 x g for 10mins at 4C | 163,469 | N/A | 163,894 | N/A | 10,000 | Parse Biosciences, Evercode WT v3 User Manual, v1.4 | 8 | 10 |
| Scale | 10x Genomics CG000447 Rev. A | Scale Biosciences, Document 1020807, Rev E, Jul 2024 | 430,000 | 95.55% | 786,000 | 91.40% | 500 x g for 10mins at 4C | 136,800 | N/A | 136,800 | N/A | 10,000 | Scale Biosciences, Document 1020796, Rev B, Jun 2024 | N/A | 14 |

Supplemental Table 1 – Details on assays performed, including fixation information, any fixation optimization (10x Genomics), fixation input amounts, assay input amounts, assay protocol details, number of cDNA cycles, and number of PCR cycles. Color-coded by platform.

**Supplemental Figure 2 – Jaccard Index Plots**

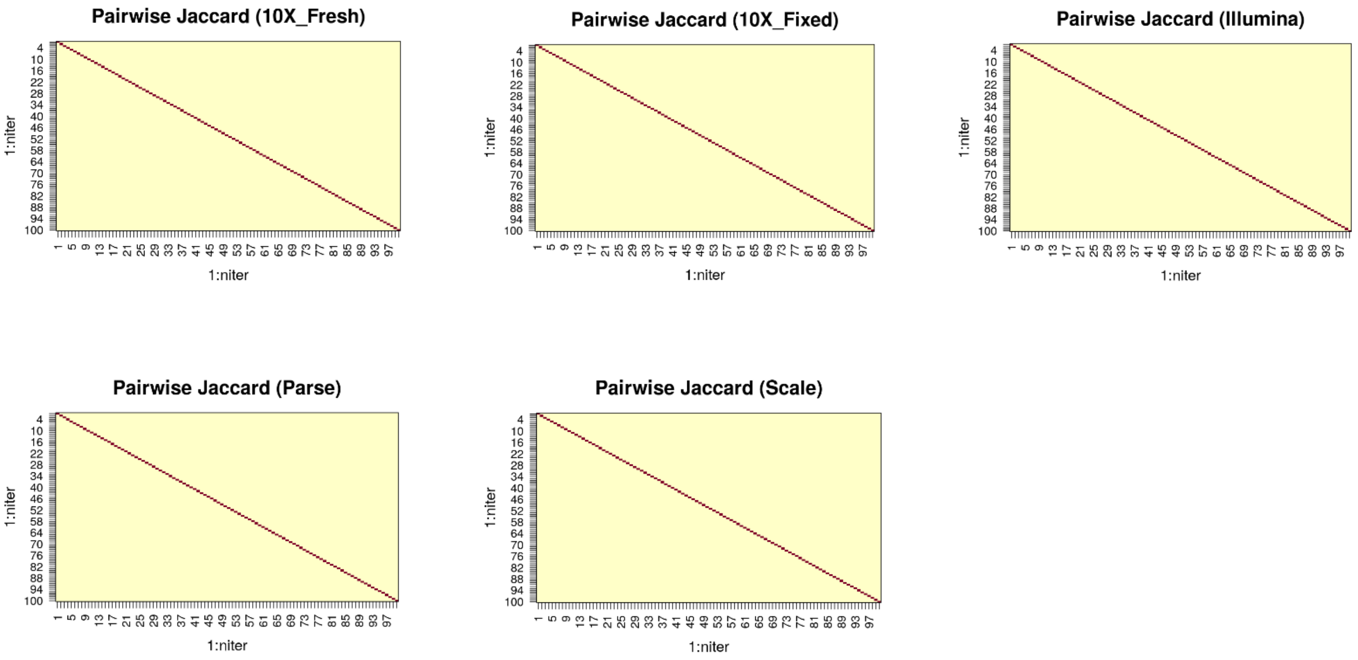


### Agnostic

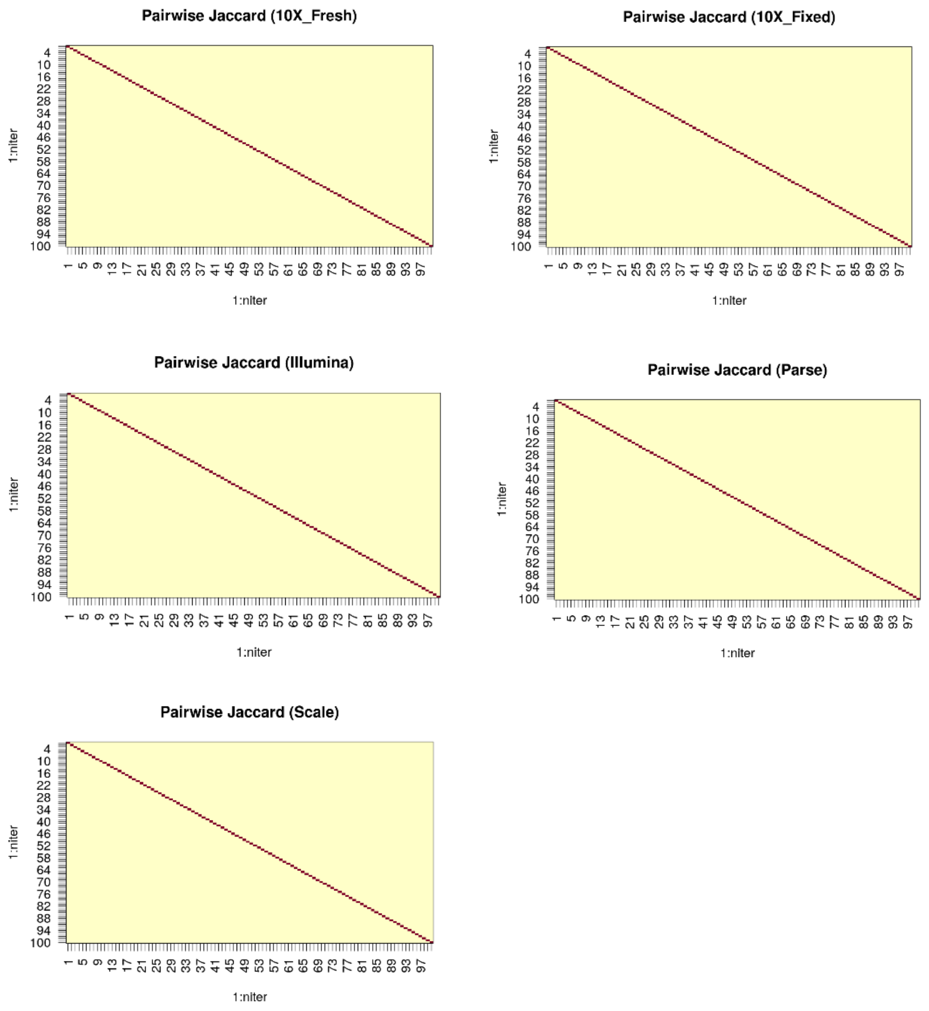


### Platform

Supplemental Figure 2 – Pairwise Jaccard overlap of down sampled cell sets across iterations.
Heatmap showing the Jaccard similarity between the sets of cell barcodes retained in each of *n* independent down sampling iterations for each platform. Each axis indexes iterations (1–100). Values on the diagonal equal 1 (an iteration compared with itself), whereas off-diagonal values quantify how many exact cells are shared between different iterations. Top panel shows agnostic data, while the bottom panel shows platform-specific data.

**Supplemental Figure 3 – Proportion Correlation**

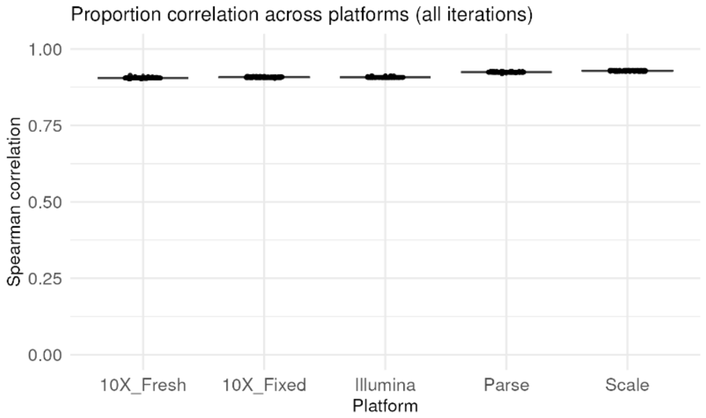


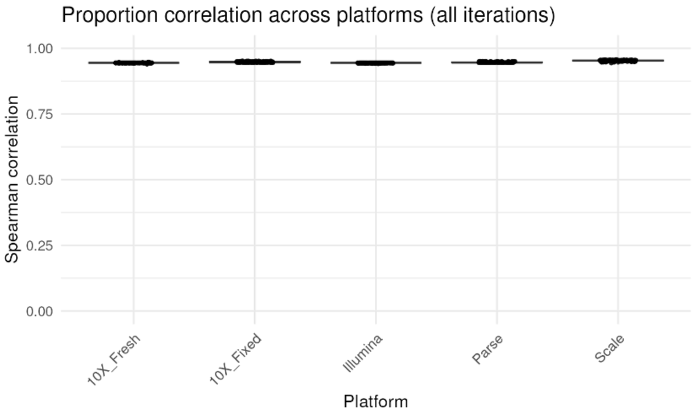


Agnostic

Platform

Supplemental Figure 3 – Proportion correlation across platforms, mapping Spearman correlation for each evaluated platform. Plot on the left is from data obtained using agnostic analysis, plot on right is from data obtained using platform specific analysis.

**Supplementary Figure 4 – Metagene Coverage by Platform**

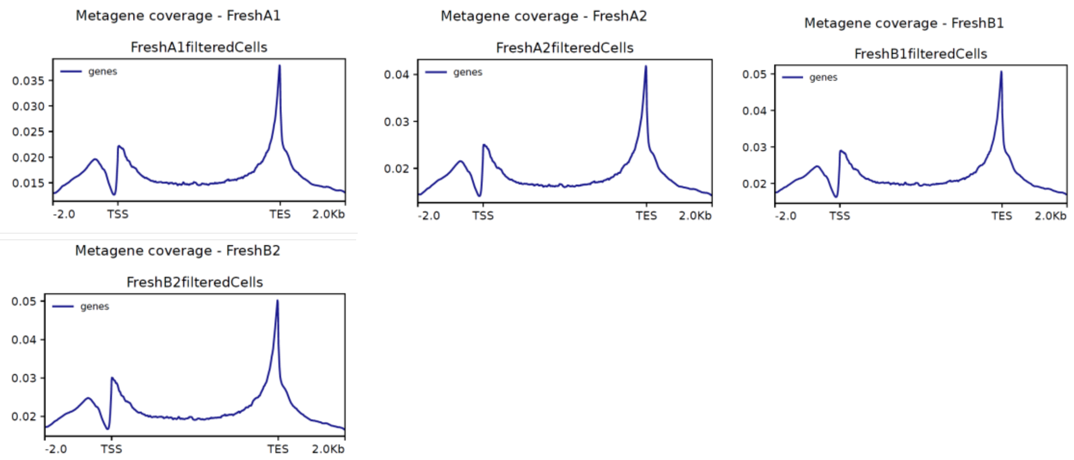


### 10x Fresh

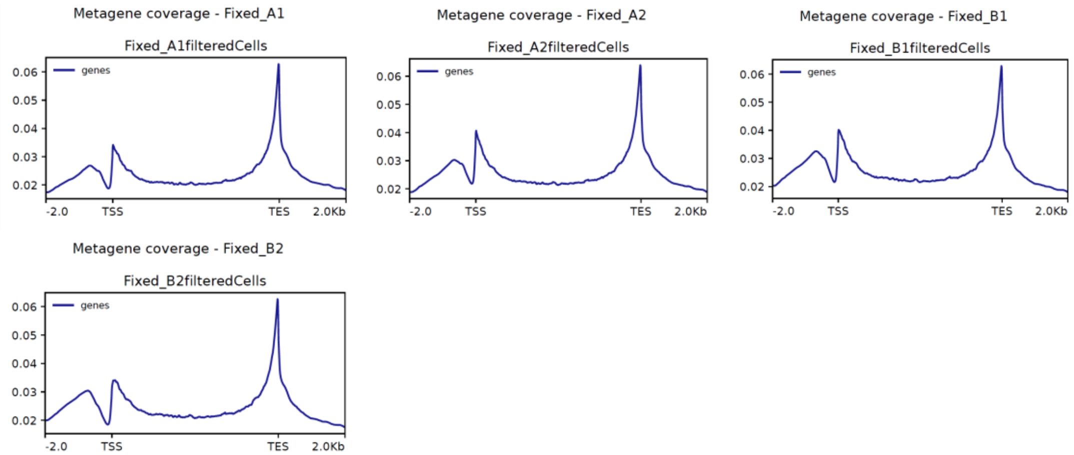


### 10x Fixed

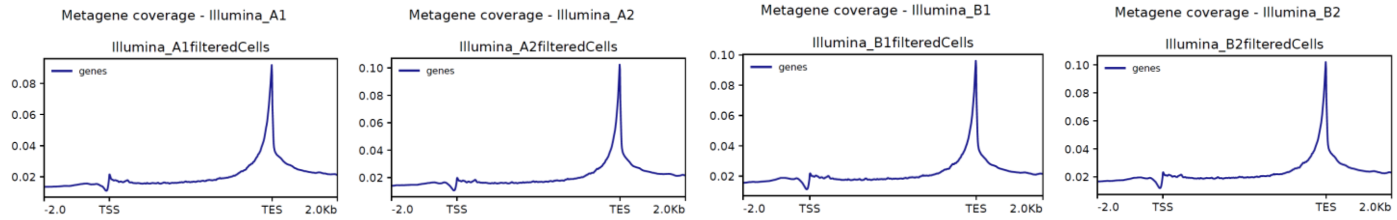


### Illumina

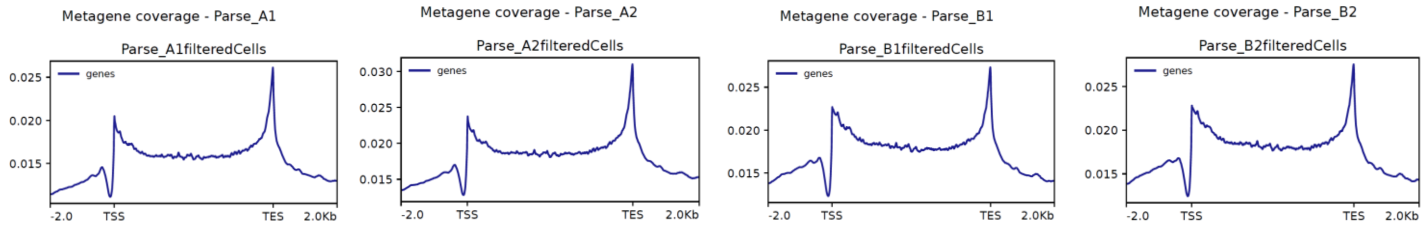


### Parse

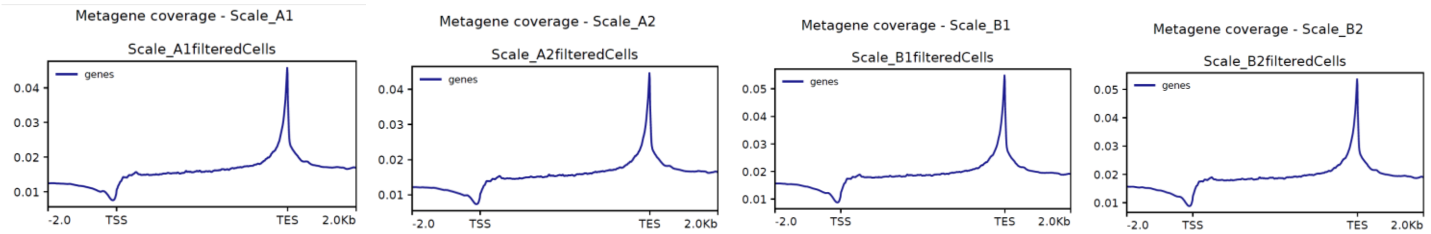


### Scale

Supplementary Figure 4 – Metagene/gene body coverage by platform, mapping the average normalized coverage (in mean Counts per Million/CPM) across metagene position on per-sample basis. All results from platform specific analysis. Transcription end site (TES) represents the 3’ and transcription start site (TSS) represents the 5’ end of the metagene.

**Supplemental Table 4 – Amplification noise/extra-Poisson variation**

| | Agnostic Results | Platform Results |
|---|---|---|
| Platform | mean_extraCV_umis | mean_extraCV_umis |
| 10X_Fresh | 0.87 | 1.13 |
| 10X_Fixed | 0.59 | 0.84 |
| Illumina | -0.19 | 0.76 |
| Parse | 0.2 | 1.14 |
| Scale | 0.44 | 0.01 |

Supplemental Table 4 – Amplification noise/extra-Poisson variability values, mean of samples for each platform.

### Supplemental Table 5 - Ease of Use Analysis

| | 10x Genomics 3' | Illumina | Parse | Scale |
|---|---|---|---|---|
| **Start-up Costs** | $$$$ | $$ | $ | N/A |
| **Clarity of Protocol** | Clear instructions | Clear instructions but discrepencies in dry bath protocols | Clear instructions | Clear instructions |
| **Accessibility of Fixation** | Fixation contains high risk techniques | Simple fixation protocol | Simple fixation protocol | Simple fixation protocol |
| **Sample Types for Input** | Fresh or fixed cells/nuclei as input | Fresh or fixed cells/nuclei as input | Only fixed cells/nuclei as input | Only fixed cells/nuclei as input |
| **Input Options for Fixation** | Can fix 100K-1M cells/nuclei per reaction | Can only fix 1M cells/nuclei per reaction (no lower input option) | Can fix 100K-1M cells/nuclei per reaction | Can fix 100K-1M cells/nuclei per reaction |
| **Storage of Fixed Samples** | Can store fixed samples up to 3 months | Can only store fixed samples for up to 1 wk | Can store fixed samples for up to 6 months | Can store fixed samples for up to 1 year |
| **Range of targeting options** | Can target 500-20,000 cells/sample (500-160,000 cells/run) | Can **input** 100-200,000 cells/sample (100-1M cells/run) | Can target 10,000-100,000 cells/sample (10K-4.8M cells/run) | Can target 1,302-125,000 cells/sample (up to 125,000 cells/run) |
| **High Throughput Targeting Options** | Can target up to 2.56M cells per run with sample multiplexing | Can target up to 19.2M cells per run with sample multiplexing | Can target up 1.92B cells per run with Evercode WT Penta Kit | Can input 168,000-4M cells per sample (168,000-36B cells/run) with Quantum Scale |
| **Throughput (# samples)** | Up to 16 samples per run | Minimum of 2 samples per run | Up to 48 samples per run | Up to 96 samples per run |
| **Hands-on Work** | Less pipetting with lower reaction volumes, limiting reagent costs | Less pipetting with lower reaction volumes, limiting reagent costs | High amount of pipetting at larger reaction volumes, increasing reagent costs | High amount of pipetting at larger reaction volumes, increasing reagent costs |
| **Stop Points (experimental flexibility)** | Increased flexibility with 72hr stop points but less possible stop points than other protocols | Mostly overnight stop points, limiting flexibility | Highest number of stop points but most last only 18 hours, limiting flexibility | Mostly overnight stop points, limiting flexibility |
| **Overnight Holds in Protocol** | No overnight holds in protocol (no UPS required) | Overnight hold on dry bath (or in sterile environment) following PIP creation, necessitating an uninterrupted power source (UPS) | No overnight holds in protocol (no UPS required) | No overnight holds in protocol (no UPS required) |
| **Thermal Cycler Programs** | Manual entry of thermal cycler programs, increasing risk of error | Protocols came pre-loaded on company-provided dry bath, minimizing potential sources of error | Manual entry of thermal cycler programs, increasing risk of error | Manual entry of thermal cycler programs, increasing risk of error |
| **Thermal Cycler Troubleshooting** | Thermal cyclers contain a log, allowing protocols to be verified post-incubation | Dry bath does not contain a log, so run protocols cannot be verified post-incubation | Thermal cyclers contain a log, allowing protocols to be verified post-incubation | Thermal cyclers contain a log, allowing protocols to be verified post-incubation |
| **cDNA Amplification, Quant, and QC** | Good cDNA quant & QC, with clear cDNA amplification cycle selection based on targeted number of cells | Minimal guidance on cDNA amplification cycle selection (only differs between high and low RNA content samples) | No cycling for cDNA amplification but contains cDNA quantification and QC steps | No cycling for cDNA amplification and no cDNA quantification or QC steps |
| **Risk Level of Experiment** | Only use 1/4 of cDNA for library preparation, leaving plenty for re-amplification if needed | Uses all of cDNA for library preparation, meaning experiment must be restarted to re-prepare libraries | All-in approach (failure in protocol causes loss across all samples) | All-in approach (failure in protocol causes loss across all samples) |
| **Total Run Time (no sequencing or analysis)** | 9hrs (fresh) & 10.5hrs (fixed) total run time | 14hrs total run time | 18 hrs total run time | 9 hrs total run time |
| **Final Library Volume** | Yields 35uL of library for sequencing, providing plenty to do multiple sequencing runs | Yields only 20uL of library for sequencing, limiting ability for multiple sequencing runs | Yields only 20uL of library for sequencing, limiting ability for multiple sequencing runs | Yields 30uL of library for sequencing, providing plenty to do multiple sequencing runs |
| **Doublet Detection** | Provides tool for doublet detection | Provides tool for doublet detection | Provides no empirical tool for doublet detection | Provides tool for doublet detection |
| **Miscellaneous Issues** | Frequent clogging in fixed samples in microfluidics system | Wording for dry bath heated lid was confusing on-instrument, increasing potential sources of error | Experienced unexplained failure at cDNA QC stage on first run | Observed an unexplained, unequal distribution of samples |

Supplemental Table 5 – Ease of use analysis including details on each platform and their protocol benefits/limitations.

### Supplemental Table 6 – Cost Analysis

| Company | Upfront equipment to purchase | Upfront equipment cost | Annual equipment service agreement |
|---|---|---|---|
| Parse | Parse Magnetic Rack, PN SB1004 | $ 300.00 | n/a |
| Scale | n/a | n/a | n/a |
| Illumina | Illumina Single Cell Prep Starter Equipment, PN 20132796 | $ 3,500.00 | n/a |
| 10x Genomics | Chromium Xo, iX, or X | $25,000 - $65,000 | $5,000 - $12,000 |

| Company | Kit Name | Kit PN | Kit Price | kit size | # of cells that can be fixed per rxn | per rxn | Fixation 3rd party consumables, per rxn | Total, per rxn |
|---|---|---|---|---|---|---|---|---|
| Parse | Evercode™ Cell Fixation v3, 12 rxn | ECFC3300 | $ 480.00 | 12 rxns | 1,000,000.00 | $ 40.00 | $ 17.34 | $ 57.34 |
| Scale | ScaleBio Sample Fixation Kit | 2020001 | $ 300.00 | 12 rxns | 1,000,000.00 | $ 25.00 | $ 5.21 | $ 30.21 |
| Illumina | n/a | n/a | n/a | n/a | 1,000,000.00 | n/a | $ 23.16 | $ 23.16 |
| 10x Genomics, Fixed | n/a | n/a | n/a | n/a | 1,000,000.00 | n/a | $ 10.09 | $ 10.09 |

| | | | | | | | | | | With Labor | | | Without Labor | | |
|---|---|---|---|---|---|---|---|---|---|---|---|---|---|---|---|
| Company | Kit Name | Kit PN | Kit Price | kit size | library prep 3rd party consumables, per 4 samples | labor (h) | labor ($) | sequencing (NovaSeq X) | Sequencing cost | TOTAL | per cell (40k cells) | per cell, maximize kit | TOTAL | per cell (40k cells) | per cell, maximize kit |
| Parse | Evercode™ WT v3 | ECWT3300 | $ 10,780.00 | 100k cells / 48 samples | $ 538.86 | 18.30 | $ 439.20 | 1 lane NovaSeq X 10B, 200 cycle | $ 1,158.75 | $ 12,916.81 | $ 0.32 | $ 0.15 | $ 12,477.61 | $ 0.31 | $ 0.15 |
| Parse | WT UDI plate | | $ 385.00 | 48 rxns | | | | | | | $ 0.0008 | $ 0.0002 | | $0.0004 | $ 0.0002 |
| Scale | ScaleBio Single Cell RNA Sequencing Kit v1.1 | 950884 | $ 8,500.00 | 125k cells | $ 439.38 | 9.20 | $ 220.80 | 1 lane NovaSeq X 10B, 100 cycle | $ 915.00 | $ 10,075.18 | $ 0.25 | $ 0.12 | $ 9,854.38 | $ 0.25 | $ 0.12 |
| Illumina | Illumina Single Cell 3' RNA Prep, T10 | 20135691 | $ 4,800.00 | 8 rxns x 10k cells ea (80k) | $ 245.19 | 14.20 | $ 340.80 | 1 lane NovaSeq X 10B, 100 cycle | $ 915.00 | $ 6,300.99 | $ 0.16 | $ 0.09 | $ 5,960.19 | $ 0.15 | $ 0.09 |
| 10x Genomics, Fixed | Chromium GEM-X Single Cell 3' Kit v4, 4 rxns | 1000686 | $ 6,498.00 | 4 rxns x 20k cells ea (80k) | $ 279.85 | 10.50 | $ 252.00 | 1 lane NovaSeq X 10B, 100 cycle | $ 915.00 | $ 7,944.85 | $ 0.20 | $ 0.11 | $ 7,692.85 | $ 0.19 | $ 0.11 |
| 10x Genomics, fresh | Chromium GEM-X Single Cell 3' Kit v4, 4 rxns | 1000686 | $ 6,498.00 | 4 rxns x 20k cells ea (80k) | $ 279.85 | 9.00 | $ 216.00 | 1 lane NovaSeq X 10B, 100 cycle | $ 915.00 | $ 7,908.85 | $ 0.20 | $ 0.11 | $ 7,692.85 | $ 0.19 | $ 0.11 |
| 10x Genomics | Dual Index Kit TT Set A | 1000215 | $ 960.00 | 96 rxns | | | | | | | $ 0.001 | $ 0.001 | | $ 0.001 | $ 0.001 |

Notes
List prices
ABRF labor rate
3rd party consumables that are recommended in each protocol
sequencing - flow cell cost only

Supplemental Table 6 – Detailed cost analysis with breakdown of each value that contributes to overall cost of each scRNAseq assay.

**Supplemental Table 7 – RNA QC**

| RNA QC | | | |
|---|---|---|---|
| | RNA Extraction Method Used | Any Adjustments to RNA Extraction Protocol | RNA QC Method |
| 10x Fresh | 'Direct-zol RNA Microprep Kit with TRI Reagent' (CAT #R2061-A, Zymo Research Corporation, Irvine, CA) | N/A | Agilent 4200 TapeStation System & High Sensitivity RNA ScreenTape assay consumables (PN G2991BA, PN 5067-5580, PN 5067-5581 & PN 5067-5579, Agilent Technologies, Santa Clara, CA) |
| 10x Fixed | 'Quick-RNA™ FFPE Kit' (CAT #R1008, Zymo Research Corporation, Irvine, CA) | skipped deparaffinization - started from the tissue digestion step | Agilent 4200 TapeStation System & High Sensitivity RNA ScreenTape assay consumables (PN G2991BA, PN 5067-5580, PN 5067-5581 & PN 5067-5579, Agilent Technologies, Santa Clara, CA) |
| Illumina | 'Quick-RNA™ FFPE Kit' (CAT #R1008, Zymo Research Corporation, Irvine, CA) | skipped deparaffinization - started from the tissue digestion step | Agilent 4200 TapeStation System & High Sensitivity RNA ScreenTape assay consumables (PN G2991BA, PN 5067-5580, PN 5067-5581 & PN 5067-5579, Agilent Technologies, Santa Clara, CA) |
| Parse | 'Quick-RNA™ FFPE Kit' (CAT #R1008, Zymo Research Corporation, Irvine, CA) | skipped deparaffinization - started from the tissue digestion step | Agilent 4200 TapeStation System & High Sensitivity RNA ScreenTape assay consumables (PN G2991BA, PN 5067-5580, PN 5067-5581 & PN 5067-5579, Agilent Technologies, Santa Clara, CA) |
| Scale | 'Quick-RNA™ FFPE Kit' (CAT #R1008, Zymo Research Corporation, Irvine, CA) | skipped deparaffinization - started from the tissue digestion step | Agilent 4200 TapeStation System & High Sensitivity RNA ScreenTape assay consumables (PN G2991BA, PN 5067-5580, PN 5067-5581 & PN 5067-5579, Agilent Technologies, Santa Clara, CA) |

Supplemental Table 7 – RNA QC details, including kits used to isolate RNA, any protocol adjustments, and the method of RNA QC. Color-coded by platform.

#### Supplemental Table 8 - Workflow Timeline per Day

**Workflow Timeline per Day (minutes)**

| 10x Fresh | | DAY 1 | DAY 2 | DAY 3 | DAY 4 | DAY 5 | | | TOTAL |
|---|---|---|---|---|---|---|---|---|---|
| Sample Preparation | Hands-on Time | 45 | | | | | | | |
| Single Cell Capture & RT | Hands-on Time | 40 | | | | | | | |
| | On-instrument | 55 | | | | | | | |
| cDNA Amplification | On-instrument | | 40 | | | | | | |
| | Hands-on Time | | 95 | | | | | | |
| | QC | | 30 | | | | | | |
| Gene Expression Library Prep | Hands-on Time | | | 110 | | | | | |
| | On-instrument | | | 95 | | | | | |
| | QC | | | 30 | | | | | |
| Library Quantification | Hands-on Time | | | | 60 | | | | |
| | On-instrument | | | | 90 | | | | |
| Sequencing | Hands-on Time | | | | | 90 | | | |
| | On-instrument | | | | | 1275 | | | |
| Cell Count & QC | Run Time | | | | | | 473 | | |
| Total Time | Hands-on Time | | | | | | | 440 | |
| Total Time | On-instrument | | | | | | | 1615 | |
| Total Run Time (without analysis) | | | | | | | | | 34hrs 15mins |
| Total Run Time (with analysis) | | | | | | | | | 42hrs 6mins |

| 10x Fixed | | DAY 1 | DAY 2 | DAY 3 | DAY 4 | DAY 5 | | | TOTAL |
|---|---|---|---|---|---|---|---|---|---|
| Sample Preparation | Hands-on Time | 45 | | | | | | | |
| Fixation | Hands-on Time | 90 | | | | | | | |
| Single Cell Capture & RT | Hands-on Time | 40 | | | | | | | |
| | On-instrument | 55 | | | | | | | |
| cDNA Amplification | On-instrument | | 40 | | | | | | |
| | Hands-on Time | | 95 | | | | | | |
| | QC | | 30 | | | | | | |
| Gene Expression Library Prep | Hands-on Time | | | 110 | | | | | |
| | On-instrument | | | 95 | | | | | |
| | QC | | | 30 | | | | | |
| Library Quantification | Hands-on Time | | | | 60 | | | | |
| | On-instrument | | | | 90 | | | | |
| Sequencing | Hands-on Time | | | | | 90 | | | |
| | On-instrument | | | | | 1275 | | | |
| Cell Count & QC | Run Time | | | | | | 476 | | |
| Total Time | Hands-on Time | | | | | | | 530 | |
| Total Time | On-instrument | | | | | | | 1615 | |
| Total Run Time (without analysis) | | | | | | | | | 35hrs 45mins |
| Total Run Time (with analysis) | | | | | | | | | 43hrs 41mins |

| Illumina | | DAY 1 | DAY 2 | DAY 3 | DAY 4 | DAY 5 | | | TOTAL |
|---|---|---|---|---|---|---|---|---|---|
| Sample Preparation | Hands-on Time | 45 | | | | | | | |
| Fixation | Hands-on Time | 60 | | | | | | | |
| Single Cell Capture & RT | Hands-on Time | 68 | | | | | | | |
| | On-instrument | 295 | | | | | | | |
| cDNA Amplification | On-instrument | | 80 | | | | | | |
| | Hands-on Time | | 132 | | | | | | |
| | QC | | 20 | | | | | | |
| Gene Expression Library Prep | Hands-on Time | | | 86 | | | | | |
| | On-instrument | | | 40 | | | | | |
| | QC | | | 25 | | | | | |
| Library Quantification | Hands-on Time | | | | 60 | | | | |
| | On-instrument | | | | 90 | | | | |
| Sequencing | Hands-on Time | | | | | 90 | | | |
| | On-instrument | | | | | 1275 | | | |
| Cell Count & QC | Run Time | | | | | | 88 | | |
| Total Time | Hands-on Time | | | | | | | 541 | |
| Total Time | On-instrument | | | | | | | 1825 | |
| Total Run Time (without analysis) | | | | | | | | | 39hrs 26mins |
| Total Run Time (with analysis) | | | | | | | | | 40hrs 54mins |

| Parse | | DAY 1 | DAY 2 | DAY 3 | DAY 4 | DAY 5 | | | TOTAL |
|---|---|---|---|---|---|---|---|---|---|
| Sample Preparation | Hands-on Time | 45 | | | | | | | |
| Fixation | Hands-on Time | 60 | | | | | | | |
| Single Cell Capture & RT | Hands-on Time | 224 | | | | | | | |
| | On-instrument | 75 | | | | | | | |
| cDNA Amplification | On-instrument | | 195 | | | | | | |
| | Hands-on Time | | 135 | | | | | | |
| | QC | | 30 | | | | | | |
| Gene Expression Library Prep | Hands-on Time | | | 215 | | | | | |
| | On-instrument | | | 85 | | | | | |
| | QC | | | 30 | | | | | |
| Library Quantification | Hands-on Time | | | | 60 | | | | |
| | On-instrument | | | | 90 | | | | |
| Sequencing | Hands-on Time | | | | | 90 | | | |
| | On-instrument | | | | | 1275 | | | |
| Cell Count & QC | Run Time | | | | | | 1064 | | |
| Total Time | Hands-on Time | | | | | | | 829 | |
| Total Time | On-instrument | | | | | | | 1780 | |
| Total Run Time (without analysis) | | | | | | | | | 43hrs 29mins |
| Total Run Time (with analysis) | | | | | | | | | 61hrs 13mins |

| Scale | | DAY 1 | DAY 2 | DAY 3 | DAY 4 | DAY 5 | | | TOTAL |
|---|---|---|---|---|---|---|---|---|---|
| Sample Preparation | Hands-on Time | 45 | | | | | | | |
| Fixation | Hands-on Time | 45 | | | | | | | |
| Single Cell Capture & RT | Hands-on Time | 45 | | | | | | | |
| | On-instrument | 22 | | | | | | | |
| cDNA Amplification | On-instrument | | 140 | | | | | | |
| | Hands-on Time | | 40 | | | | | | |
| | QC | | 20 | | | | | | |
| Gene Expression Library Prep | Hands-on Time | | | 96 | | | | | |
| | On-instrument | | | 64 | | | | | |
| | QC | | | 30 | | | | | |
| Library Quantification | Hands-on Time | | | | 60 | | | | |
| | On-instrument | | | | 90 | | | | |
| Sequencing | Hands-on Time | | | | | 40 | | | |
| | On-instrument | | | | | 1155 | | | |
| Cell Count & QC | Run Time | | | | | | 830 | | |
| Total Time | Hands-on Time | | | | | | | 371 | |
| Total Time | On-instrument | | | | | | | 1521 | |
| Total Run Time (without analysis) | | | | | | | | | 31hrs 32mins |
| Total Run Time (with analysis) | | | | | | | | | 45hrs 22mins |

Supplemental Table 8 – Workflow timeline per day (in minutes), demonstrating a detailed breakdown of hands-on vs. on-instrument time at each step of the scRNAseq protocols, including the time required for basic analysis. Summative value is provided in hours and minutes. Color-coded by platform.

**Supplemental Figure 5 – Mitochondrial RNA content per sample**

### Platform Pre-QC

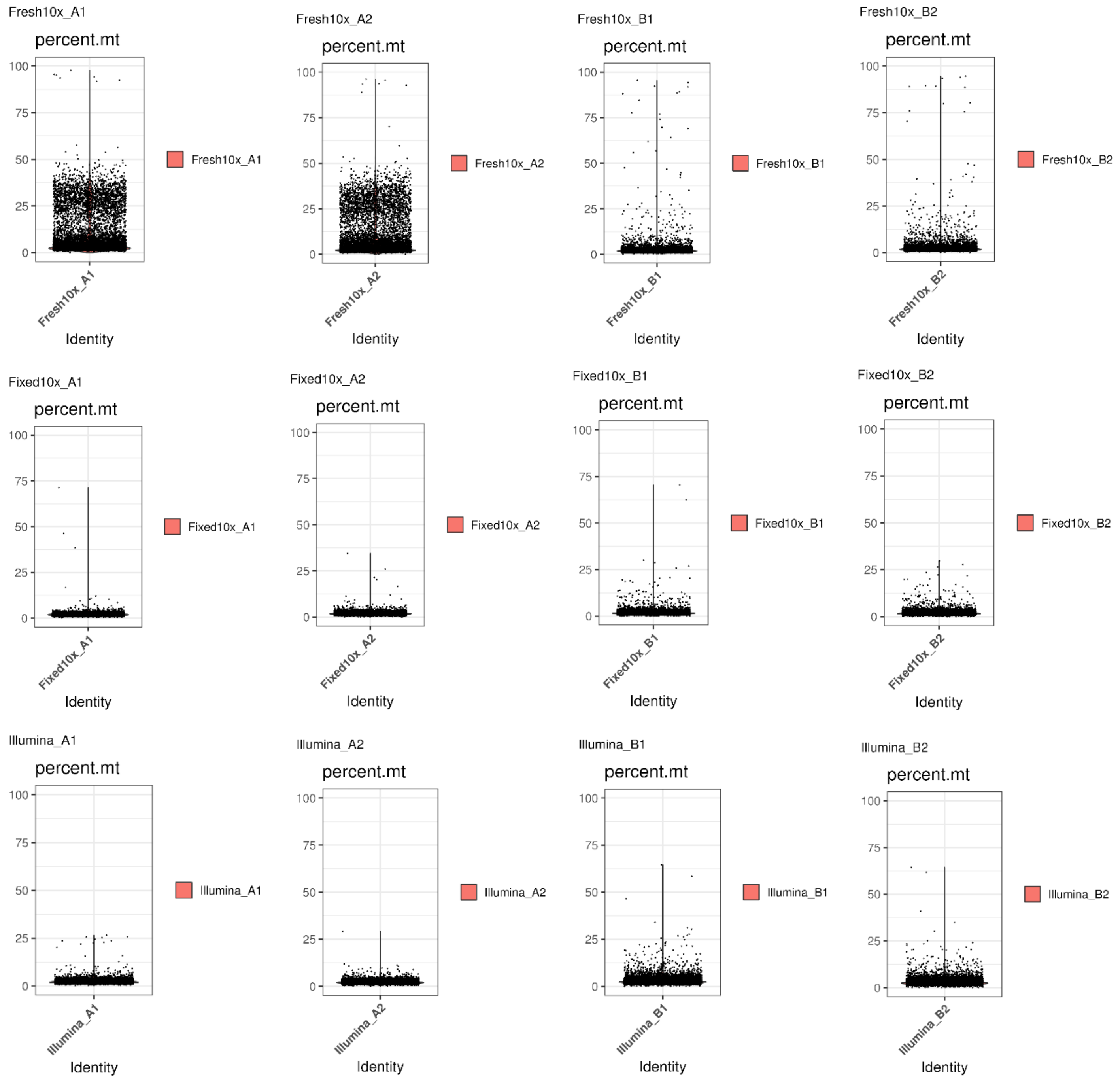

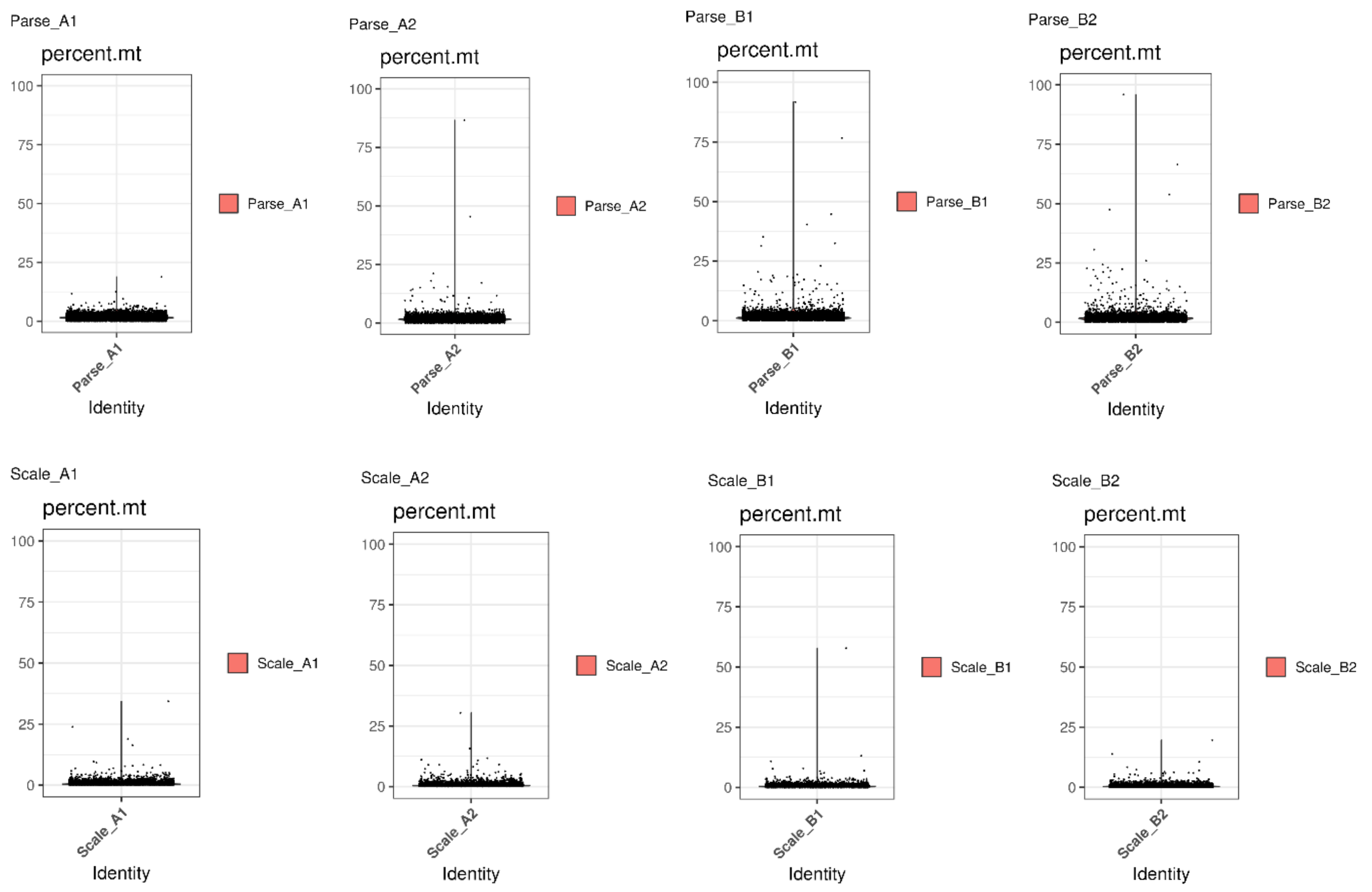


### Platform post-QC

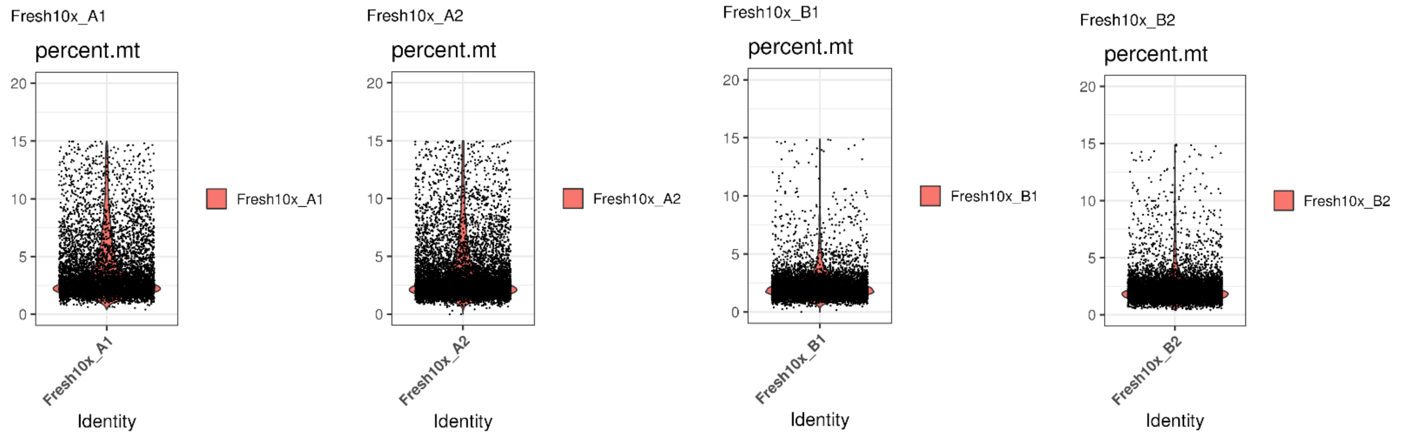

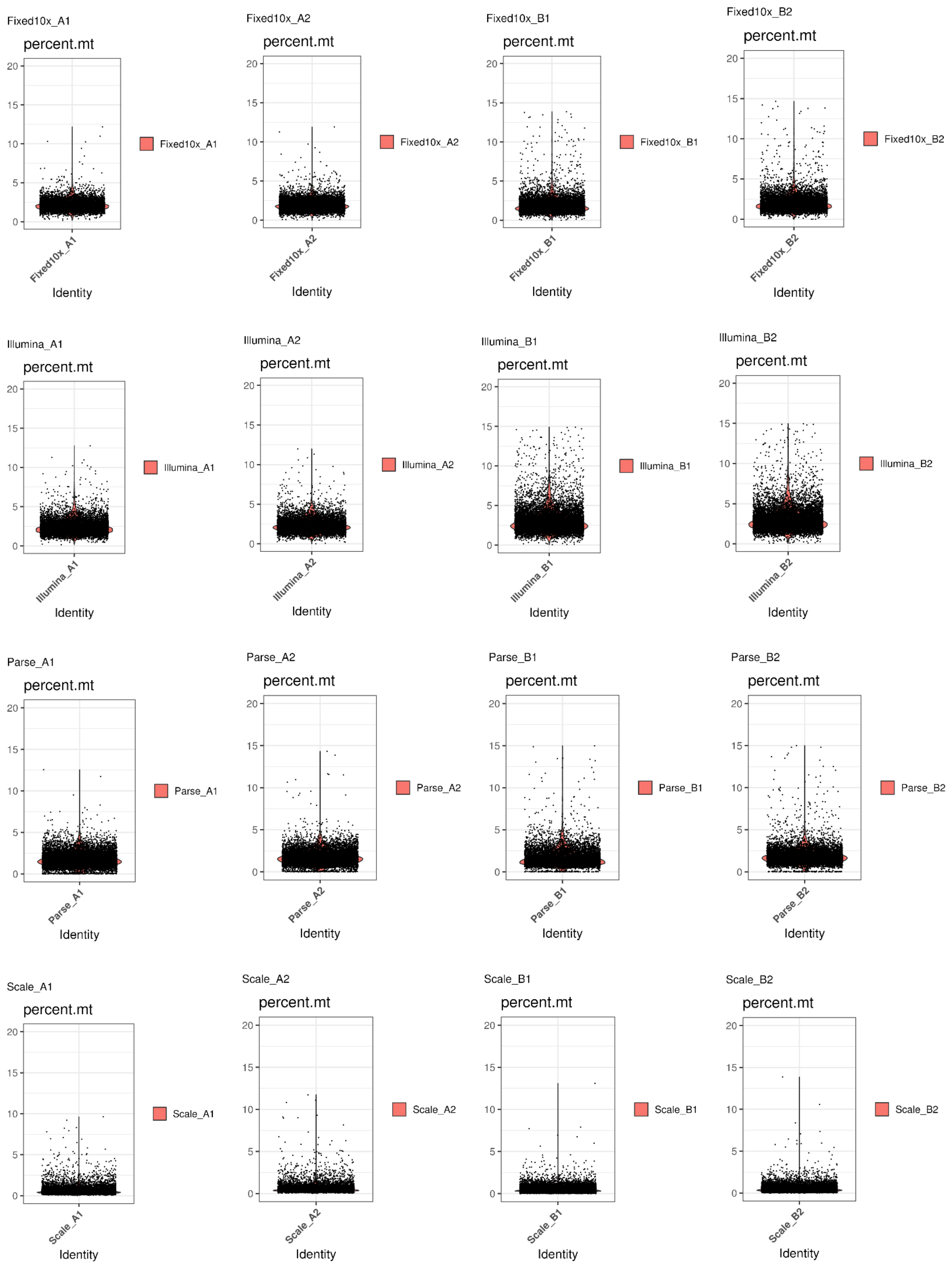

Fixed10x_A1
percent.mt
Fixed10x_A1
Identity
Fixed10x_A2
percent.mt
Fixed10x_A2
Identity
Fixed10x_B1
percent.mt
Fixed10x_B1
Identity
Fixed10x_B2
percent.mt
Fixed10x_B2
Identity
Illumina_A1
percent.mt
Illumina_A1
Identity
Illumina_A2
percent.mt
Illumina_A2
Identity
Illumina_B1
percent.mt
Illumina_B1
Identity
Illumina_B2
percent.mt
Illumina_B2
Identity
Parse_A1
percent.mt
Parse_A1
Identity
Parse_A2
percent.mt
Parse_A2
Identity
Parse_B1
percent.mt
Parse_B1
Identity
Parse_B2
percent.mt
Parse_B2
Identity
Scale_A1
percent.mt
Scale_A1
Identity
Scale_A2
percent.mt
Scale_A2
Identity
Scale_B1
percent.mt
Scale_B1
Identity
Scale_B2
percent.mt
Scale_B2
Identity
0
5
10
15
20

Supplemental Figure 5 – Percentage of mitochondrial RNA content per sample, modelled for platform-specific analysis pre- and post-QC. Threshold was set to 15% mitochondrial content post-QC.

#### Supplemental Figure 6 – Scale Experimental Worksheet

| | 2 | 3 | 4 | 5 | 6 | 7 | 8 | 9 | 10 | 11 | 12 |
|---|---|---|---|---|---|---|---|---|---|---|---|
| A1 | A1 | A2 | A2 | A2 | B1 | B1 | B1 | B2 | B2 | B2 | |
| A1 | A1 | A2 | A2 | A2 | B1 | B1 | B1 | B2 | B2 | B2 | |
| A1 | A1 | A2 | A2 | A2 | B1 | B1 | B1 | B2 | B2 | B2 | |
| A1 | A1 | A2 | A2 | A2 | B1 | B1 | B1 | B2 | B2 | B2 | |
| A1 | A1 | A2 | A2 | A2 | B1 | B1 | B1 | B2 | B2 | B2 | |
| A1 | A1 | A2 | A2 | A2 | B1 | B1 | B1 | B2 | B2 | B2 | |
| A1 | A1 | A2 | A2 | A2 | B1 | B1 | B1 | B2 | B2 | B2 | |
| A1 | A1 | A2 | A2 | A2 | B1 | B1 | B1 | B2 | B2 | B2 | |

Cell Sequencing (Number of cells you plan to target in final library, MAX 125,000)

| Sample | No. Wells | Cells per Wel | Cell Loading | Estimated Sequenced | | |
|---|---|---|---|---|---|---|
| A1 | 24 | 5700 | 136800 | 31250 | Requires User Input | |
| A2 | 24 | 5700 | 136800 | 31250 | | |
| B1 | 24 | 5700 | 136800 | 31250 | | |
| B2 | 24 | 5700 | 136800 | 31250 | Total Loading | 547200 |

Supplemental Figure 6 – Scale experimental worksheet with detailed sample loading layout and total number of cells loaded per well into the Scale scRNAseq assay.

#### Supplemental Figure 7 – Parse Experimental Worksheet

Parse Biosciences
700 Dexter Ave
Suite 600
Seattle, WA 98109

PARSE BIOSCIENCES

Support Suite: support.parsebiosciences.com

WT - Version 2.0

##### Evercode WT Sample Loading Table

- For more details on using this Sample Loading Table, see the Important Guidelines section of the User Manual

**This sheet should be filled out prior to starting Section 1.**

| Step | Instructions |
|---|---|
| 1 | Ensure Macros are enabled. |
| 2 | Input the number of samples. |
| 3 | Input the target number of barcoded cells. Note: The default is 100,000 cells for Evercode WT. |
| 4 | Input your sample names. |
| 5 | Input the target percentage representation of each sample in the final library. CRITICAL: No percentage can be lower than 2.09%. |
| | If not already done, count the samples as described in Section 1.1 of the Evercode WT User Manual. |
| 6 | Input stock cell concentration for each sample. |
| 7 | Prepare the dilutions as described. CRITICAL: Ensure that Sample Dilution Buffer is completely thawed before use. |
| 8 | Open the "Plate Configuration" sheet. With the plate on ice, add 14 uL of each diluted sample to the appropriate well(s) of the Round 1 Plate as shown in the plate map. CRITICAL: Follow the instructions in the User Guide with respect to sample mixing and changing tips. |

CRITICAL: We do not recommend editing cells highlighted in grey.

**Number of Samples (Step 2):** 4
**Target Number Barcoded Cells (Step 3):** 170,000

| Sample # | Sample Name (Step 4) | Percent of Library (Step 5) | Stock Concentration (cells/uL) (Step 6) | Number of Wells | Targeted Number of Barcoded Cells | Required Sample Concentration (cells/uL) | Required Volume + 10% overage (uL) | Volume of Sample Stock Dilution (uL) (Step 7) | Volume of Sample Dilution Buffer (uL) (Step 7) |
|---|---|---|---|---|---|---|---|---|---|
| 1 | Parse_A1_Run2 | 25.00% | 3,840 | 12 | 42500 | 884 | 185 | 42.57 | 142.23 |
| 2 | Parse_A2_Run2 | 25.00% | 3,840 | 12 | 42500 | 884 | 185 | 42.57 | 142.23 |
| 3 | Parse_B1_Run2 | 25.00% | 3,600 | 12 | 42500 | 884 | 185 | 45.40 | 139.40 |
| 4 | Parse_B2_Run2 | 25.00% | 3,600 | 12 | 42500 | 884 | 185 | 45.40 | 139.40 |
| TOTALS: | | 100% | | 48 | 170,000 | | | | |

Supplemental Figure 7 – Parse experimental worksheet, demonstrating library distribution, input concentration, targeted number of barcoded cells, required volumes and concentrations for sample loading.

**Supplemental Figure 8 – Library Traces**

### 10x Fresh A1

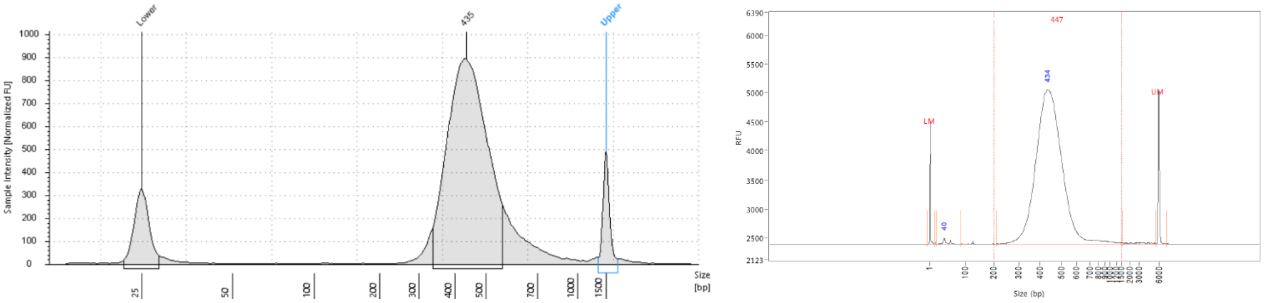


### 10x Fresh A2

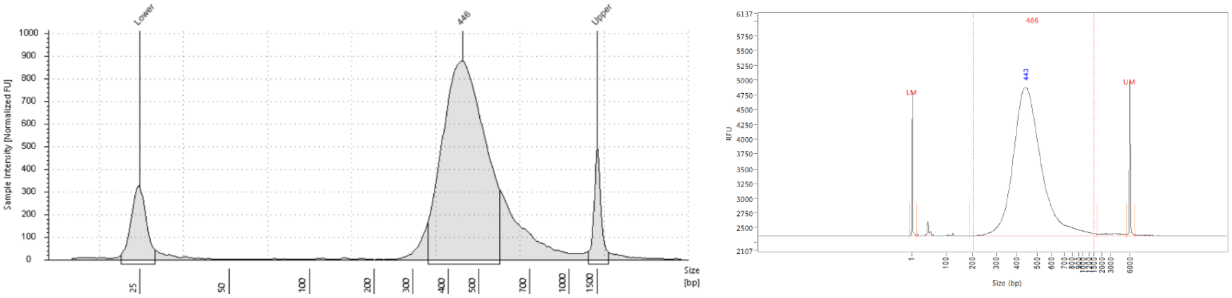


### 10x Fresh B1

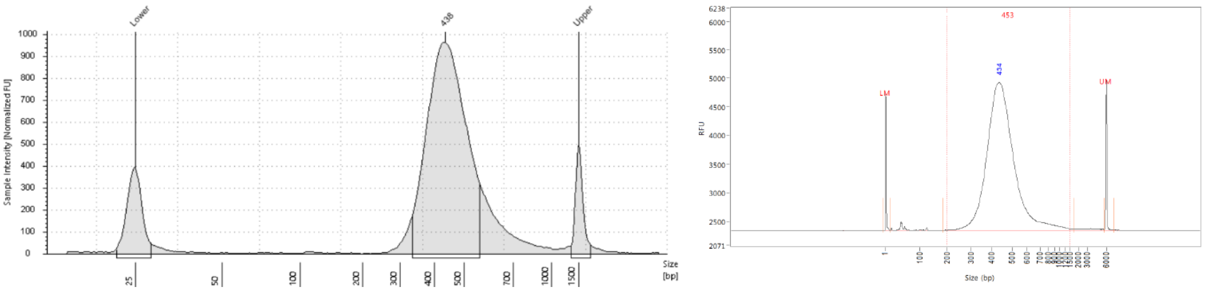


### 10x Fresh B2

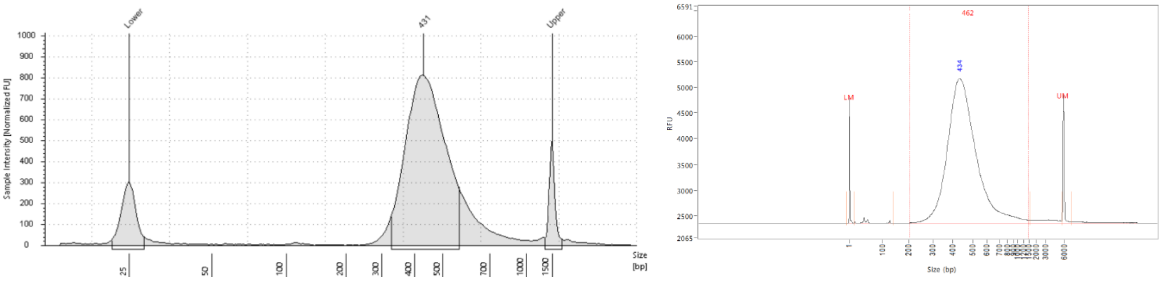


### 10x Fixed A1

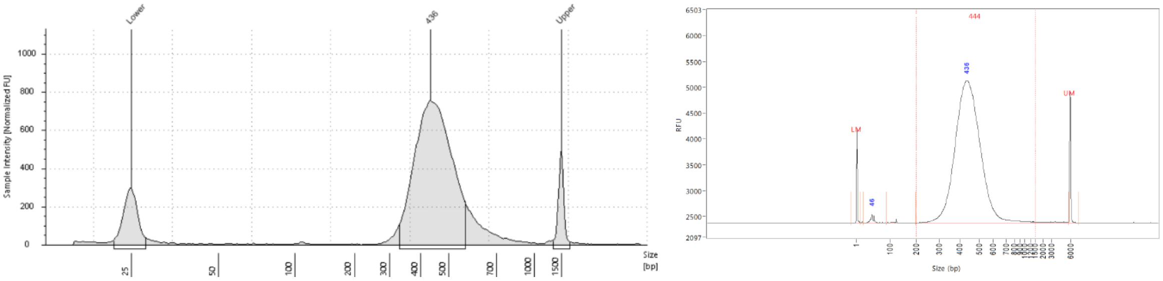


10x Fixed A2

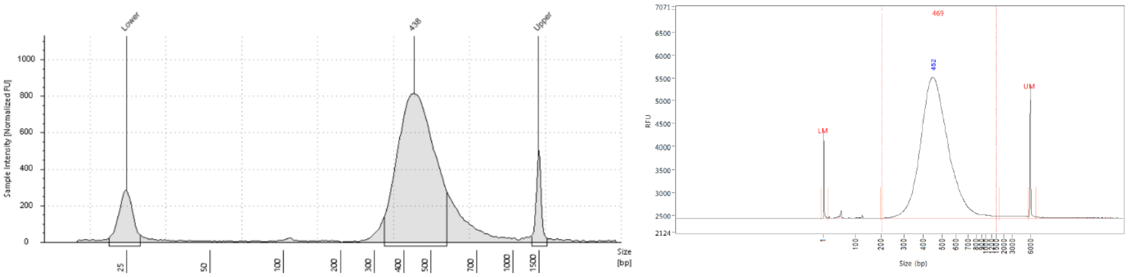


10x Fixed B1

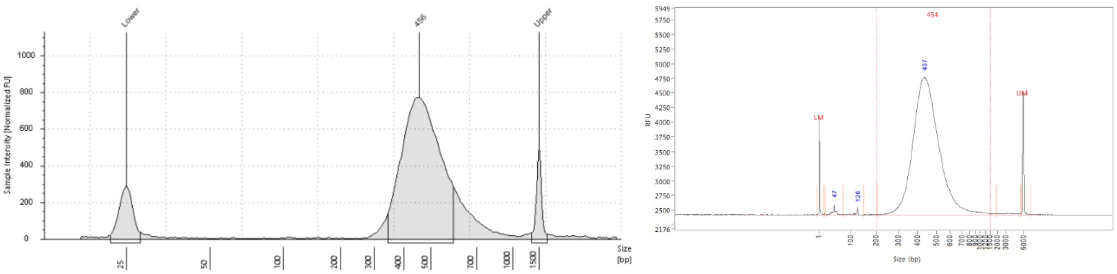


10x Fixed B2

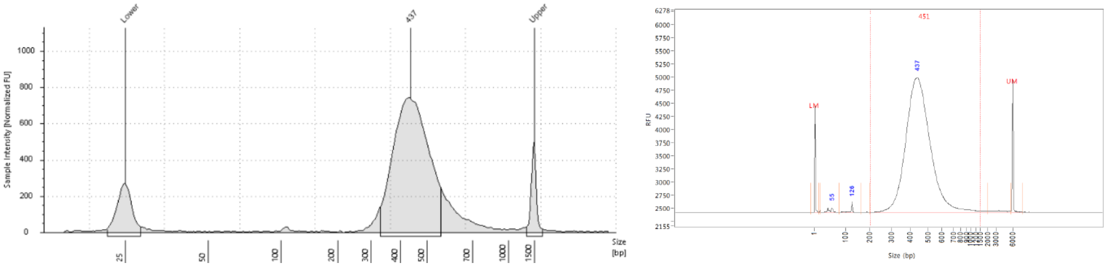


Illumina A1

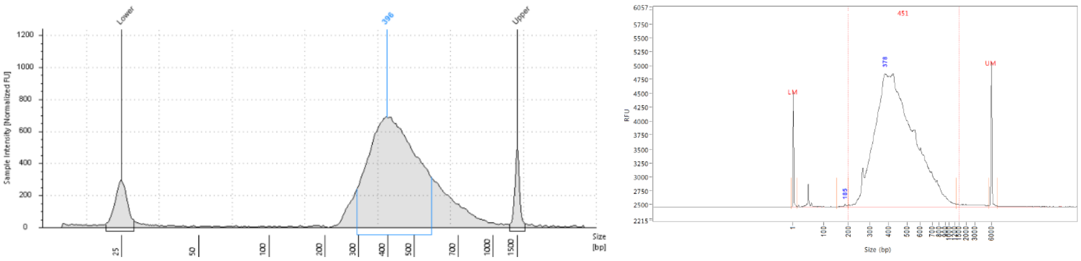


Illumina A2

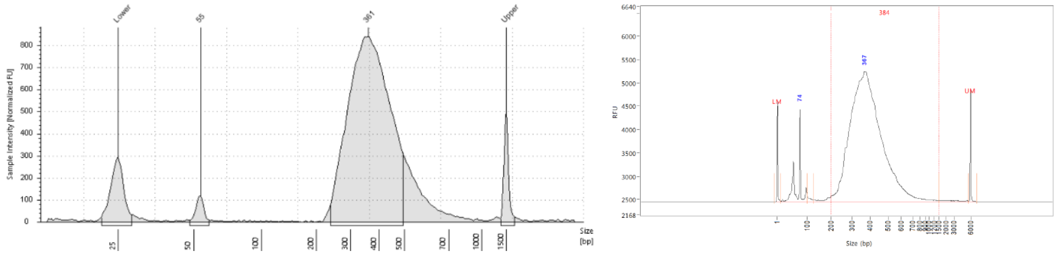


Illumina B1

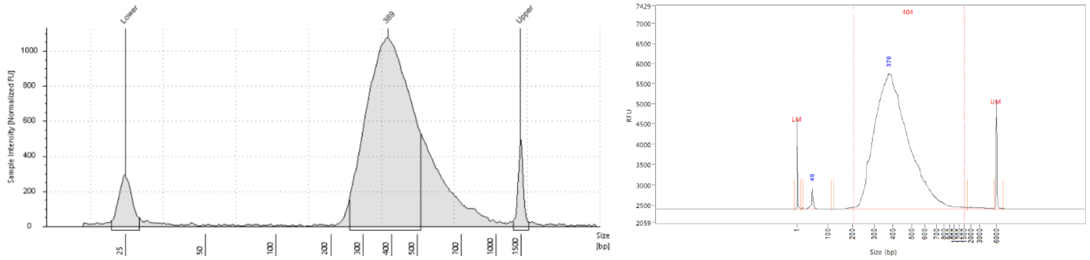


Illumina B2

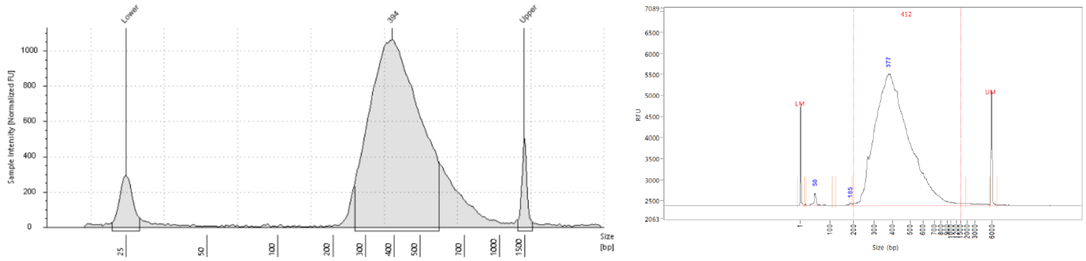


Parse A1

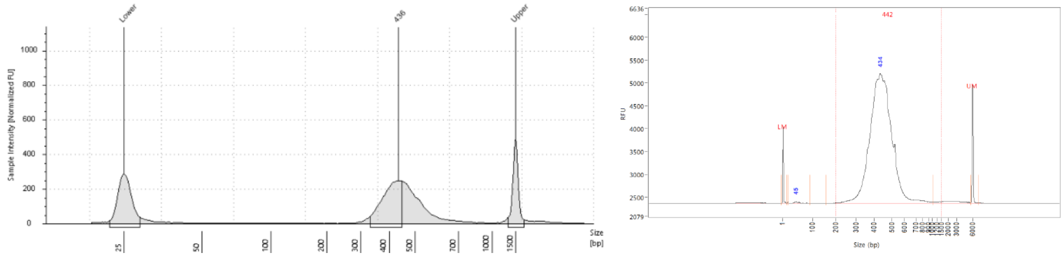


Parse A2

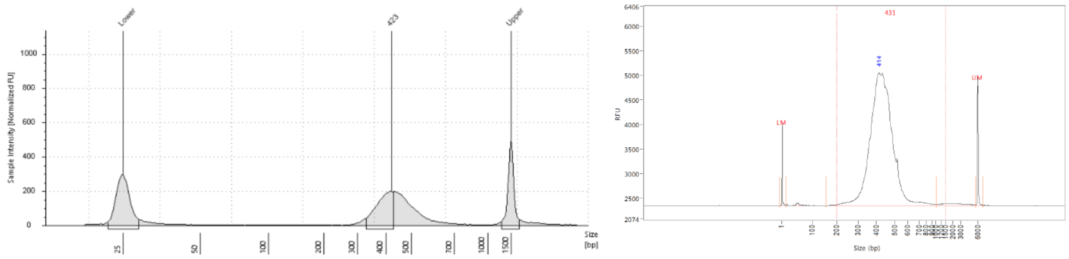


Parse B1

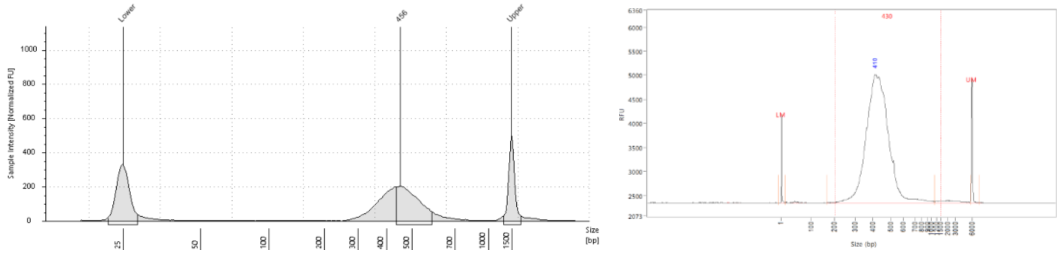


### Parse B2

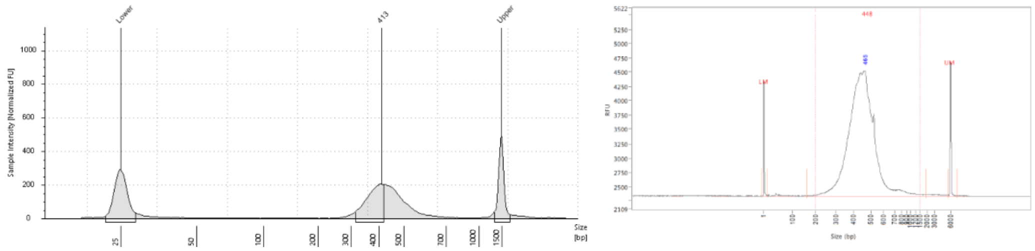


### Scale Initial Sequencing Run

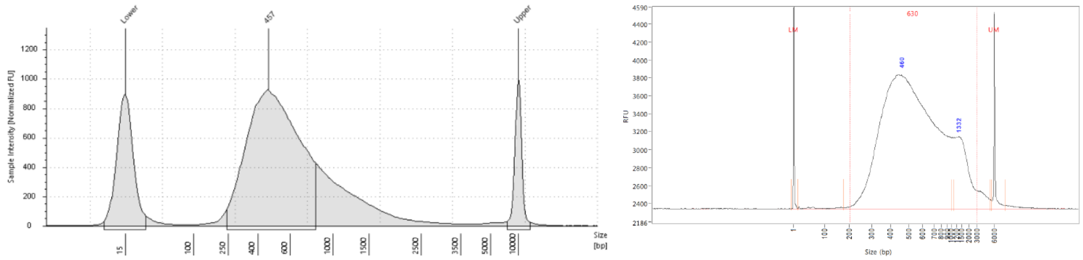


### Scale Re-sequencing Run Pool 2

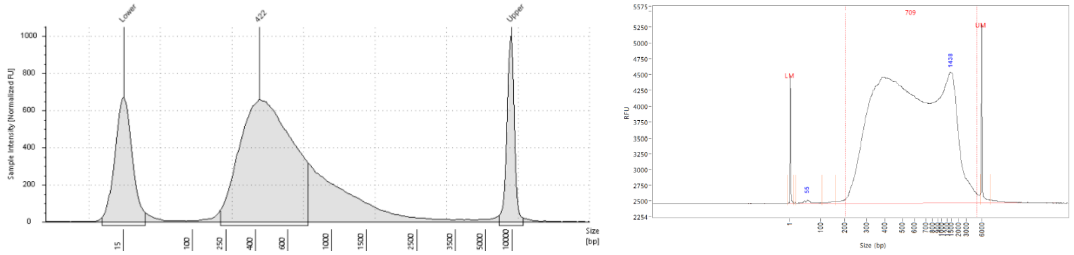


### Scale Re-sequencing Run Pool 3

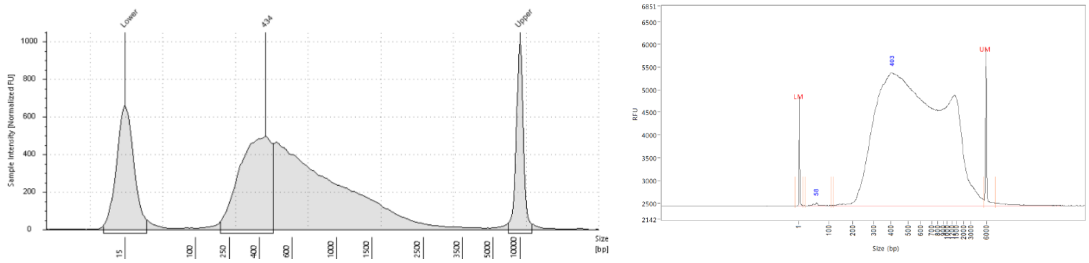


Supplemental Figure 8 - Library traces from the 4200 TapeStation (left) and Bioanalyzer (right), demonstrating the size of final libraries.

**Supplemental Figure 9 – Pass/Fail Sequencing Filtration**

A

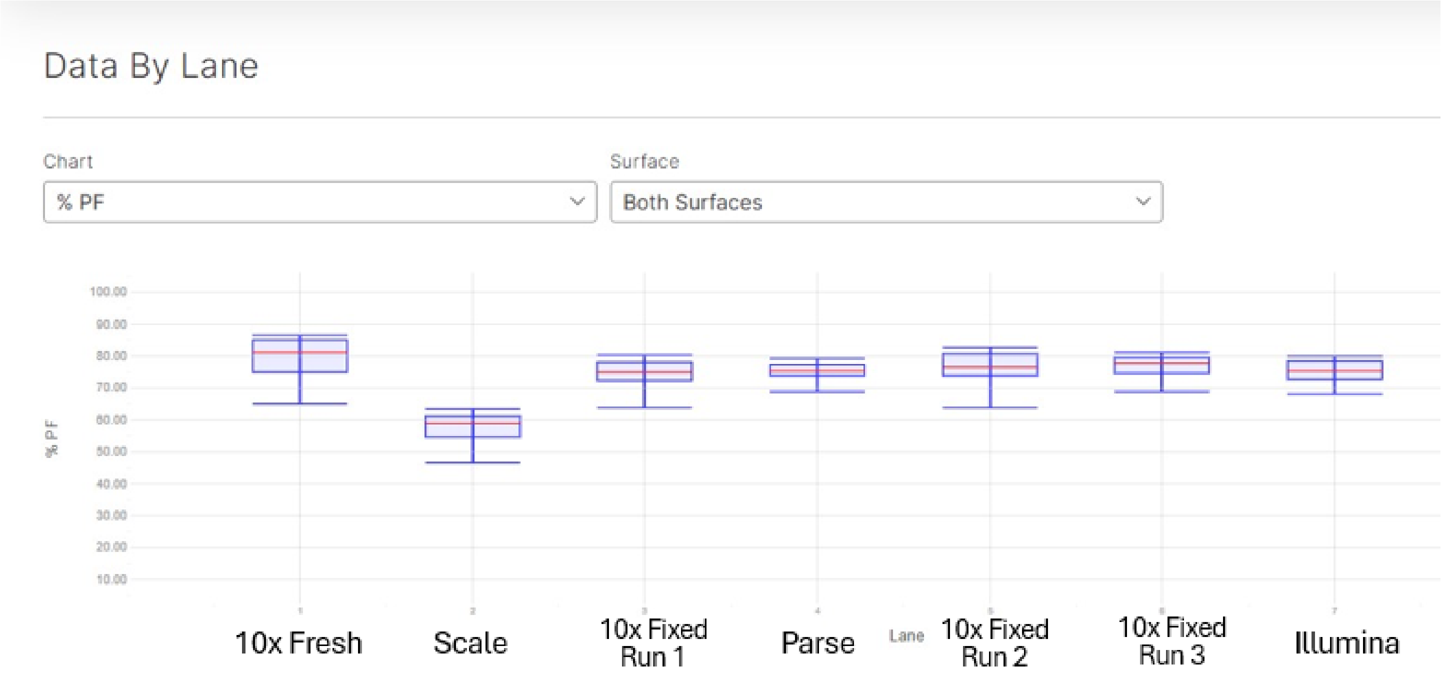


B

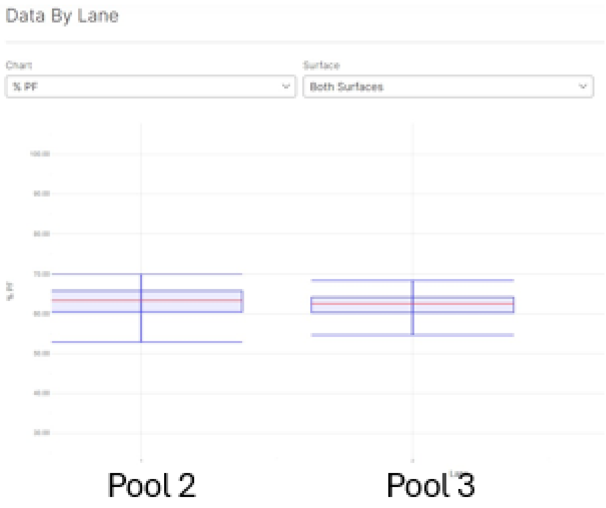


Supplemental Figure 9A – Pass/Fail for sequencing reads, divided by lane. Each lane is designated for a different technology. 10x fixed has three runs of data due to repeated clogging at GEM generation step.

Supplemental Figure 9B – Pass/Fail for Scale re-sequencing reads, divided by lane. Each lane is designated for a different pool ofScale's final distribution plate. Pool 2 is columns 5-8, Pool 3 is columns 9-12.

**Supplemental Figure 10 - Pseudobulk Principal Component Analysis (PCA)**

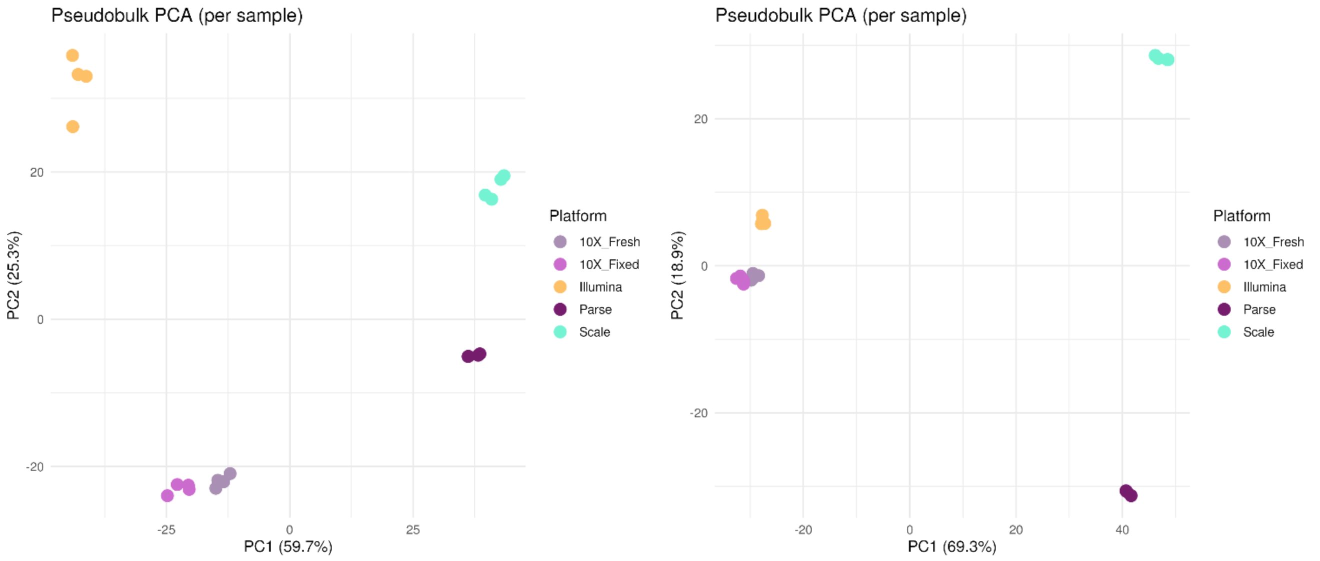


Agnostic Platform

Supplemental Figure 10 - Pseudobulk Principal Component Analysis (PCA), demonstrating high level of similarity between replicates.

**Supplemental Figure 11 - Pseudobulk sample-to-sample Pearson correlation**

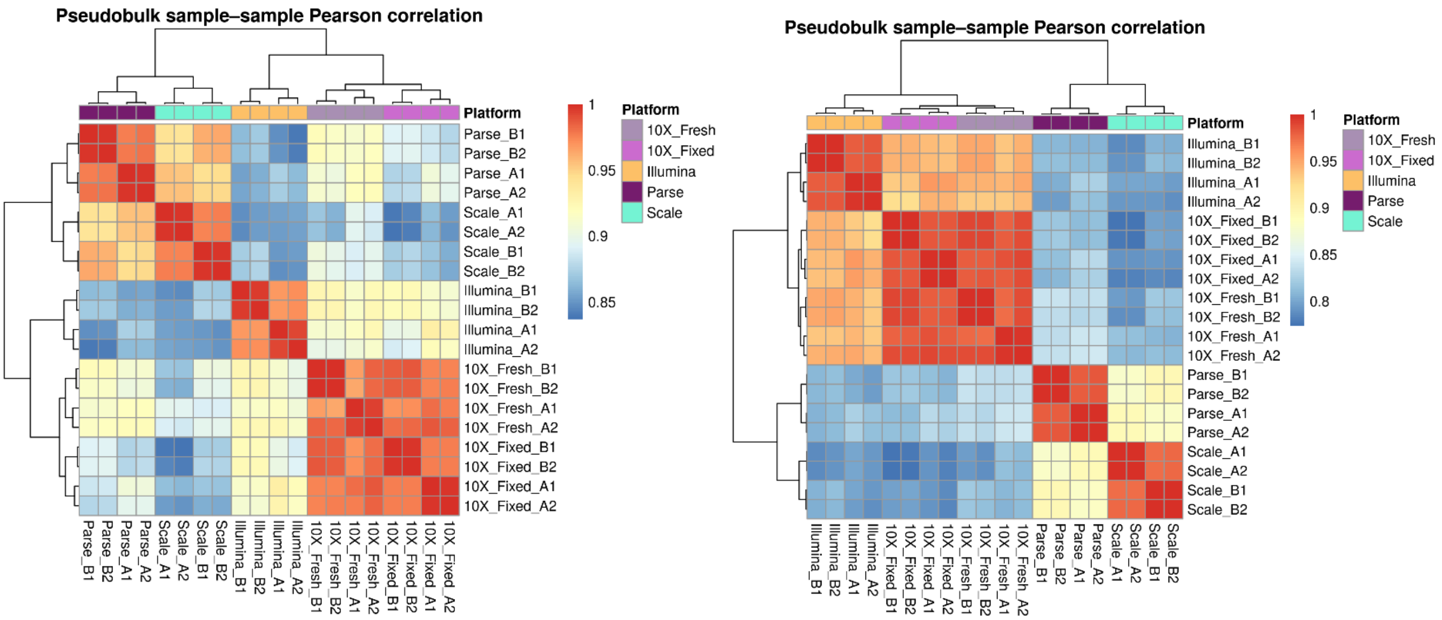


Agnostic                                        Platform

Supplemental Figure 11 - Pseudobulk sample-to-sample Pearson correlation, demonstrating high level of similarity between replicates.

**Supplemental Figure 12 – Unsupervised Clustering**

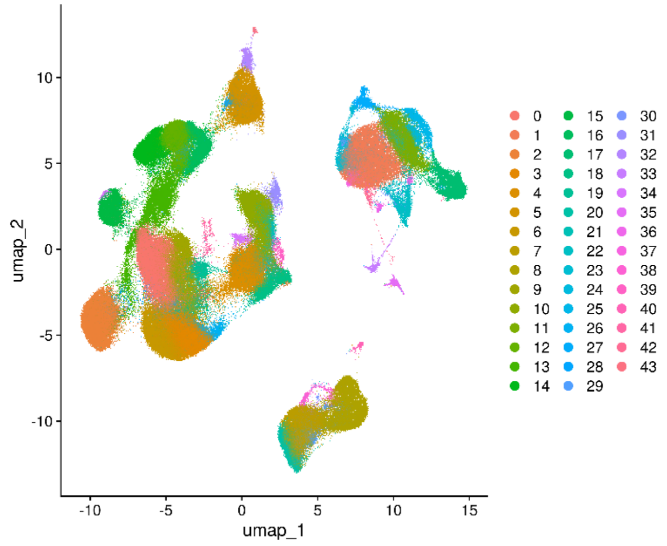


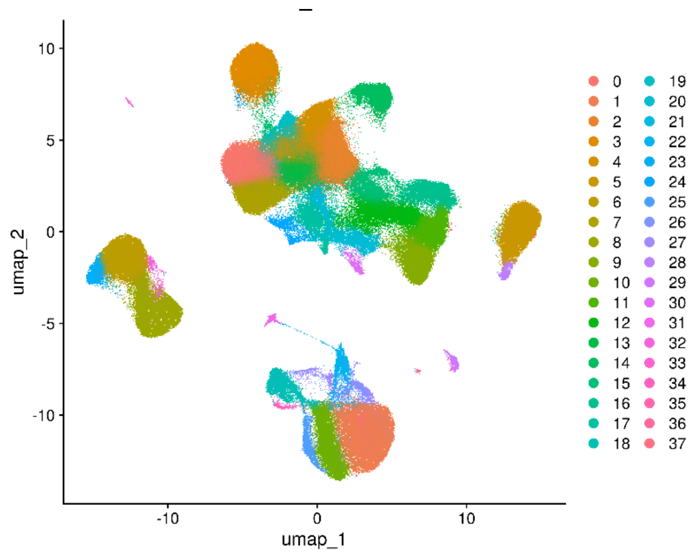


Agnostic

Platform

Supplemental Figure 12 – Unsupervised clustering of cell types, performed in Seurat. Color-coded by unlabelled cell type cluster.

#### Supplemental Figure 13 – Broad-level Cell Type UMAP

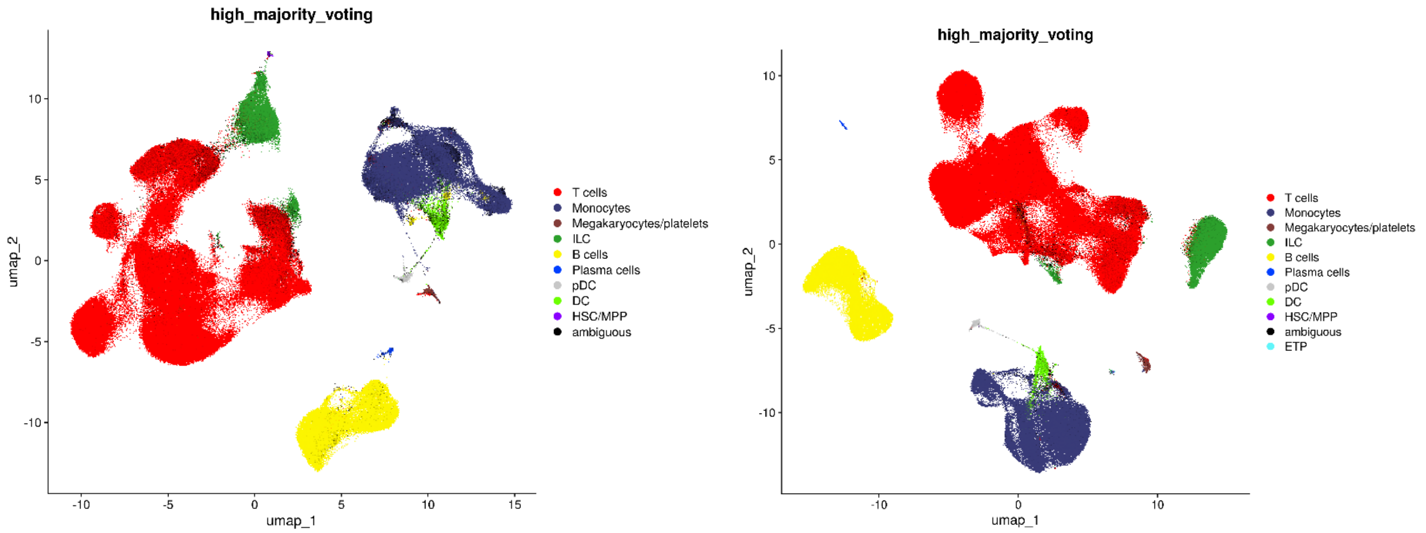


Agnostic Platform

Supplemental Figure 13 – Broad-level cell type classification, plotted as Uniform Manifold Approximation and Projection (UMAP). Demonstrates high-level cell types identified by clustering similar cell types in Seurat.

**Supplemental Figure 14 – Cell Subtype Classification UMAP**

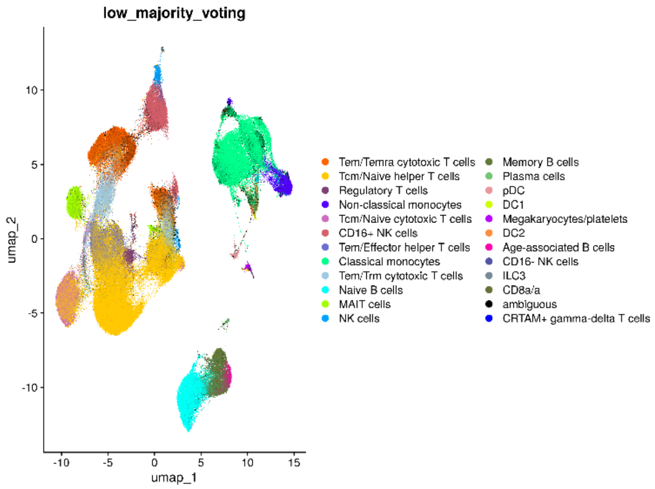


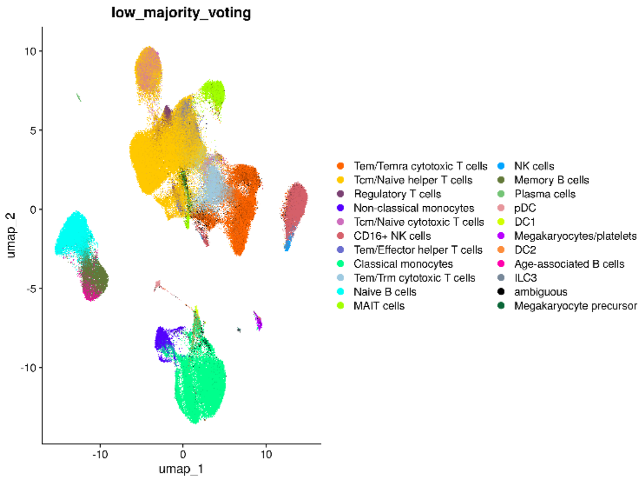


Agnostic Platform

Supplemental Figure 14 – Cell subtype classification, plotted as Uniform Manifold Approximation and Projection (UMAP). Demonstrates cell subtypes identified by clustering similar cell types in Seurat.
